# Macrophages target PVR/CD155 on colorectal cancer cells via REVERBα

**DOI:** 10.64898/2026.01.29.702476

**Authors:** Dominik Kato, Yu Han, Laura Helm, Frank Herweck, Natalia Schichta, Lennart Malz, Katharina Herrmann, Mohammad S. Farooq, Veronika Hauber, Tobias Gutting, Johannes Betge, Carsten Sticht, Timo Gaiser, Adelheid Cerwenka, Matthias P. Ebert, Elke Burgermeister

## Abstract

**Background&Aims:** Patients with microsatellite-instable (MSI) colorectal cancer (CRC) benefit from immune checkpoint therapy. For patients with microsatellite-stable (MSS) non-MSI tumors targeting alternative immune checkpoints, such as the *Poliovirus* receptor (PVR/CD155), may extend response to checkpoint inhibitors and thereby improve outcomes. The drugable transcription factor REVERBα (*NR1D1*) is a master repressor of macrophage function. We hypothesized that regulation of PVR allows elimination of tumor cells by macrophage-directed precision therapy.

**Methods:** Human CRC cell lines (MSS+: HT29, SW480; MSI+: HCT116), patient-derived organoids (MSS+ PDOs) and tissues were assessed by PCR, Western blot, immunohistochemistry and flow cytometry. 3D co-cultures of CRC cells with macrophages derived from THP1 monocytic leukemia cells or peripheral blood of healthy donors were analysed by microscopy and viability assays. Functional perturbation of PVR and REVERBα was achieved by CRISPR/Cas9-sgRNA gene modification, synthetic ligands or blocking antibodies (Abs).

**Results:** REVERBα bound a cognate DNA-element in the -1 kb human *PVR* gene promoter, and its agonist (SR9009) super-repressed, whereas its antagonist (SR8278) de-repressed transcription of *PVR* mRNA in macrophages. Macrophages with CRISPR-modified REVERBα were unresponsive to ligand, devoid of PVR protein, showed more phagocytosis, tumor cell efferocytosis and expression of genes related to host immunity (*PDL1*, *TLR4 e.a.*) than clones with the wild-type receptor. Macrophages lowered the viability of tumor cells, potentiated by PVR blocking Ab or *PVR* knock-out in tumor cells. Consistently, REVERBα antagonist augmented tumor cell death in co-cultures with macrophages and PVR blocking Ab.

**Conclusion:** Co-addressing the checkpoint axis REVERBα-PVR may represent a novel intervention strategy for patients with MSS+ CRC.

## Introduction

Gastrointestinal cancers including colorectal cancer (CRC) are classified into subtypes with distinct genetic alterations ^1^ and tissue microenvironments (TME) ^2^. Albeit, translation of molecular profiles to the clinic for individualized treatments and response prediction remains challenging ^3^. Only microsatellite instable (MSI) tumors (<10%) with high mutational burden, neo-antigen load and immune cell infiltration respond to antibodies (Abs) against T cell checkpoints (e.g. PD1, PDL1, CTLA4) ^2^. Thus, increasing the eligibility rates of MSS (non-MSI) patients is of medical need. This may demand inclusion of additional cell types of the TME for intervention, specifically from the innate immune system.

*Poliovirus* receptor (CD155/PVR) (and its paralogue CD112/PVRL2) belong to the nectin/like family of cell adhesion molecules ^4^ implicated in tissue homeostasis and anti-viral immunity. While forming PVR-Nectin/like *cis*-homodimers and *trans*-heterodimers in normal tissues, PVR up-regulation on tumor cells (including CRC) and macrophages enforces binding to inhibitory checkpoints such as the “T-Cell-Immunoreceptor-with-Ig-and-ITIM-domains” (TIGIT) and the CD112 receptor (PVRIG) on natural killer (NK) and T cells ^5^. Steric competition with activatory ligands such as DNAM1/CD226 or CD96/Tactile, among others, further contributes to the complexity of signaling outcomes. Nectins are transcriptionally induced by oncogenic driver pathways (e.g. WNT, RAS) in tumors, and by xenobiotic sensors (e.g. AHR) in macrophages.

These events mitigate immune cell expansion, differentiation and effector functions, e.g. phagocytosis, chemokine and cytokine secretion or cytotoxicity. In patients, gain of the PVR-TIGIT axis correlates with poor prognosis and metastasis ^6^. Accordingly, therapeutic Abs have entered clinical trials that disrupt mutual pairing of PVR with its counter-ligand TIGIT, thereby restoring immune effector activities ^5^.

REVERBα/β (*NR1D1/2*) are two members of the nuclear hormone receptor superfamily targetable by synthetic and physiological ligands ^7^. The latter comprise proto-porphyrins and oxidized forms of iron, including Fe^2+^ heme or Fe^3+^ hemin ^8^ . REVERBs lack a classical transactivation domain and bind as monomers to “retinoic acid receptor-related orphan receptor” (ROR)-responsive DNA-elements (ROREs) in regulatory regions (e.g. promoters, enhancers) of target genes (e.g. *IL6*).

Overall, they function as constitutive repressors of transcription by recruitment of nuclear receptor co-repressors (e.g. NCOR1) and epigenetic factors (e.g. HDAC3) ^9^. The physiological roles of REVERBs range from the regulation of circadian rhythms to metabolic diseases (e.g. atherosclerosis), cancer and immunity. Thereby, these receptors constitute promising targets within the hemorrhagic and hypoxic TME of CRC, enriched with iron-loaded macrophages.

We therefore explored the hypothesis that elimination of MSS+ CRC cells by macrophages is increased by inhibition of PVR expression and/or function. This intervention shall be achieved by pharmacological and/or genetic modulation of REVERBα in combination with PVR blocking Abs. To test this concept, we employed a translational program studying molecular aspects of *PVR* regulation by REVERBα in cell lines, PDOs and patients’ tissues.

## Materials and Methods

### Reagents

Chemicals were purchased from Merck or Sigma (Darmstadt, Germany) if not mentioned otherwise. Antibodies (Abs) are listed in **Table S1**. Recombinant human growth factors and cytokines were from Preprotech (Hamburg, Germany).

### DNA-constructs

pSpCas9(BB)-2A-Puro (PX459) vector was purchased from Addgene (#62988, Teddington, UK), sgRNAs were queried using E-CRISP target site identification tool from the German Cancer Research Center ^10^ (DKFZ, Heidelberg, Germany) corresponding to the following sequences in the human *NR1D1/REVERBA* (NM_021724.5) and *CD155/PVR* (NM_006505.5) genes (**Table S2**).

Synthetic sgRNA oligonucleotides (Eurofins Genomics Germany GmbH, Ebersberg, Germany) were annealed and inserted into the backbone vector generating recombinant plasmids pX459-sg*REVERBA* and pX459-sg*PVR* using NEB® Golden Gate Assembly Kit as specified by the manufacturer (New England Biolabs, Ipswich, MA). Genomic DNA encoding ∼1 kb from the human *PVR* (NM_006505.5) promoter was PCR-amplified (**Table S2**) and inserted into the KpnI/HindIII site of pGL3-basic luciferase (luc) reporter plasmid (Promega, Walldorf, Germany). *BMAL1*-luc was described before ^11^.

### Patients

The study followed the principles of the Declaration of Helsinki and was approved by the Medical Ethics Committee II of the Medical Faculty Mannheim, Heidelberg University (2014-633N-MA; 2016-607N-MA). All patients provided written informed consent prior to biopsy or peripheral blood collection. Study criteria and generation of PDOs have been detailed previously ^12^. From this prospective pseudonymized database combining clinical parameters and molecular tumor characteristics, four patients with MSS+ CRC (n=4) were selected for the present work (**Table S3**). Tissue microarrays (TMAs) from patients with CRC (Co484b) were purchased from TissueArray.Com LLC (US Biomax, Derwood, MD) together with the corresponding clinical information.

### Organoid culture

As detailed in ^12^, PDOs were grown in MatriGel™ (Corning, Thermofisher) in Advanced DMEM/F12 (Thermofisher) medium supplemented with 100 U/ml penicillin/streptomycin, 1 % (*v/v*) GlutaMAX and 1 % (*v/v*) HEPES (pH 7.2-7.5), 100 ng/ml noggin (Preprotech), B27 (Thermofisher), 1.25 mM n-acetyl cysteine, 10 mM nicotinamide, 50 ng/ml human epidermal growth factor (EGF), 10 nM gastrin (both from Peprotech), 500 nM A83-01 (Biocat, Heidelberg, Germany), 10 nM prostaglandin E_2_ (Santa Cruz Bio., CA), 10 µM Y27632 (Selleckchem, Houston, TX) and 100 mg/ml Primocin® (Invivogen, Toulouse, France), herewith defined as complete medium (“ENA”). PDOs were kept in a humidified incubator at 37 °C, 5% CO_2_, were passaged once a week, and medium was refreshed every second day.

### Cell lines

Human cell lines including embryonic kidney cells transformed by large T-antigen from Simian Virus-40 (HEK239T), monocytic leukemia (THP1), MSS+ (SW480, HT29) and MSI+ (HCT116) colorectal adenocarcinoma were obtained from the American Type Culture Collection (ATCC, Manassas, VA) and cultivated in complete Roswell Park Memorial Institute (RPMI) 1640 medium or Dulbecco’s Modified Eagle’s Medium (DMEM) according to the guidelines of the distributors. Basal media, herewith termed “complete” media, were supplemented with 10 % (*v/v*) fetal calf serum, 2 mM L-glutamine and 100 U/ml penicillin/streptomycin (all from Thermofisher).

### Transfection and generation of stable clones

Adherent, human CRC cell lines were transfected with empty vector (EV, pX469), pX459-sg*REVERBA* or -sg*PVR*, respectively, using Turbofect™ Transfection Reagent (Thermofisher) according to the manufacturer’s protocol. For selection of stable clones, transfectants were harvested 2 days after transfection and seeded in 96-well plates for limiting dilution cloning (∼1 cell per well) in complete medium supplemented with puromycin (1-10 µg/ml). THP1 monocytes (Mo/s) were subjected to electroporation (Neon™ Transfection System Kit, Thermofisher) as suspension culture, followed by clonal selection as above. Luciferase reporter plasmids were transiently transfected for 48 h, and total cell lysates subjected to Steady-Glo® Assay (Promega) using a multi-plate fluorescence reader (Tecan, Crailsheim, Germany).

### Cultivation of monocytic leukemia cell line

THP1 monocytes (Mo/s) were suspended in complete RPMI1640 medium and distributed to 6-well plates at a density of 5 x 10^5^ cells/ml. Phorbol 12-myristate 13-acetate (PMA) was added for 48 h (8 nM) to induce differentiation. Adherent M0 macrophages were washed once with PBS and incubated for additional 48 h at 37 °C, 5 % CO_2_ in complete RPMI1640 medium supplemented with cytokines (all from Peprotech) inducing polarization with either a combination of IFNγ (20 ng/ml) and LPS (*E. coli* O111:B4, 10 pg/ml) or IL4 and IL3 (20 ng/ml, each), respectively, herewith defined as “inflammatory” M1(IFNγ/LPS) and “regulatory” M2(IL4/IL13) macrophages according to current nomenclatures ^13^ ^14^.

### Isolation and cultivation of primary human monocytes

Peripheral blood mononuclear cells (PBMCs) were isolated by Ficoll® Paque Plus (GE Healthcare, Chicago, IL) centrifugation from fresh whole blood of healthy donors (German Red Cross DRK, Blood Donation Center, Mannheim, Germany) following standard procedures of the manufacturer^15^. Mononuclear cells were resuspended in complete RPMI1640 medium supplemented with 1 % (*v/v*) GlutaMAX and 1 % (*v/v*) HEPES (pH 7.2-7.5) (all from Thermofisher), seeded into 6-well plates at a density of ≤1x10^7^ per well and incubated at 37 °C, 5 % CO_2_ for 24 h to allow monocytes to adhere. Thereafter, non-adherent suspension cells (containing lymphocytes) were harvested, adherent cells washed with PBS, and monocytes were maintained for 5 days in complete medium in presence of 100 ng/ml M-CSF (Preprotech), respectively, followed by a 48 h incubation with or without cytokines, as described for the monocytic leukemia cell line above. Alternatively, primary monocytes were enriched by magnetic-activated cell sorting (MACS) following the recommendations of the manufacturer (Pan Monocyte Isolation Kit™, Miltenyi, Bergisch Gladbach, Germany).

### Live cell imaging

Phagocytosis was quantified in single cultures (SC) of THP1-derived macrophages by uptake of fluorescent beads following the guidelines of the manufacturer (Cell Meter™ Fluorimetric Assay Kit, AAT Bioquest, Biomol, Hamburg, Germany). For detection of efferocytosis in contact-dependent co-cultures, HT29 cells were labelled (marked by asterisk *) with pHrodo® (λ ex = 532 nm) dye as detailed by the manufacturer (Molecular Probes, Thermofisher). Subconfluent monolayers of THP1-derived or primary macrophages were prepared in 6-well (for RNA/Protein) or 48-well (for MTT) plates before the experiment. Macrophages were then overlaid with a 3:1 ratio of HT29 cells and incubated for additional 5 days in mixed media at 37 °C, 5 % CO_2_. Macrophages were counted by autofluorescence (488 nm), surviving cancer cells in the red channel (λ em = 655 nm). Alternatively, uptake of tumor cells, which had been pre-labelled with carboxyfluorescein succinimidyl ester (CSFE, CellTrace™, Thermofisher), by macrophages was quantified after 24 h using flow cytometry (FC). In addition, cell death was determined by annexin (ANX, apoptosis) and propidium iodide (PI, necrosis) following the manufacturer’s procedure (Annexin-V-FLUOS Kit, Roche). Images were acquired at an original magnification of 100-200x using a digital camera-connected AXIO Observer.Z1 - ApoTome.2. fluorescence microscope and ZEN software (Zeiss, Jena, Germany). Finally, co-cultures were harvested as mixed samples for RNA or protein extraction without further separation.

### Colorimetric viability assay

Colorimetric *in-situ* cell viability assay based on 3-(4,5-dimethylthiazol-2-yl)-2,5-diphenyltetrazolium bromide (MTT) was conducted according to the manufacturer’s protocol (Roche).

### Organoid cocultures

For contact-dependent 3D co-cultures, PBMC-derived macrophages were prepared as above. The next day, intact PDO spheroids were mixed into ice-cold MatriGel™ at an 50:1 effector:target ratio (in 30 µl MatriGel™ per 1.000 PDO rings/50.000 macrophages) and seeded into pre-cooled 48-well plates. In brief, 3 drops per well were gently pipetted on the bottom of the ice-cold well, and plates were kept on ice for 10 min to let gravity to evenly disperse the PDO mixture on top of the adherent macrophage monolayer. Plates were then left to solidify (up-side-up) for 1 h at 37 °C in the incubator before addition of 500 µl medium [50 % ENA+Y / 50 % RPMI1640 containing 10 % FCS, 1 % Pen./Strep., 1 % glutamine, 1 % HEPES, 1 % GlutaMAX, all (*v/v*)] supplemented with treatments as indicated in the legends to figures. Co-cultures were kept for 5 days before subjection to assays as described in legends to figures.

### Immunofluorescence microscopy (IF)

*In situ* whole mount staining of PDOs was done after fixation as detailed in ^16^. In brief, PDOs were seeded in 8-well chamber slides (10 µl in 3 MatriGel™ drops per well), followed by fixation in 4 % (*w/v*) paraformaldehyde (PFA) buffered in PBS for 30 min at room temperature and quenching of autofluorescence using 50 mM NH_4_Cl for 30 min. PDOs were permeabilized by 0.5 % (*v/v)* Triton X-100 in PBS for 30 min, and subsequently blocked with 5 % (*w/v*) bovine serum albumin (BSA) in PBS for 60 min and stained as below for paraffin sections using 5 % (*w/v*) BSA/PBS.

Formalin-fixation and agarose/paraffin-embedding (FFPE) of PDOs has been described before ^17^, employing MicroTissues™ 3D Petri Dish micromolds (Sigma). Sections (3-5µm) were cut with a microtome (Leica RM 2145) and transferred to glass slides (Superfrost™, Thermofisher).

Hematoxylin and eosin (H&E) stainings were performed using automated staining devices. Imaging of FFPE-sections from PDOs was done as described for tissues ^18^. In brief, dewaxed and rehydrated slides were subjected to antigen retrieval as recommended by the manufacturer (Vectorlabs, Burlingame, CA). After 1 h blocking with 100 % (*v/v*) FCS at room temperature, primary Ab [in 0.3-0.5 % Triton X-100, 10 % FCS / PBS, all (*v/v*)] was added overnight at 4 °C, followed by an 1 h incubation with secondary Ab (in 10 % FCS/PBS (*v/v)*) and DAPI (50 ng/ml) for 10 min in the dark at RT. Mounting medium (Dako, Hamburg, Germany) was used to cover the slide, and images were acquired in triple color mode as before. Staining of adherent cells followed a previous protocol ^18^.

Fluorescence signals (n>20 cells or nuclei per field, n=5 fields per image) from Abs, phalloidin dye and 4’,6-diamidino-2-phenylindol (DAPI) were manually counted with Image J (imagej.nih.gov/ij). Images were acquired using Zeiss Light Microscope Axio Observer.Z1 (invers), Leica Light Microscope DMRE (upright) or Leica Confocal Microscope (inverted, TCS SP5 MP (Multiphoton). **Immunohistochemistry (IHC)**

Ab staining on sections from FFPE tissues or PDOs was done as before ^18^. Antigen retrieval was performed by heating in antigen unmasking solution (Vectorlabs, Burlingame, CA) and blocking with H_2_O_2_. Abs were diluted as given in **Table S1**, and staining was processed as recommended by the Vectastain ABC kit (Vectorlabs). The substrate 3,3′-diamino benzidine (DAB) (Vectorlabs) served as a detection reagent followed by counterstaining with hematoxylin. Frequency and intensity of staining positivity was determined in epithelial and lamina propria (stroma) cells. The staining scores were defined as: 0+ = negative (0-25%), 1+ = weak (25-50%), 2+ = moderate (50-75%), 3+ = strong (75–100%). Percentages were calculated as positive cells divided by the total number of cells within the field. Signals were quantified observer-blinded at a standard bright-field microscope using Image J (imagej.nih.gov/ij) (n >20 signals per field; n=5 fields per image). The field area was defined by the morphology (e.g. crypt-villus unit, PDO spheroids or macrophage clusters) and measured in square mm (mm^2^).

### Flow cytometry (FC)

Procedures followed previous protocols ^17^. In brief, for detection of surface markers, PDOs were collected from 6-well plates in ice-cold DPBS (supplemented with 2 % (*v/v*) FCS) by scrapping, and MatriGel™ drops were dissociated by repetitive pipetting. PDOs were pelleted by centrifugation (400 x g, 10 min, 4 °C) followed by removal of the supernatant. Then, PDOs were resuspended in 1 ml of StemPro Accutase® (Thermofisher) per 6-well plate and incubated for 35 min under continuous shaking at 400 rpm at 37°C in a Thermomixer. After confirming dissociation into single cells by microscopy, cells were washed with ice-cold DPBS / 2 % (*v/v*) FCS, filtered through a 35µm cell-strainer and recovered in PBS or growth media for flow cytometry (FC). Cell lines were harvested accordingly. Single cells were then incubated for 10 min at 4°C with 200 µl DPBS, 2 % (*v/v*) FCS and FcR human blocking reagent (Miltenyi, Bergisch Gladbach, Germany) prior to extracellular staining. Extracellular staining was performed with cells or dissociated organoids in ice-cold PBS containing 2 % (*v/v*) FCS, 2 mM EDTA and 0.05 % (*w/v*) NaN_3_ with fluorochrome-labelled Abs (as detailed in **Table S1**) for 30 min in the dark at 4 °C. Dead cells were excluded upon labelling with 7-amino-actinomycin D (7-AAD, BioLegend). Signals were acquired with Fortessa or FACSCantoII® devices (BD Biosciences, San Jose, CA). Analysis was performed using FACSDiva® (version 8.0.1) and FlowJo (version 10) (all from BD Biosciences). For quantification of cell death in dissociated co-cultures, epithelial/tumor cells were labelled with Abs against EpCAM and macrophages with CD11b (**Table S1**). The percentages of early and late apoptotic *vs*. necrotic cells was counted in quadrant plots using the dyes Annexin V-FITC (Roche, Merck) and SYTOX™ Blue Dead Cell Stain (Thermofisher), respectively.

### Protein extraction and Western blot

Methods were conducted as described previously ^18^. Abs are listed in **Table S1**. Briefly, total cell lysates were prepared from cells using lysis buffer with detergent (50 mM Tris-HCl, pH 7.4, 1 % (*w/v*) SDS, 1 mM Na_3_VO_4_, 1 mM DTT, Protease Inhibitor Complete®, Roche) and subjected to SDS-PAGE and immunoblot detection by chemoluminescence. Membranes were imaged using the Fusion Solo S/L CCD imaging system (Vilber Lourmat Deutschland GmbH, Germany).

### Nucleic acid isolation, chromatin immunoprecipitation (ChIP), reverse transcription (RT) quantitative PCR (RT-qPCR)

Samples were processed and analyzed as outlined before ^19^, and methods were performed on genomic DNA or total RNA. Oligonucleotides are listed in **Table S2**.

### Software and statistics

Results are displayed as means ± S.E. from at least 3 independent experiments from different cell passages or patients. Optical densities (O.D.) of bands in gels from Western blots and PCRs were measured using automated imaging devices and quantified with Image J (imagej.nih.gov/ij). Data were normalized to house-keeping genes or proteins as indicated in the legends to figures and calculated as -fold or % compared to control. Statistical analysis was conducted according to the guidelines on non-*vs.* parametric data distribution and appropriate tests for 1-to-multiple-groups provided by Graphpad Prism (version 4.0, La Jolla, CA). All tests were unpaired and two-sided if not stated otherwise. P-values < 0.05 were considered significant (*). Amino acid (aa) and gene sequence alignments (**Table S4-S6**) and computational modelling of protein 3D structures were performed using BLAST (https://blast.ncbi.nlm.nih.gov) and PHYRE2 Protein Fold Recognition Server (https://www.sbg.bio.ic.ac.uk). Prediction of transcription factor binding sites was done by using Alibaba (Transfac) version 2.1 (http://gene-regulation.com/pub/programs/alibaba2/).

## Results

### REVERBα binds to and regulates the human *PVR* gene

We first explored, if REVERBα can be targeted by synthetic ligands to change expression of cell surface immune checkpoints, as exemplified here by *PVR*. To this end, we examined if macrophages *per se* express PVR (**Fig.1A).** This conjecture was confirmed by data sets from NCBI GEOprofiles^®^ on the expression of *PVR* mRNA in human myeloid subsets (“Monocyte differentiation to macrophage and subsequent polarization” HG-U133B/A; *Homo sapiens* n=3 replicate per cell type; ID32429947).

**Fig. 1.**
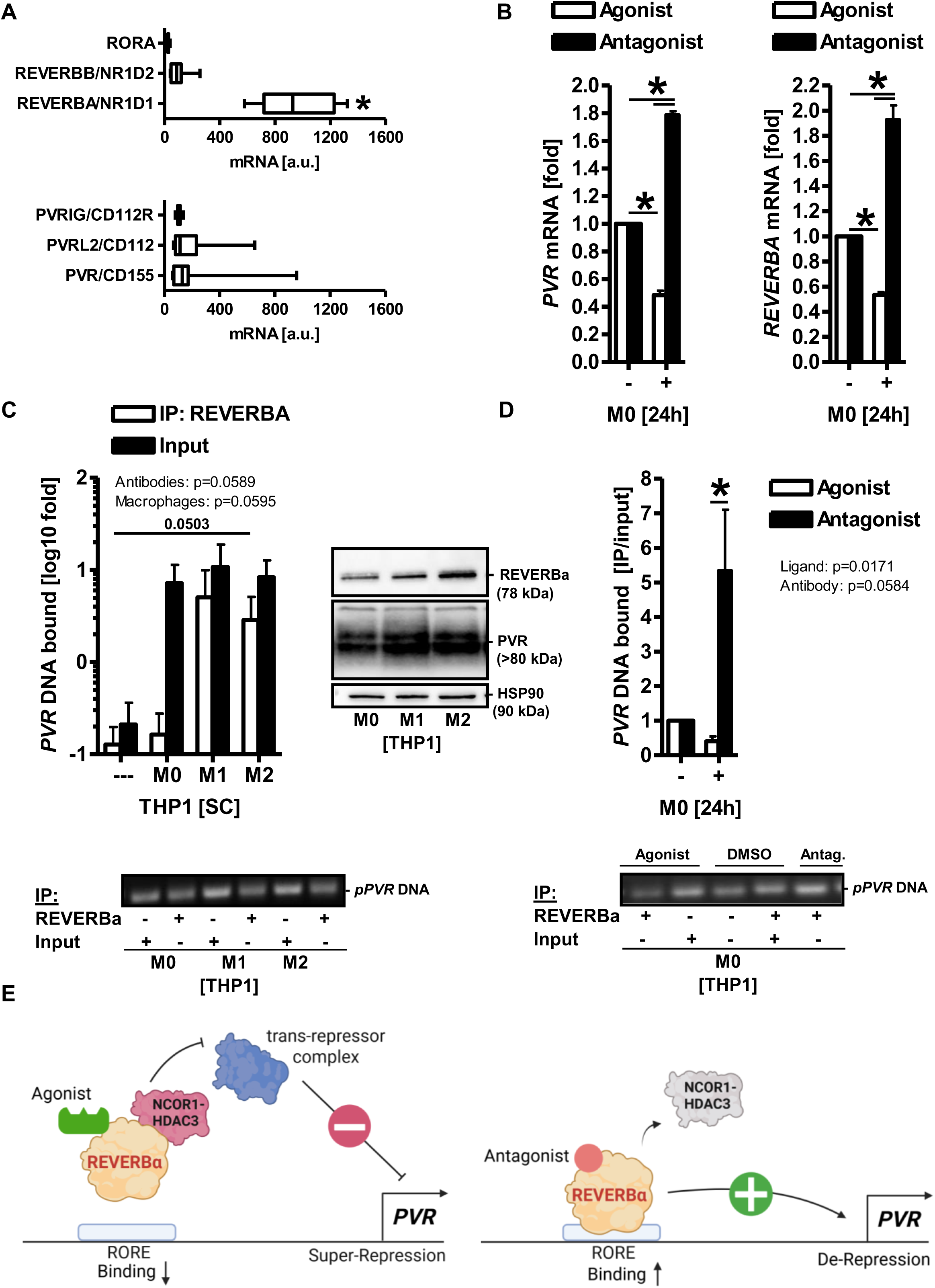
REVERBα binds to and regulates the human *PVR* gene. **A**, Association of *NR1D1* and *PVR* mRNA with myeloid subsets. Data were retrieved as log2 values from NCBI GEOprofiles^®^ and presented as means ± S.E. (*p<0.05, Friedman test with Dunn post-test, subgroup analysis with Wilcoxon-matched-pairs test). “Monocyte differentiation to macrophage and subsequent polarization” (HG-U133B/A; *Homo sapiens* n=3 replicate per cell type; ID32429947). **B,** REVERBα agonism leads to super-repression of *PVR* mRNA. THP1 monocytes were differentiated to macrophages and then treated with vehicle controls (DMSO), REVERBα agonist (SR9009, 1 μM) or antagonist (SR8278, 0.67 μM) for additional 24 h. Ct-values from RT-qPCRs were normalized to *B2M* and calculated as -fold ± S.E. (*p<0.05 *vs.* vehicle, 2way-ANOVA with Tukeýs post-test, subgroup analysis with Wilcoxon signed rank test, n=3 replicates per macrophage type). Additional genes are presented in (**S1**). **C-D,** REVERBα binds the predicted RORE-like motif 5’-*gagac **AGG TC**T-3’* (NG_008781.2: 4556 to 4561 bp) in the *PVR* promotor. THP1 macrophages were treated as in A. ChIP was performed using REVERBα Ab for pull-down followed by extraction of precipitated genomic DNA and genomic PCR. Quantitative analyses (top) and corresponding ethidium bromide-stained agarose gels (bottom) showcasing (**C**) basal and (**D**) ligand-depending binding of REVERBα to the RORE DNA. Ct-values from qPCRs were normalized to total chromatin input of the *B2M* gene and calculated as -fold ± S.E. (*p<0.05 *vs.* no-Ab control or vehicle, 2way-ANOVA with Tukeýs post-test, subgroup analysis with One-Sample t-test, n=3 replicates per macrophage type). Insert: Macrophages express REVERBα and PVR proteins. THP1-derived macrophages were subjected to total protein extraction. Representative images from Western blots. **E, Model of *PVR* gene regulation.** REVERBα binds a cognate RORE-like DNA-element in the -1 kb human *PVR* gene promoter in macrophages. Left panel: Agonist (SR9009) binding to REVERBα leads to recruitment of a “nuclear receptor co-repressor” complex comprising NCOR1 and HDAC3, followed enhanced repressor activity, reduced RORE binding and super-repression of the *PVR* promoter, presumably involving an additional, yet to be identified, trans-repressor complex; Right panel: Antagonist (SR8278) binding to REVERBα triggers detachment of NCOR1-HDAC3, followed by mitigation of its repressor activity, enhanced RORE binding and de-repression of the *PVR* promoter, suggesting REVERBα as an indirect positive modulator of *PVR* expression.

We next asked if, REVERBα ligands alter expression of *PVR* mRNA (**Fig.1B).** M1(LPS+/IFNγ+) and M2(IL4+/IL13+) polarised macrophage subsets were generated from PMA-differentiated, unpolarized, adherent M0 macrophages originating of the human THP1 monocytic leukemia cell line. Macrophages were then treated with vehicle controls (DMSO), REVERBα agonist (SR9009, 1 μM) or antagonist (SR8278, 0.67 μM) for 24 h, respectively. Quantitative analyses from RT-qPCRs demonstrated that REVERBα agonism super-repressed *PVR* mRNA, whereas REVERBα antagonism de-repressed it (*p<0.05 *vs.* vehicle, 2way-ANOVA with Tukeýs post-test, subgroup analysis with Wilcoxon signed rank test, n=3 replicates per macrophage type).

This result was consistent with the model of REVERBα acting as a constitutive transcriptional repressor at the DNA ^9^, and was further strengthened by the same reciprocal expression pattern exerted on its *bona fide* RORE at the human *BMAL1* promoter (**S1a**). Notably, similar patterns could also be observed with *VEGFA, PDL1* and *PDCD1* (*PD1*) mRNAs (**S1b**), both in THP1- and PBMC-derived macrophages, indicating that drugging the clock protein REVERBα in macrophages may be useful to alter immune checkpoint expression.

This effect could be reproduced on the tumor side (**S1c**). Again, REVERBα agonism led to super-repression of the *PVR* promoter in the human MSS+ HT29 CRC and HEK293T (non-cancer control) cell lines. Here, cells were transiently transfected with the pGL3 reporter plasmid harbouring the proximal -1 kb promoter of the human *PVR* gene, followed by an 48 h treatment with REVERBα ligands (as in A) and quantification of luciferase activity in total cell lysates (*p<0.05 *vs.* vehicle, 2way-ANOVA with Tukeýs post-test, subgroup analysis with Wilcoxon signed rank test, n=3 replicates per cell line), alluding at a more general principle of *PVR* regulation by REVERBα across cell types.

To detect the corresponding proteins, THP1 macrophages (**Fig.1C**) were subjected to extraction of total protein lysates. Western blots evinced a high *a priori* expression of PVR and REVERBα in macrophages. This finding led us to the hypothesis that REVERBα directly binds to DNA-elements in the human *PVR* gene promoter. To explore this, we conducted an *in-silico* search on the upstream regulatory sequence of the human *PVR* gene using the Alibaba (Transfac) engine ^20^, and discovered a RORE-like motif 5’-*gagac **AGG TC**T-3’* (NG_008781.2: 4556 to 4561 bp) in the -1 kb proximal *PVR* promotor **(Fig.1D, Table S4)**. To measure binding of REVERBα protein to this element, THP1 macrophages were treated as in A, and chromatin was extracted from whole cell lysates.

ChIP was then performed using REVERBα Ab for pull-down followed by enrichment of precipitated genomic DNA. For quantification of IP-ed DNA, the Ct-values from genomic qPCRs were normalized to the total chromatin input of the *B2M* gene. REVERBα binding to the DNA-element compared to the “no-Ab” (IgG) control was evident in all macrophage types and augmented by REVERBα antagonist compared to the agonist (*p<0.05 *vs.* no-Ab control or vehicle, 2way-ANOVA with Tukeýs post-test, subgroup analysis with One-Sample t-test, n=3 replicates per macrophage type).

These data were consistent with a mode of action, where REVERBα acts as an indirect positive modulator of *PVR* gene expression (see model in **Fig.1E**). In sum, these data indicated that REVERBα binds to and regulates the *PVR* gene.

### REVERBα “loss-of-function” augments effector functions in THP1 macrophages

Based on this model (**Fig.1E**), we then intended to silence the constitutive expression of REVERBα in macrophages. To achieve this, THP1 monocytes were transfected with CRISPR/Cas9-*REVERBA* sgRNA knock-out (KO) plasmid or empty vector (EV) control by electroporation, followed by clonal selection and RNA extraction. Quantitative analysis of RT-qPCRs showed that the sgRNA clones expressed similar levels of total *REVERBA* mRNA compared to EV clones (n.s., Wilcoxon signed rank test, n≥3 replicates per clone). (**Fig.2A)**.

**Fig. 2.**
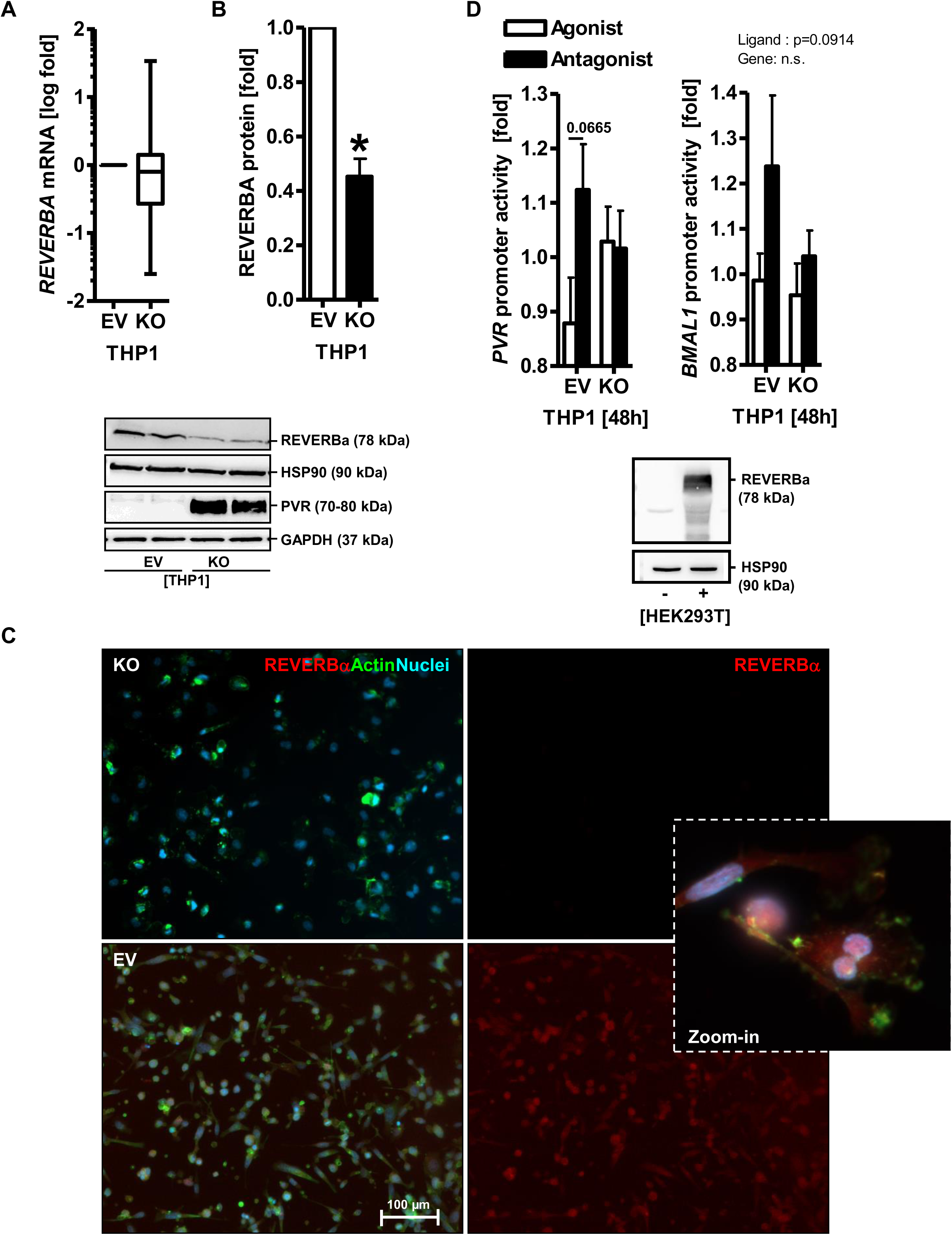
REVERBα loss-of-function in THP1 macrophages. **A**, LOF clones express *REVERBA (NR1D1)* mRNA. THP1 monocytes were transfected with CRISPR/Cas9-*REVERBA*sgRNA knock-out (KO) or empty vector (EV) plasmid by electroporation, followed by clonal selection and RNA extraction. Ct-values from RT-qPCRs normalized to *B2M* were calculated as -fold ± S.E. (n.s., Wilcoxon signed rank test, n≥3 replicates per clone). **B**, LOF clones express less REVERBα protein. Clonal cells from A were differentiated and polarised to macrophages and then subjected to extraction of total cell lysates. Non-cancer HEK293T cells were transiently transfected with REVERBα expression plasmid for 96 h. Representative images (bottom) and quantitative analyses (top) from Western blots using an Ab against the N-terminus (#14506-1-AP) of the FL REVERBα protein. O.D. values of bands in gels normalized to HSP90 are -fold ± S.E. (*p<0.05 *vs.* EV, 2way-ANOVA with Bonferroni post-test, subgroup analysis with Wilcoxon signed rank test, n=3 per replicates per clone). Note that loss of REVERBα correlated with loss of PVR protein. **C**, LOF clones express less nuclear REVERBα protein. Cells from B were fixed, permeabilized and stained for immunofluorescence (IF) microscopy (**Table S1)**. Color code: red = REVERBα; green = actin (phalloidin); blue = nuclei (DAPI). Representative fluorescence images are shown. Original magnification 100x; Scale bars = 100 µm. **D**, LOF clones lack ligand sensitivity towards RORE+ promoters. Cells from B were transiently transfected with pGL3 reporter plasmid harbouring the proximal -1 kb promoter of the human *PVR* gene and *BMAL1 (ARNTL)* as pos. control, followed by an 48 h treatment with vehicle controls (DMSO) or REVERBα ligands. Luciferase activity normalized to protein content is presented as - fold ± S.E. (*p<0.05 *vs.* vehicle, 2way-ANOVA with Tukeýs post-test, subgroup analysis with Mann Whitney test, n=3 replicates per clone).

Nonetheless, when clonal monocytic cells were differentiated to macrophages and subjected to extraction of total cell lysate, quantitative analyses from Western blots using an Ab against the N-terminus (#14506-1-AP) of the full-length (FL) REVERBα protein confirmed that the sgRNA clones expressed significantly less REVERBα protein compared to clones transfected with control sgRNAs (not shown) or EV (**Fig.2B)** (*p<0.05 *vs.* EV, 2way-ANOVA with Bonferroni post-test, subgroup analysis with Wilcoxon signed rank test, n=3 per replicates per clone).

Notably, REVERBα knock-down was accompanied by a reduction in *PVR* mRNA and protein expression, underscoring the model (in **Fig.1E)** that REVERBα is an indirect positive modulator of *PVR* gene expression.

To visualise the nuclear localisation of REVERBα, clonal cells were fixed, permeabilized and stained with REVERBα Ab for intracellular immunofluorescence (IF) microscopy (**Fig.2C)**. Therein, the KO cells had substantially less positivity for nuclear REVERBα protein compared with the EV control.

To further test, if the KO also confers a lack of functionality, clonal cells were transfected with pGL3 reporter plasmid harbouring the proximal -1 kb promoter of the human *PVR* gene and *BMAL1 (ARNTL)* as pos. control, followed by 48 h treatment with REVERBα ligands (**Fig.2D)**. Quantification of luciferase activity in total cell lysates (*p<0.05 *vs.* vehicle, 2way-ANOVA with Tukeýs post-test, subgroup analysis with Mann Whitney test, n=3 replicates per clone) demonstrated that the KO clones lack ligand sensitivity towards all promoter constructs that harboured a RORE-like DNA-element.

Sequence alignment and *in-silico* secondary structure prediction (**S2**) of the CRISPR/Cas9-modified REVERBα underscored the apparant “loss-of-function” (LOF) phenotype. The target site for the C-terminal sgRNA (1878-1897 bp) was localized in the C-terminal exon E6 (1744-1929 bp), the N-terminal sgRNA in the 5’-UTR upstream of the ATG start codon of the *NR1D1* mRNA (NM_021724.5) (**Table S5**).

Upon alignment of human wild-type (WT) full-length (FL, aa 1-614) (NP_068370.1; P20393_HUMAN) and the C-terminally truncated ΔCT (aa 1-468) proteins (**Table S6**), secondary structure prediction ^21^ with *Phyre2* suggested a disordered folding of the peptide region in the C-terminal ligand-binding domain (LBD) of the truncated REVERBα protein ΔCT (aa 132-468), whereas the WT protein (aa 434-611) gave full match [c3n00A: Confidence 100 / Identity 100 %] to the PDB X-ray structure of the LBD of human “*nuclear receptor subfamily 1 group D member* 1” (NR1D1).

Nonetheless, *NR1D1* locus sequencing and PCR genotyping of the C-terminal exon 6 did not reveal explicit alterations (e.g. truncations, indels) at the C-terminal sgRNA target site (unpublished observation). Consistent with the predicted disordered LBD, the LOF phenotype was hence restricted to the protein function, specifically on the ligand sensitivity, as shown by the luciferase reporter assays.

Having generated the stable clones, we tested the effect of REVERBα LOF on cognate macrophage effector functions such as phagocytosis (**Fig.3A).** To this end, clonal cells were differentiated and polarised to macrophages before exposure to Protonex™ Red 600-labelled latex beads for 4 h followed by incubation with CytoTrace™ Green for additional 30 min. Data from live imaging were the “phagocytotic index” calculated as the ratio of red bead counts divided by the number of green cells per field (p<0.1 *vs.* EV, 2way-ANOVA with Tukeýs post-test, subgroup analysis with Wilcoxon signed rank test, n=5 fields per sample, n=3 replicates per clone). Overall, the LOF clones trended to show enhanced phagocytosis compared to the EV control.

**Fig. 3.**
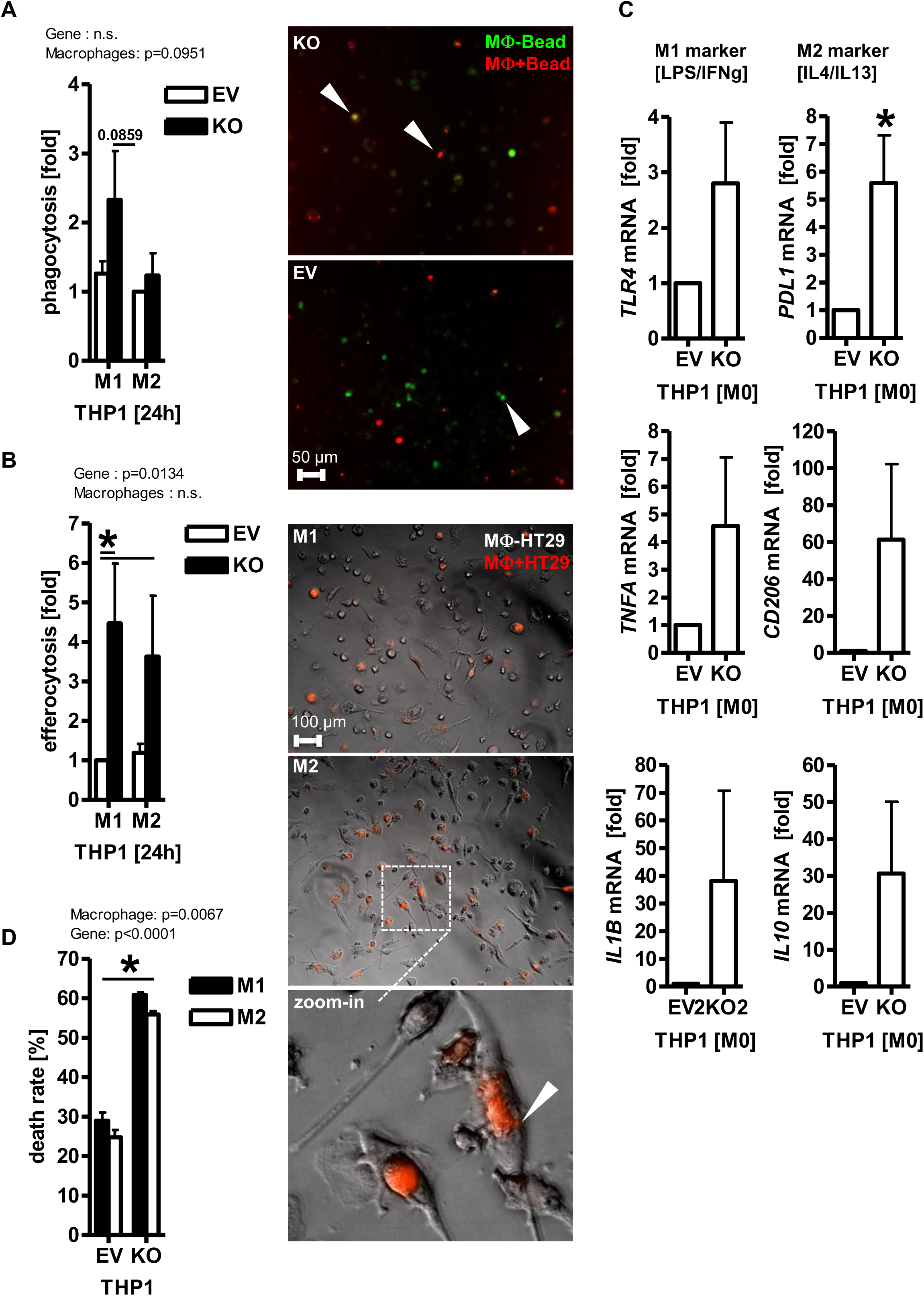
REVERBα loss-of-function enhances effector phenotypes of THP1 macrophages. **A**, LOF clones show enhanced phagocytosis. Clonal cells were differentiated and polarised to macrophages before exposure to Protonex™ Red 600-labelled latex beads for 4 h followed by incubation with CytoTrace™ Green for additional 30 min. Representative images (right) and quantitative analyses (left). Data from live imaging is the “phagocytotic index” calculated as the ratio of red bead counts divided by the number of green cells per field ± S.E. (p<0.1 *vs.* EV, 2way-ANOVA with Tukeýs post-test, subgroup analysis with Wilcoxon signed rank test, n=5 fields per sample, n=3 replicates per clone). Color code: red = beads; green = cells. Original magnification 100x; Scale bar = 50 µm. **B**, LOF clones show enhanced efferocytosis of CRC cells. Clonal cells as in A were exposed to CSFE-labelled HT29 cells for 24 h before analysis by flow cytometry (FC). Data are mean numbers of CSFE+ macrophages ± S.E. (*p<0.05 *vs.* EV, 2way-ANOVA with Tukeýs post-test, subgroup analysis with Wilcoxon signed rank test, n=3 replicates per clone). Insert: Efferocytosis imaging. THP1 macrophages were grown as in A before exposure to phRodo®-labelled HT29 cells for additional 48 h. Representative images capturing the ingested and remaining red-labelled HT29 cells. Color code: red = HT29; grey (bright field) = macrophages. Original magnification 100x; Scale bar = 100 µm. **C**, LOF clones show enhanced cytokine expression. Clonal cells from A were subjected to total RNA extraction. Ct-values from RT-qPCRs normalized to *B2M* were calculated as -fold ± S.E. (*p<0.05 *vs.* EV, Mann Whitney test, n=3 replicates per clone). **D**, LOF clones reduce overall viability of mixed co-cultures. Clonal cells as in A were overlaid with HT29 cells at an effector:target ratio of ∼3:1 and subsequent co-culture for 5 days. Adherent cells were subjected to colorimetric MTT viability assay. O.D. values were calculated as % of dead (= 100 - % surviving) cells ± S.E. in co-culture (CC) *vs.* the sum of the single cultures (ΣSC) (*p<0.05 *vs.* ΣSC, 2way-ANOVA with Bonferroni post-test, n=3 replicates per clone).

Similar results were obtained from efferocytosis assays (**Fig.3B)**. Here, clonal cells as in A were exposed to CSFE-labelled HT29 cells for 24 h before analysis by flow cytometry (FC). Mean numbers of CSFE+ macrophages with ingested tumor cells (*p<0.05 *vs.* EV, 2way-ANOVA with Tukeýs post-test, subgroup analysis with Wilcoxon signed rank test, n=3 replicates per clone) were increased in the LOF clones compared with the EV control.

Alternatively, THP1 macrophages were grown as in A before exposure to phRodo®-labelled HT29 cells for 48 h. Live cell imaging (**Fig.3B)** visualised the numbers of remaining, non-phagocytosed, red-labelled HT29 cells per field. Again, this *in vitro* surrogate assay for efferocytosis revealed an enhanced number of ingestion events in the LOF clones.

Consistent with a global de-repression program after REVERBα inhibition, a series of genes encoding for immune cell surface receptors or cytokines were elevated in the LOF clones (**Fig.3C**), as demonstrated in RT-qPCRs (*p<0.05 *vs.* EV, Mann Whitney test, n=3 replicates per clone), however not all reaching statistical significance. The latter comprised both “inflammatory” M1-type (e.g. *IL1B, TNFA, TLR4, STAT1, NOS2*) and “regulatory” M2-type (*CD206, PD1, PDL1, STAT6,*

*IL10*) subset markers. Taken together, this de-repression sequel seems to be independent of the polarisation state of the macrophage.

To assess the “killing” activity of macrophages towards tumor cells **(Fig.3D)** in a contact-dependent manner, 3D co-cultures were set-up between REVERBα LOF and EV clones together with the human MSS+ CRC cell line HT29. In brief, clonal THP1 monocytes were differentiated and polarised to macrophages as 2D subconfluent monolayers, followed by an overlay with parental HT29 cells at an effector:target ratio of ∼3:1 and subsequent co-culture for 5 days. Adherent cells were then subjected to colorimetric MTT viability assay. O.D. values were calculated as % of dead (= 100 - % surviving) cells ± S.E. in co-culture (CC) *vs.* the sum of the single cultures (ΣSC) (*p<0.05 *vs.* ΣSC, 2way-ANOVA with Bonferroni post-test, n=3 replicates per clone). As expected, the LOF clone reduced the overall viability of mixed co-cultures compared with the EV clone.

Conclusively, REVERBα “loss-of-function” can augment tumoricidal effector functions in THP1 macrophages.

### *PVR* KO reduces CRC cell viability

In contrast to high PVR expression in both macrophages and CRC cells, expression of TIGIT and DNAM1, the PVR counter receptors on macrophages, were very low (not shown). Therefore, homotypic PVR-PVR *cis* or heterotypic PVR-nectin/like receptor *trans* interactions are likely to be predominant in the co-cultures studied here. This observation prompted us to test if *PVR* mRNA KO itself augments anti-tumor functions of macrophages. To this end, the HT29 cell line was subjected to transfection with CRISPR/Cas9-*PVR* sgRNA KO plasmid or EV, followed by clonal selection and RNA extraction **(S3a)**. RT-PCRs using primers against a C-terminal amplicon detected loss of *PVR* mRNA **(Table S7)**, and Western blots from total cell lysates using a C-terminal Ab (*p<0.05 *vs.* EV, t-test, n=3 per clone) confirmed the knock-down of PVR protein **(S3b)**. Similar results were obtained for expression of PVR protein on the cell surface **(S3c)**. Single, live cells from A were stained with fluorescence-labelled PVR detection Ab and viability dye (7AAD) as indicated in **Table S1** and analysed by flow cytometry (FC). Intensity plots evinced loss of PVR+ cells compared to the isotype control (ITC).

Notably, loss of PVR *per se* reduced CRC cell viability **(S3d)**. HT29 clonal *PVR* KO and EV cell clones were grown as 2D monolayers to subconfluency for 1 to 5 days. Adherent cells were then subjected to MTT viability assay (*p<0.05 *vs*. EV, t-test, n=3 per clone). This assay plus representative images from bright-field (BF) microscopy revealed reduced cell counts in *PVR* KO clones compared with EV transfected cells. These data indicated that the intrinsic loss of PVR on tumor cells already yields an anti-tumoral effect, even in absence of macrophages.

### PVR KO in CRC cells or blocking Ab sensitize co-cultures to killing by THP1 macrophages

We next asked if loss or blockage of PVR augments elimination of tumor cells by macrophages. As above, monolayers of THP1 macrophages were overlaid with HT29 *PVR* KO and EV cell clones at an effector:target ratio of ∼3:1 and subsequent co-culture for 5 days. Cocultures were then subjected to MTT assay **(Fig.4A)**, which demonstrated a viability reduction of ∼40 % by macrophages, with a trend of M1 > M2, however no overt effect of the gene modification (*p<0.05 *vs.* ΣSC, 2way-ANOVA with Bonferroni post-test, n=3 replicates per clone). However, this *in-situ* assay only measured the cumulative viability (i.e. mitochondrial enzymes) of both immune and tumor cells and did not discern, which cell type was dying.

**Fig. 4.**
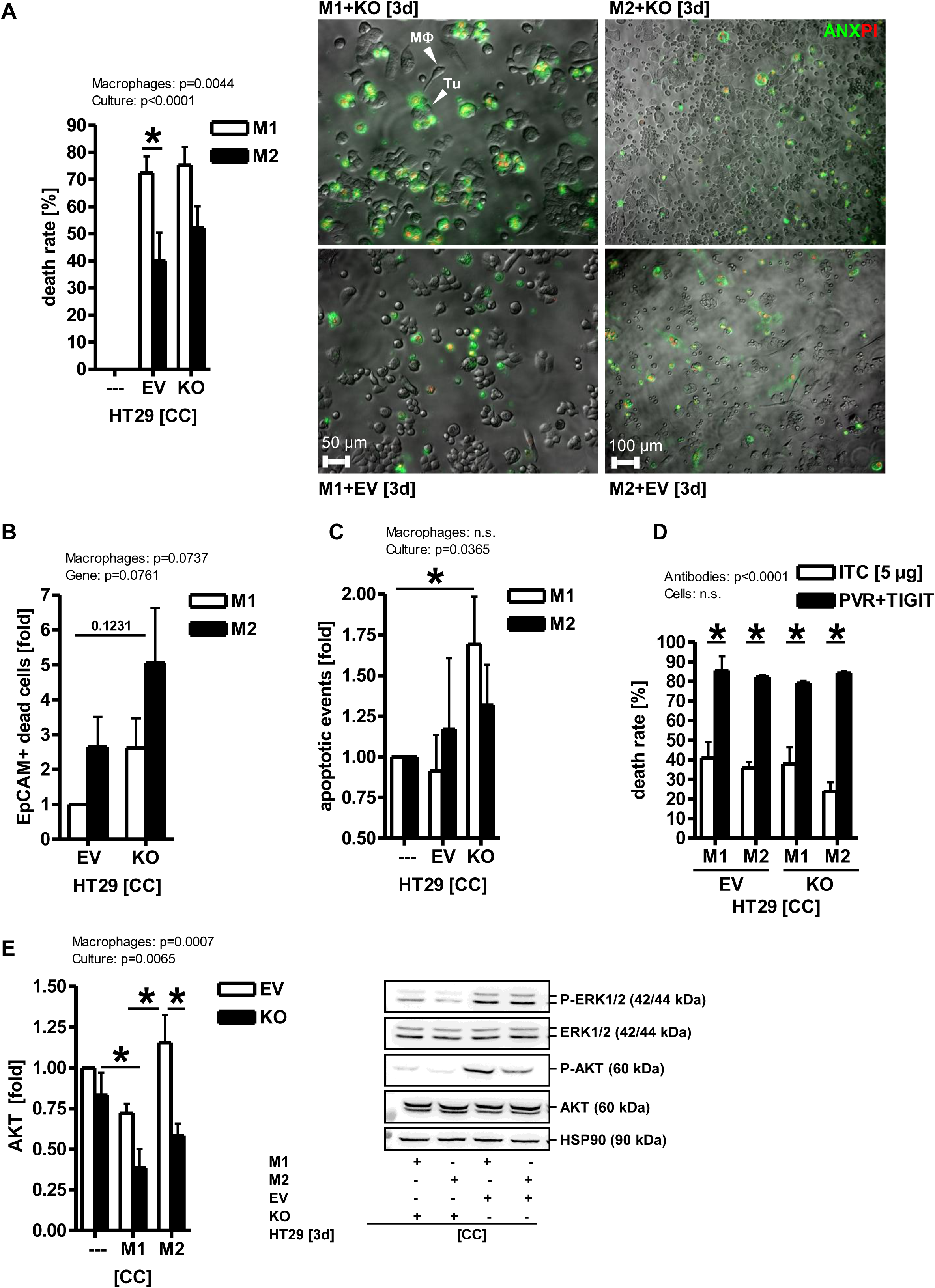
PVR inhibition reduces CRC cell viability in co-cultures with THP1 macrophages. **A**, Macrophages reduce overall viability of mixed co-cultures. THP1 monocytes were differentiated and polarised to macrophages as 2D subconfluent monolayers, followed by an overlay with HT29 clonal *PVR* KO and EV cell lines at an effector:target ratio of ∼3:1 and subsequent co-culture for 5 days. Adherent cells were subjected to colorimetric MTT viability assay. O.D. values were calculated as % of dead (= 100 - % surviving) cells ± S.E. in co-culture (CC) *vs.* the sum of the single cultures (ΣSC) (*p<0.05 *vs.* ΣSC, 2way-ANOVA with Bonferroni post-test, n=3 replicates per clone). **B,** Macrophages reduce the viability of CRC cells. Cocultures from A were dissociated with Accutase™ after 3 days, and single cells were stained with Abs (CD11b+ for macrophages, EpCAM+ for tumor cells), annexin (ANX, apoptosis) and SYTOX Blue™ (SYT, necrosis) as indicated in **Table S1** and analysed by flow cytometry (FC). Data from quadrant plots were separately calculated for EpCAM+ and CD11b+ (not shown) cell populations and expressed as % ANX+ (early apoptotic) or SYT+ (necrotic) single positive or ANX+/SYT+ double positive (late apoptotic) cells ± S.E. compared to unstained samples (*p<0.05 *vs.* EV, 2way-ANOVA with Bonferroni post-test, n=3 replicates per clone). Y-axis: “Dead” cells comprise the arithmetic Σ sum of the percentages (%) from early, late and necrotic EpCAM+ cells. **C**, Macrophages induce cell death. Cocultures were grown as in A for 3 days. Data from live cell imaging are mean numbers of annexin (ANX, apoptosis) and propidium iodide (PI, necrosis) cells per field ± S.E. and presented as -fold *vs.* SC (*p<0.05 *vs.* SC, 2way-ANOVA with Bonferroni post-test, n=10 fields per sample, n=2 replicates per clone). Representative images (top) and quantitative analyses (bottom). Color code: red = PI; green = ANX; white/grey = brightfield (BF). Original magnification 100x; Scale bars = 50-100 µm. **D**, *PVR* KO sensitizes CRC cells to PVR blocking Ab. HT29 *PVR* KO and EV clonal cell lines were co-cultured with macrophages as in A for 5 days in presence or absence of PVR and TIGIT blocking Abs (5 µg/ml, IgG1) or isotype control (ITC). Cell viability was measured by MTT assay. O.D. values were calculated as % of dead (= 100 - % surviving) cells ± S.E. in co-culture (CC) *vs.* the sum of the single cultures (ΣSC) (*p<0.05 *vs.* ITC, 2way-ANOVA with Bonferroni post-test, n=3 replicates per clone). **E**, PVR KO reduces growth and survival signaling. HT29 *PVR* KO and EV clonal cell lines were co-cultured with macrophages as in A for 5 days before extraction of total cell lysates. Representative images (right) and quantitative analyses (left) from Western blots. O.D. values of bands in gels normalized to HSP90 are -fold ± S.E. (*p<0.05 *vs*. EV, 2way-ANOVA with Bonferroni post-test, subgroup analysis with Wilcoxon signed rank test, n=3 replicates per clone).

Hence, to accurately quantify the viability of the two cell types separately, cocultures from A were dissociated with Accutase™ after 3 days, and single cells stained with fluorescence-labelled Abs against the surface marker CD11b for macrophages, EpCAM for tumor cells, annexin (ANX) for apoptosis, and SYTOX Blue™ (SYT) for necrosis detection, as indicated in **Table S1,** and subsequently analysed by flow cytometry (FC). Quantifications of quadrant plots were separately conducted for CD11b+ (not shown) and EpCAM+ **(Fig.4B)** cells and expressed as % of single positive early apoptotic (ANX+), single positive necrotic (SYT+) or double positive late apoptotic (ANX+SYT+) cells compared to unstained samples (*p<0.05 *vs.* EV, 2way-ANOVA with Bonferroni post-test, n=3 replicates per clone). This setup revealed an increased death rate in EpCAM+ *PVR* KO, both upon exposure to M1- or M2-type macrophages. Cells were consecutively subjected to microscopic imaging **(Fig.4C)**. To this end, co-cultures were set-up as in B and supplemented with propidium iodide (PI) for detection of necrosis and annexin (ANX) for apoptosis for additional 30 min. Thereafter, dying cells were counted. Again, this *in-situ* surrogate assay for contact-dependent cell death revealed an enhanced number of apoptotic events in the co-cultures with *PVR* KO cells.

Conclusively, M2-type macrophages reduced the viability of CRC cells in a contact-dependent manner to a similar extend as M1-type one (*p<0.05 *vs.* SC, 2way-ANOVA with Bonferroni post-test, n=10 fields per sample, n=2 per clone), underscoring functional plasticity and redundancy of macrophage subsets ^13^ beyond the former dichotome M1/M2 classification system ^22^.

To finally assess if *PVR* KO sensitizes to PVR/TIGIT blockage, HT29 clonal *PVR* KO and EV cell lines were co-cultured with THP1 macrophages as in A for 5 days in presence or absence of PVR and TIGIT Abs (0.1-10 µg/ml, both IgG1 each) or isotype control (ITC) **(Fig.4D)**. Cumulative cell viability was again measured by MTT assay. Notably, one observed a dominant effect of the Abs on the overall death rate, which was independent of the macrophage polarisation state or the *PVR* gene status in all concentrations of Abs tested (*p<0.05 *vs.* ITC, 2way-ANOVA with Bonferroni post-test, n=3 replicates per clone). Western blots (**Fig.4E**) on total cell lysates extracted from these co-cultures confirmed reduced amounts of the growth and survival kinase AKT upon PVR loss. Since TIGIT expression on macrophages was very low (not shown), disruption of homotypic PVR-PVR/nectin-like receptor *cis-trans* interactions were likely to account for the Ab effect observed.

These data may thus emphasize the potency of PVR targeting by an ADCC/ADCP-proficient IgG1 Ab.

### PDOs express PVR and genes related to myeloid cells

To detect expression of macrophage-ligands in a setting more relevant for the clinics, a series of PDOs from MSS+ CRC patients (n=7 cases) (**Table S3**) was investigated ^12^. First, FFPE-sections of early passage PDOs were stained for confocal immunofluorescence microscopy using phalloidin for visualisation of the actin cytoskeleton and E-cadherin as a marker for epithelial cells. Subsequent imaging emphasized the polarized structure of the organoids (**S4a)**. Single CD68+ E-cadherin-negative myeloid cells with kidney-shaped nuclei could be observed in all PDOs tested (n=3 cases) prior to any co-culture, indicative of a retainment of certain innate immune cell subsets in PDOs generated from biopsies. RT-PCRs confirmed expression of immune-related mRNAs in PDOs (n=6 cases), predominantly monocyte/macrophage markers (e.g. *CD14, CD68, CXCL11, TNFA, NOS2*) (**S4b**). Hence, we could not exclude the *a priori* inclusion of isolated myeloid cells in our co-culture systems, however, we were unable to detect lymphoid markers (e.g. *PD1)* or CTLA4 receptors (e.g. *CD86)*. This base line characterization allowed us to study the effect of myeloid cells on PDOs.

We previously ^17^ characterized immune ligand-receptor systems which mediate physical interaction between NK/T cells and PDOs using custom-made PCR arrays (n=42 genes; n=2 patients). Therein, the differential expression of checkpoint molecules allowed stratification of PDOs as responders *vs.* non-responders to NK/T cells. Hence, we were interested whether this holds true for molecules which ligate to antigen-presenting cells (e.g. macrophages). However, quantitative analysis of the PCR data **(S5a)** revealed that the exemplary NK/T cell non-responder (P22) expressed similar types and amounts of activatory (CPA) and inhibitory (CPI) macrophage-relevant checkpoint molecules as the *bona fide* NK/T cell responder (P7). Thus, unlike for NK/T cells ^17^, allogenic macrophages don’t seem to discriminate between these two patients. We therefore hypothesized that this uniform sensitivity towards macrophages follows a non-dichotome pattern and may thus be exploited for innate immunotherapy in otherwise NK/T cell-resistant patients.

This conjecture was confirmed, by retrospective analysis of the mRNA expression profiles from public Affymetrix U133 plus 2.0 arrays (NCBI GEO data set: GSE117548) **(S5b,c)** ^12^. Hierarchical clustering identified a panel of macrophage ligands on PDOs, e.g. with “*chemokine receptors*” as enriched signature (not shown), but did not uncovered any significant differences in macrophage-specific mRNA profiles between the patients (n≥47.000 transcripts; n=7 patients).

Conclusively, these array-based data ^12^ ^17^ indicated that the resistance of MSS+ PDOs to T cell immunotherapy (e.g. PD1, PDL1 Abs) may be overcome by targeting alternative checkpoints, e.g. on macrophages (**S5d**).

Our arrays also evinced that all PDOs tested expressed mRNAs encoding for *PVR/CD155* and *PVRL2/CD112* (**S6a)**. This finding was again underscored by mRNA expression profiles extracted from the NCBI GEO data set GSE117548 (n=7 patients, n=2 replicates per patient) and by Western blots using total cell lysates from selected PDOs (**S6b) (**n=2 patients, n=3 replicates per patient).

For the detection of proteins at the cell surface, PDOs were treated with vehicle (DMSO) or IFNγ (100 ng/ml) for 48 h followed by dissociation with Accutase™. Single cells were stained with fluorescent Abs (**Table S1)** and viability dye (7AAD) and analysed by flow cytometry (FC). As expected ^17^, PDOs responded with up-regulation of ICAM1 (CD54) and MHC class I (HLA-A/B/C) (**S6c)** (*p<0.05 *vs.* ITC, Kruskal Wallis test with Dunn post-test, n=3 patients, n=2 replicates per patient). Quantitative analyses of intensity plots revealed an *a priori* high expression of the nectin family receptors (PVR/CD155; PVRL2/CD112).

This result was in accordance with imaging of IHC slides (**S6d)**. FFPE-sections were stained with the same Ab as for Western blots. The majority of untreated PDOs were already positive for PVR/PVRL2, and INFγ did not further increase positivities in all PDOs tested (5 of 6 / 83 %).

### PVR blocking Ab also sensitizes PDOs to killing by primary macrophages

To now investigate the anti-tumoral efficacy of human PBMC-derived macrophages towards PDOs, 3D co-cultures were generated as above. First, the localisation of macrophages was determined by standard bright field microscopy. PMBCs were isolated from healthy donors, then adherent monocytes were differentiated and polarised to macrophages as detailed in the method section.

Intact PDOs (“spheroids”) were suspended in MatriGel™ and added, with at an effector:target ratio of 50:1, on top of the macrophage monolayer, followed by live cell imaging after 5 days.

Macrophages adhered to PDOs, but did not invade into the lumen **(Fig.5A**).

**Fig. 5.**
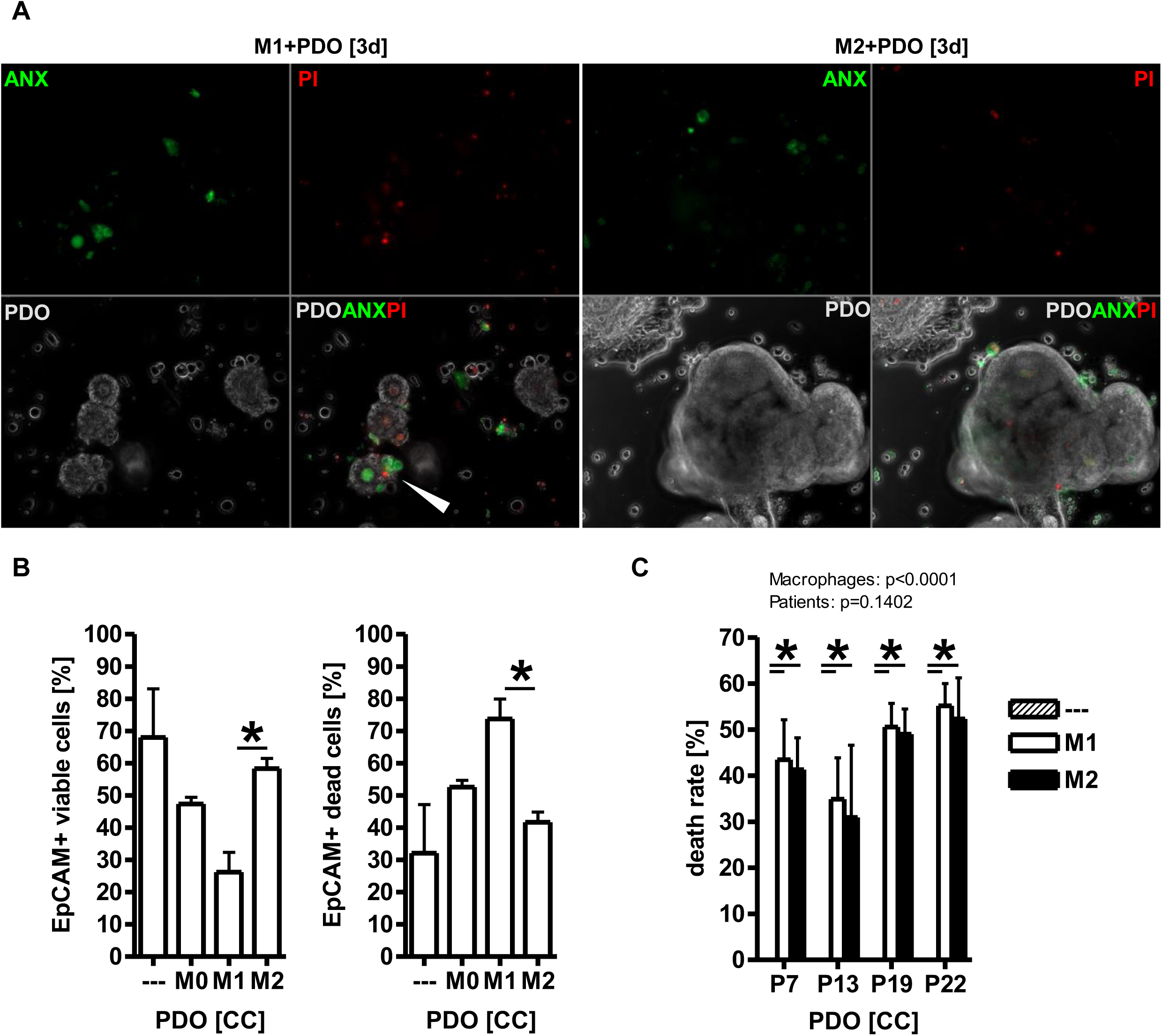
Primary blood-derived human macrophages reduce viability of PDOs in cocultures. **A**, Macrophages adhere to and induce cell death in PDOs. Monocytes from healthy donor PBMCs were differentiated and polarised to macrophages as 2D subconfluent monolayers followed by an overlay with PDOs as intact spheroids at an effector:target ratio of ∼50:1 and subsequent co-culture for 5 days in MatriGel™. Representative images from live cell imaging showcasing annexin (ANX, apoptosis) and propidium iodide (PI, necrosis) positive cells. Color code: red = PI; green = ANX; white/grey = brightfield (BF). Original magnification 100x; Scale bars = 200 µm. **B,** Macrophages reduce viability of PDOs. Cocultures from A were dissociated with Accutase™, and single cells were stained and analysed by FC as indicated in **Fig.4B** (*p<0.05 *vs.* SC, 2way-ANOVA with Bonferroni post-test, n=4 patients, n=3 replicates per patient, n=1 healthy donor per replicate). Y-axis: “Dead” cells comprise the arithmetic Σ sum of the percentages (%) from early, late and necrotic EpCAM+ cells. **C**, Macrophages reduce overall viability of mixed co-cultures. Cells from A were subjected to MTT assay. O.D. values were calculated as % of dead (= 100 - % surviving) cells ± S.E. in co-culture (CC) *vs.* the sum of the single cultures (ΣSC) (*p<0.05 *vs.* ΣSC, 2way-ANOVA with Bonferroni post-test, n=4 patients, n=3 replicates per patient, n=1 healthy donor per replicate).

To validate that Ab-target engagement is possible in our experimental set-up, FC and Western blot were employed to detect PVR expression on the surface of PBMC-derived macrophages, as consistent with others ^23^ (not shown), allowing the use of PVR blocking Ab to explore whether its efficacy can be boosted by targeting REVERBα.

To measure cell death *in situ* **(Fig.5A**), PDOs were co-cultured as in A, and viability was detected by staining with annexin (ANX) and propidium iodide (PI), followed by fluorescence imaging after 5 days. Both necrotic (PI+) and apoptotic (ANX+) cells were recorded. Importantly, also macrophages suffered from cell death, not only the tumor cells.

These finding were corroborated, when co-cultures from A had been dissociated with Accutase™, and single cells stained as before with CD11b/EpCAM Abs plus ANX/SYT and analysed by FC **(Fig.5B**). Quantitative analyses from quadrant plots showcased an increase in PDO death of ∼50 % in co-cultures compared to naïve PDOs without *prior* exposure to macrophages (*p<0.05 *vs.* SC, 2way-ANOVA with Bonferroni post-test, n=4 patients, n=3 replicates per patient, n=1 healthy donor per replicate). Here, the M1-type was more tumoricidal than the M2-type macrophage subset.

We then asked if co-culture of PDOs with macrophages in 3D reduces overall cell viabilities **(Fig.5C**). Organoids were co-cultivated as in A, followed by MTT assay after 5 days. Notably, the PDOs responded equally to allogenic macrophages, either M1 or M2 types, with a reduction in cumulative cell viability by 30-50 % compared to the single cultures (*p<0.05 *vs.* ΣSC, 2way-ANOVA with Bonferroni post-test, n=4 patients, n=3 replicates per patient, n=1 healthy donor per replicate). This data was also consistent with those obtained from the THP1-HT29 mixed co-culture system.

### REVERBα antagonist and PVR blockage augment tumor cell “killing” by macrophages

We then inquired, if the “killing” response can be boosted by combination of REVERBα ligands and blocking Abs against PVR, TIGIT and the “do-not-eat-me” receptor CD47 (as a pos. control) at the synapse of macrophages and tumor cells. To test for overall viabilities of mixed co-cultures, macrophage monolayers were again overlaid by intact PDO spheroids in MatriGel™ at a 50:1 effector:target ratio for 5 days supplemented with PVR and TIGIT blocking Abs (all ADCC/ADCP+ proficient IgG1) or isotype control (ITC, all Abs at 5-10 µg/ml). Quantitative analyses from MTT assays (**Fig.6A**) revealed that the simultaneous blockage of PVR and TIGIT increased the overall death rate of the co-cultures by ∼2-3 fold compared to the ITC (*p<0.05 *vs*.

**Fig. 6.**
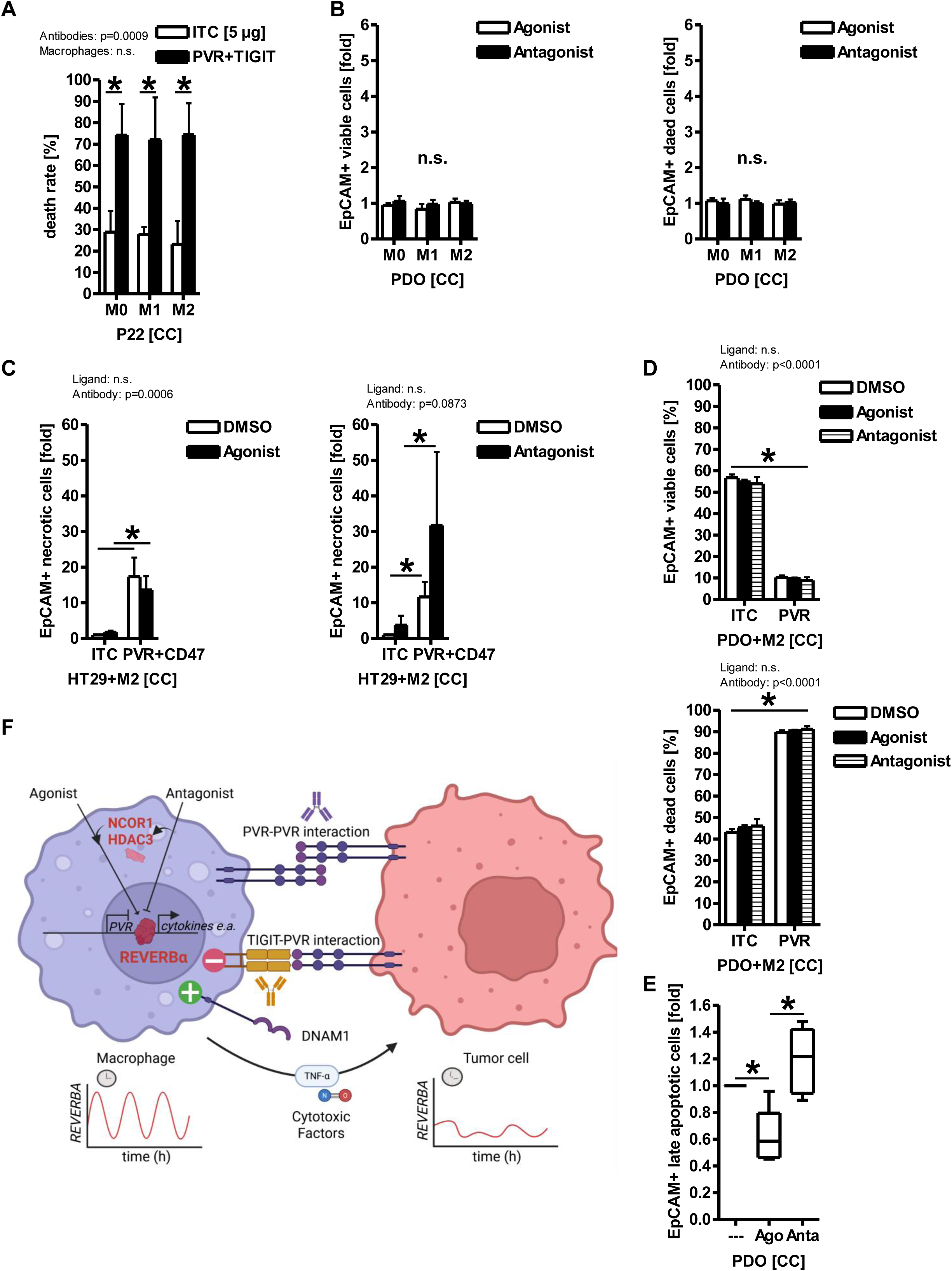
PVR blockage enhances “killing” of CRC cells by macrophages pre-treated with REVERBα antagonist. **A**, PVR/TIGIT blockage reduces overall viability of mixed co-cultures of PDOs and primary macrophages. Adherent monolayers from human PBMC-derived macrophages were overlaid with PDOs as intact spheroids at an effector:target ratio of ∼50:1 and subsequent co-culture for 5 days in MatriGel™ supplemented with PVR and TIGIT blocking Abs (5-10 µg/ml, IgG1) or isotype control (ITC). Cell viability was measured by MTT assay. O.D. values were calculated as % of dead (= 100 - % surviving) cells ± S.E. in co-culture (CC) *vs.* the sum of the single cultures (ΣSC) (*p<0.05 *vs.* ITC, 2way-ANOVA with Bonferroni post-test, n=2 patients P7/P22, n=3 replicates per patient, n=1 healthy donor per replicate). **B,** PDOs are resistant to REVERBA ligands. Cells from A were incubated for additional 48 h in absence (DMSO, vehicle control) or presence of REVERBα agonist (SR9009, 10 µM) or antagonist (SR8278, 6.7 µM), respectively. Ligand-treated macrophage monolayers were then overlaid with PDOs as intact spheroids at an effector:target ratio of ∼50:1 and subsequent co-culture for 5 days in MatriGel™ . After Accutase™ dissociation, single cells were stained and analysed by FC as detailed in **Fig.4B** (n.s., 2way-ANOVA with Bonferroni post-test, n=1 patient P22 (P7 not shown), n=3 replicates per macrophage type, n=1 healthy donor per replicate). Y-axis: “Dead” cells comprise the arithmetic Σ sum of the percentages (%) from early, late and necrotic EpCAM+ cells. Quantitative analyses are detailed in **S7**. **C,** PVR/CD47 blockage in combination with REVERBα antagonist enhances tumor cell death. Ligand-treated macrophage monolayers from B were overlaid with HT29 cells at an effector:target ratio of ∼10:1 and subsequent co-culture for 5 days in presence of PVR/CD47 Abs (as pos. control) or IgG1 isotype control (ITC, 2 µg/ml each). After Accutase™ dissociation, single cells were stained and analysed by flow cytometry (FC) as in B (*p<0.05 *vs.* ITC, 2way-ANOVA with Bonferroni post-test, n=1 patient P22 (P7 not shown), n=3 replicates per macrophage type, n=1 healthy donor per replicate). Quantitative analyses are detailed in **S8**. **D-E,** PVR blockage (**D**) in combination with (**E**) REVERBα antagonist enhances PDO death in a patient-dependent manner. Cocultures with ligand-pretreated macrophages were set-up as in B and additionally treated with PVR blocking Ab as detailed in A. After Accutase™ dissociation, single cells were stained and analysed by flow cytometry (FC) (*p<0.05 *vs.* SC, 2way-ANOVA with Bonferroni post-test, n=2 patients P22/P7, n=3 replicates per patient, n=1 healthy donor per replicate). Y-axis: “Dead” cells comprise the arithmetic Σ sum of the percentages (%) from early, late and necrotic EpCAM+ cells. Quantitative analyses are detailed in **S9**. **F, Model of REVERB**α**-PVR signaling in macrophage-tumor crosstalk.** “Responder” PDOs (P7) are sensitive, whereas “non-responder” (P22) PDOs resistant to LAK (NK/T)-mediated MHC-restricted recognition and cytotoxicity, presumably due to differential expression of regulatory immune checkpoints and other immune-ligand-receptor systems as demonstrated before ^17^. We hypothesize that these highly individualized expression profiles may be exploited by precise tailoring of innate immune-therapeutic agonistic or blocking Abs and complemented by ADCC and ADCP via non-MHC restricted macrophages. Pharmacological or genetic down-regulation of PVR on tumor and antigen-presenting cells (APC) may facilitate Ab-target engagement for agonistic receptor pairs like DNAM1, CD40, CD47 e.a. for efficient macrophage effector functions.

ITC, 2way-ANOVA with Bonferroni post-test, n=2 patients P7&P22, n=3 replicates per patient, n=1 healthy donor per replicate). Similar to the THP1-HT29 system, this effect was also dominated by the Ab and not by the macrophage type.

To discriminate between the viability of CD11b+ macrophages and EpCAM+ tumor cells and to explore if Abs and REVERBα ligands can improve the “killing” performance in the cocultures, we once more resorted to FC. PBMC-derived macrophages were incubated for 48 h in absence (DMSO, vehicle control) or presence of REVERBα agonist (SR9009, 10 µM) or antagonist (SR8278, 6.7 µM), respectively. These pre-treated macrophage monolayers were then overlaid with PDOs (P22) as intact spheroids at an effector:target ratio of ∼50:1 and subsequent co-culture for 5 days in MatriGel™ . After Accutase™ dissociation, single cells were stained as before with Abs (CD11b, EpCAM) and ANX/SYT and analysed by FC. Data from quadrant plots were separately calculated for CD11b+ and EpCAM+ cells as detailed before. Albeit, the *KRAS* mutant PDO (P22) was resistant to REVERBα ligands (n.s., 2way-ANOVA with Bonferroni post-test, n=1 patient P22, n=3 replicates per macrophage type, n=1 healthy donor per replicate) (**Fig.6B**) (**S7)**, consistent with previous studies on nuclear receptor ligands ^17^ (e.g. PPAR). Similar results were obtained for other PDOs (e.g. *KRAS* mutant P7, not shown).

To investigate if the *KRAS* wild-type (WT) CRC cell line HT29 also showcases this resistance phenotype, PBMC-derived macrophages were again pre-treated for 48 h in absence (DMSO, vehicle control) or presence of REVERBα agonist (SR9009, 10 µM) or antagonist (SR8278, 6.7 µM), respectively. Pre-treated macrophage monolayers were overlaid with HT29 cells at an effector:target ratio of ∼10:1 and subsequent co-culture for 5 days with PVR and CD47 blocking Abs (all ADCC/ADCP+ proficient IgG1) or isotype control (ITC, all Abs at 2 µg/ml). After Accutase™ dissociation, single cells were stained, analysed by FC as above.

Notably, PVR/CD47 blockage in combination with primary macrophages pre-treated with REVERBα antagonist enhanced “killing” of the CRC cells (*p<0.05 *vs.* ITC, 2way-ANOVA with Bonferroni post-test, n=1 patient P22 (&P7, not shown), n=3 replicates per macrophage type, n=1 healthy donor per replicate) (**Fig.6C) (S8)**. Interestingly, this cytotoxic effect of the Abs on EpCAM+ tumor cells was higher in co-cultures with M2- than in M1-type macrophages, which *per se* exhibited a higher *a priori* basal killing rate that could not be boosted by the treatments.

However, this discrepancy is a likely benefit, since most infiltrating tumor-associated macrophages (TAMs) belong to M2-type subsets in patients’ tissues ^22^, so their reprograming from an immunosuppressive to a more tumor-attacking mode is of clinical interest, rather then making existent M1-type subsets more cytotoxic. Taken together, either genetic or pharmacological inhibition of REVERBα improved the anti-tumoral effector functions of macrophages.

To also explore the performance of the macrophages under the same conditions, we separately analysed CD11b+ cells (not shown) by FC. Notably, we found that the two REVERBα ligands did not negatively impact the viability of macrophages, indicative that the apoptosis and necrosis signals in the FC originated from the dying tumor cells, thus, advising a use of these drugs as a potential anti-cancer treatment.

Finally, co-cultures of two selected PDOs (P7, P22), treated with REVERBα ligands and PVR Ab and as in C, were dissociated with Accutase™, and single cells stained and analysed by FC as above. Similar to the THP1-HT29 system, both PDOs were more sensitive to PVR blockage after co-culture with M2- than with M1-type macrophages (*p<0.05 *vs.* SC, 2way-ANOVA with Bonferroni post-test, n=2 patients P22/P7, n=3 replicates per patient, n=1 healthy donor per replicate) (**Fig.6D-E) (S9, S10)**. Again, the REVERBα antagonist but not the agonist lowered the viability of the PDOs, warranting future combination studies.

## Discussion

In this study, we demonstrate that, consistent with its appreciated function as a transcriptional repressor, synthetic ligands or genetic modification of REVERBα were able to target the expression of the immune checkpoint PVR in human macrophages. Consequently, dual inhibition of the REVERBα-PVR axis promoted the anti-tumoral effector functions of macrophages towards CRC cells.

Mechanistically, we found that REVERBα protein bound a RORE-like DNA-element in the -1kb human *PVR* gene promoter, and its agonist (SR9009) super-repressed, whereas its antagonist (SR8278) de-repressed transcription of *PVR* mRNA in macrophages. Macrophages with CRISPR-modified REVERBα were unresponsive to ligand, devoid of PVR protein, showed more phagocytosis, tumor cell efferocytosis and expression of genes related to host immunity (*PDL1*, *TLR4 e.a.*) than clones with the wild-type receptor. Macrophages lowered the viability of tumor cells, potentiated by PVR blocking Ab or PVR-KO in tumor cells. Consistently, REVERBα antagonist augmented tumor cell death in co-cultures with macrophages and PVR blocking Ab.

These observations were in accordance with the general knowledge that REVERBα, in its role as a transcriptional repressor ^9^, inhibits Th1/M1-type of inflammation and host immune responses ^24^ , but favors Th2/M2-type of anti-inflammatory and regenerative responses in cells and tissues. Thus, efforts have been taken to skew the balance of type-2 immunosuppressive towards immunoactivatory type-1 host responses by the development of synthetic anti-cancer drugs. As such, both exemplary ligands of the SR-series used in the current study, have been shown to exert beneficial effects in preclinical models of malignant, allergic and auto-immune diseases as well as on wound repair, pathogen clearance and metabolism (e.g. retina, vasculature, muscle, bone).

Specifically, REVERBα/β impact on chemosensitivity and resistance in solid tumors (e.g. breast cancer, glioblastoma and others).

Within the host immune system, REVERBα is not only a global repressor of the macrophage program ^25^, but also active in microglia, dendritic and mast and T-cells. Consistently, we found that genetic or pharmacological inhibition of REVERBα expression or function, triggered a global de-repression program in macrophages ^26^, which led to up-regulation of cytokine and other immune-related genes (e.g. *PDL1, TLR4*). This de-repression culminated in enhanced anti-tumoral effector functions of macrophages including phagocytosis and tumor cell efferocytosis. In contrast to a previous report that REVERBα agonist acts lethal against glioblastoma ^27^, but consistent with data from breast cancers ^28^, we evinced that REVERBα antagonist lowered tumor cell viability in direct contact co-cultures with macrophages, thus potentially advising its use as a therapeutic agent, at least in CRC. Nonetheless, the agonist was able to super-repress *PVR* and other genes (e.g. *VEGFA, PDL1*) in macrophages, in line with its appreciated role as an NCOR1-HDAC3 recruiting repressor.

Thus, to resolve the apparent paradox that *PVR* as a novel REVERBα target was subjected to de-repression by the antagonist alike other host immune factors (*ILB, IL10, NOS2, TNFA, STAT1/6* e.a.), but the antagonist was still more effective in inducing tumor cell death than the agonist which clearly evoked super-repression of the *PVR* gene (see model in **Fig.1**), we suggest the following model: In the physical synapse of our 3D cocultures, the overexpression of PVR protein on the tumor cell side, as detected in PDO and cell models by us and others, shall engage into close contact with the homo- or heterotypic PVR/nectin-like counter molecules on the macrophage, stabilising their mutual proximity. This cell-cell adhesion synapse (see model in **Fig.6**) could then be attacked by two agents, on the one hand the PVR (or TIGIT) blocking Ab, which is designed to disrupt the inhibitory signal mediated by TIGIT to unleash the activatory signal by DNAM1, though little of the former two checkpoints were detectable on macrophage, thus, this may be more relevant on T cells; on the second hand by the intracellular global de-repression of the macrophage effector program by the REVERBα antagonist.

This combination approach is expected to synergise at this cell-cell synapse to boost direct killing by soluble factors (e.g. nitric oxide, reactive oxygen) and/or ADCP or ADCC via the IgG1 Fc parts of the PVR blocking Ab bound to opsonizes tumor cells and Fc receptors on macrophages.

Likewise, the CD47 antibody directed against the “do-not-eat-me” signal on tumor cells shall exploit a similar mechanism.

Future experiments have to include physiological REVERBα ligands, such as [Fe^2+^] heme, [Fe^3+^] hemin and proto-porphyrins, previously co-crystallized with the receptor. Preliminary observations confirmed that these compounds also target *PVR* gene expression in the same reciprocal fashion as the synthetic drugs, with heme evoking super-repression *vs.* hemin de-repression, and again revealed a positive correlation between *REVERBA (NR1D1)* and *PVR* transcript levels, consistent with our finding in the CRISPR-modified clones. However, since iron/heme-binding domains are distributed among a plethora of cellular proteins, specifity of these natural ligands cannot be attributed to their binding to REVERBα, warranting further studies on second generation ligands under chemical development.

As a bridge to close the gap between the molecular and cell-based mechanisms towards the clinical setting, we could show that PDOs, alike to NK/T cells, were sensitive to macrophages, but largely resistant to REVERBα ligands, consistent with previous observations on ligands for other nuclear receptors (e.g. PPAR) ^17^. Oncogenic drivers like mutant K-RAS ^29^ or MYC ^30^ disrupt the circadian rhythm in human and murine cancers , and *vice versa*, disruption of circadian clock drives tumor initiation (e.g. *APC* mutations ^31^ ). Hence, this vicious circle may underly this PDO resistance phenomenon. In contrast, macrophages reacted well to REVERBα ligands, confirming the general notion that non-transformed primary cells exhibit a functional circadian rhythm, which is disrupted, shifted or lost in cancer cells ^32^ ^33^.

Thus, systemic administration of REVERBα ligands is expected to target the non-malignant immune compartment with a higher likeliness of efficacy than the tumor cells with a dysfunctional clock.

In contrast to the drug resistance observed in PDOs, they, however, reacted well to blocking Abs against PVR or TIGIT. This sensitivity may be explained by the fact that all tumor cell lines and PDOs tested had high expression of TIGIT receptors (PVR/PVRL2). High PVR expression in patients’ tissues *per se* is negative prognostic, marking immune exhaustion and anergy, but positive predictive regarding PVR Ab-antigen engagement and potency of the Fc-part of IgG1 blocking Abs for killing by ADCC or ADCP using Fc-receptors on innate immune cells (e.g. NK cells, macrophages). Likewise, overexpression of the HER2+ oncogene is unfavorable regarding overall survival, but allows Ab-target engagement in the patient subgroups overexpressing HER2+ such as in breast and gastric cancer, who then profit from HER2 blocking Abs. This paradox is an integral part of the therapeutic concept to strengthen the immune-tumor cell synapse. Previously, individual response profiles of PDOs (n=5 cases) allowed the stratification of single cases into “responder” (R) *vs*. “non-responder” (NR) to the cytostatic or cytotoxic activity of allogenic or autologous NK/T cells. Overall, PDOs with high TIGIT receptor (PVR/PVRL2), low MHC class I, high Ki67+ and ERK1/2+ proliferative status, were less efficiently recognized and killed by NK/T cells than those with low status for the same markers. In contrast, allogenic macrophages did not discriminate between the same PDOs.

Thus, alternative intracellular and cell surface checkpoints may undermine the efficacy of current clinical immunotherapies (e.g. with CTLA4, PD1 and PDL1 Abs). TIGIT blocking Abs (e.g. tiragolumab) are under clinical development to boost T-cell anti-tumor responses ^5^. Macrophage targeting Abs (e.g. CD47) hold the promise to achieve efficacy in an allogenic setting thereby reducing costs for application to a broader patient population. Novel insights are further expected from circadian control of PD1 immunotherapy efficacy ^34^ ^35^. However, how factors of the circadian clock may be exploited to improve clinical responses and impact chronotherapy remains to be deciphered in future clinical studies. Conclusively, co-addressing of the REVERBα-PVR checkpoint axis may represent a novel intervention strategy for patients with MSS CRC, previously excluded from immunotherapies.

## Acknowledgements

We express our gratitude to Profs. Viktor Umansky (UMM/DKFZ, Heidelberg) and Mathias Müller (MedVetUni, Vienna) for scientific advice and discussion. We want to thank Shrihar Kanikar and Alexandra Kerner for excellent technical support. We acknowledge the support of the FlowCore Mannheim and the LIMa Live Cell Imaging Mannheim at the Microscopy Core Facility Platform Mannheim (CFPM).

## Supplementary Figures

**S1 Supportive data on REVERB**α **target genes.**

**A-B,** REVERBα agonism leads to super-repression, its antagonism to de-repression of target gene mRNAs.

**A**, THP1-derived macrophages were treated with vehicle controls (DMSO), REVERBα agonist (SR9009, 1 μM) or antagonist (SR8278, 0.67 μM) for 24 h, respectively. Ct-values from RT-qPCRs were normalized to *B2M* and calculated as -fold ± S.E. (*p<0.05 *vs.* vehicle, 2way-ANOVA with Tukeýs post-test, subgroup analysis with Wilcoxon signed rank test, n=3 replicates per macrophage type).

**B,** Human PBMC-derived macrophages were treated, and data calculated as in A. (*p<0.05 *vs.* vehicle, 2way-ANOVA with Tukeýs post-test, subgroup analysis with Mann Whitney test, n=3 replicates, n=1 healthy donor per replicate).

**C,** REVERBα agonism leads to super-repression of the *PVR* promoter. HT29 and HEK293T cell lines were transiently transfected with pGL3 reporter plasmid harbouring the proximal -1 kb promoter of the human *PVR* gene, followed by 48 h treatment with REVERBα ligands (as in A). Luciferase activity was normalized to total protein content and calculated as -fold ± S.E. (*p<0.05 *vs.* vehicle, 2way-ANOVA with Tukeýs post-test, subgroup analysis with Wilcoxon signed rank test, n=3 replicates per cell line).

**S2 Structure prediction of CRISPR/Cas9-modified REVERB**α

*Phyre2* secondary structure prediction of CRISPR/Cas9-modified REVERBα. **A**, Peptide sequences (**Table S5-6**) of the C-terminal domains of WT (aa 434-611) and ΔCT (aa 132-468) REVERBα proteins. **B**, Best fit models (n=20) were generated based on PDB data. Alignment and length coverage were highest for the ligand-binding domain (LBD) of human “*nuclear receptor subfamily 1 group D member* 1” [c3n00A: Confidence 100 / Identity 100 %] for the WT/FL, and human “*retinoic acid receptor rxr-alpha*” [c5uanA: Confidence 99.8 / Identity 72 %] for the truncated protein.

**S3 PVR-KO in CRC cells by CRISPR/Cas9 sgRNA**

**A**, Loss of *PVR* total mRNA. Human CRC cell lines (MSS+: HT29 *KRAS* WT, SW480 *KRASG12V*; MSI+: HCT116 *KRASG13D*) were subjected to transfection with CRISPR/Cas9-*PVR*sgRNA knock-out (KO) plasmid or empty vector (EV), followed by clonal selection and total RNA extraction. Representative ethidium bromide-stained agarose gels visualising amplification products using primers against the C-terminus of the *PVR* mRNA and *Cas9*.

**B**, Loss of PVR total protein. Selected clonal cells from A were subjected to total cell lysate extraction. Representative images (top) and quantitative analyses (bottom) from Western blots.

O.D. values of bands in gels normalized to HSP90 are -fold ± S.E. (*p<0.05 *vs*. EV, t-test, n=3 replicates per clone).

**C**, Loss of PVR surface protein. Single, live clonal cells from A were stained with fluorescence-labelled PVR detection Ab or isotype control (ITC). Abs and viability dye (7AAD) as indicated in **Table S1** and analysed by flow cytometry (FC). Representative intensity plots are shown.

**D**, Loss of PVR *per se* reduces CRC cell viability. Selected clonal cells from A were grown as 2D monolayers to subconfluency for 5 days. Adherent cells were subjected to colorimetric MTT viability assay. O.D. values were calculated as -fold ± S.E. (*p<0.05 *vs*. EV, t-test, n=3 replicates per clone). Representative images (right) from bright-field (BF) microscopy showing reduced cell counts in KO compared with EV-transfected cells. Original magnification x 100, Scale bars = 100 µm.

**S4 Supportive data on myeloid gene signatures in PDOs**

**A**, Localization of myeloid cells in PDOs. FFPE tissue sections (n=3 patients: P7, P18, P19) were stained with CD68 Ab for immunofluorescence (IF) microscopy. Arrows mark cells with kidney-shaped nucleus. Representative images are shown. Color code: red = CD68 (monocyte/macrophage marker), green = E-cadherin (epithelial), blue = DAPI (nuclei). Original magnification x 200-400, Scale bars = 100 µm.

**B**, Expression of immune-related mRNAs in PDOs. Amplification products from RT-PCRs were separated on agarose gels and stained with ethidium bromide. Results are summarized as heatmap in false color code: BLUE = mRNA expressed *vs*. RED = mRNA undetectable (n=6 patients).

**S5 Supportive data on CP expression in PDOs**

**A**, **PCR array.** Expression profiling of 42 genes involved in recognition and binding of tumor cells by innate immune cells (e.g. NK and T cells, macrophages). cDNA from two exemplary PDOs (n=2 patients; n=14 cDNA samples) was analysed in custom-made RT^2^ Profiler PCR Arrays. Ct-values from RT-qPCRs were normalized to the mean Ct-values of n=3 house-keeping genes [*HSP90*, *ACTB*, *GAPDH*] and calculated with the ΔΔCT method as -fold change *vs.* calibrator (*p<0.05, 2way-ANOVA with Bonferroni post-test, n=2 patients) as published ^17^. Note that P7 and P22 differentially express activatory (CPA) *vs*. inhibitory (CPI) immune checkpoint genes relevant for recognition by macrophages (TNFRSF/CD40; B7H family).

**B, cDNA array.** Expression profiling of the nectin gene family. cRNAs from PDOs (n=7 patients, n=2 cDNA samples per patient) were hybridised to Affymetrix U133 plus 2.0 arrays as described ^12^ (doi: http://dx.doi.org/10.1101/660993). Anti-log2 values were calculated as means ± S.E. (*p<0.05, Kruskal Wallis test with Dunn post-test).

**C, cDNA array**. Association of *PVR* mRNA with myeloid subsets. Data were retrieved as log2 values from NCBI GEOprofiles^®^ and presented as means ± S.E. (*p<0.05, Friedman test with Dunn post-test, subgroup analysis with Wilcoxon-matched-pairs test). “Monocyte differentiation to macrophage and subsequent polarization” (HG-U133B/A; *Homo sapiens* n=3 replicate per cell type; ID32429947).

**D, Association of functional PDIO phenotypes with PDO characteristics.** Expression (IHC against PVR and PVRL2) and functional data from co-cultures were grouped by their response (R) *vs*. non-response (NR) to “lymphokine-activated killer” (LAK) CD8+ NK/T cells (right) or “killer” macrophages (left) irrespective of their polarisation status (*p<0.05, Fisher Exact test, n=6 patients). Patient-wise information is listed in **(Table S3)**. Legend: n.c. = no case recorded.

**S6 PVR expression in PDOs**

**A**, PDOs express *PVR (*and *PVRL2)* mRNA. Original log2 values were extracted from the NCBI GEO data set GSE117548 [Affymetrix Human Genome U133 Plus 2.0 Array] calculated as means

± S.E. (*p<0.05, Kruskal Wallis test with Dunn post-test, n=7 patients, n=2 replicates per patient).

**B**, PDOs express PVR (and PVRL2) protein. Organoids were treated with IFNγ (100 ng/ml) for 48 h, followed by extraction as total cell lysates. Representative images from Western blots (n=2 patients, n=3 replicates per patient).

**C**, PDOs express surface PVR (and PVRL2). Organoids were treated as in B, then dissociated with Accutase™, and single cells were stained with Abs and viability dye (7AAD) as indicated in **Table S1** and analysed by flow cytometry (FC). Representative dot (top) and intensity plots (middle) and quantitative analyses (bottom). Data are calculated from intensity plots as -fold ± S.E. PVR+ positive cells compared to isotype control (ITC) (*p<0.05 *vs.* ITC, Kruskal Wallis test with Dunn post-test, n=3 patients, n=2 replicates per patient).

**D**, PDOs express *in situ* PVR (and PVRL2) protein. Organoids from B were processed for FFPE. Sections from untreated PDOs were stained with Abs (**Table S1**) by immunohistochemistry (IHC). Representative pictures (top) and quantitative analyses (bottom). Original magnification 200x. Data are staining intensity scores: 0=negative, 1=weak, 2=moderate, 3=strong positive (n=6 patients).

Legend: n.v. = no value recorded.

**S7 Supportive data on flow cytometry in PDO cocultures**

PDOs are resistant to REVERBα ligands. Human PBMC-derived macrophages were incubated for 48 h in absence (DMSO, vehicle control) or presence of REVERBα agonist (SR9009, 10 µM) or antagonist (SR8278, 6.7 µM), respectively. Ligand-treated macrophage monolayers were then overlaid with PDOs as intact spheroids at an effector:target ratio of ∼50:1 and subsequent co-culture for 5 days in MatriGel™. After Accutase™ dissociation, single cells were stained with Abs (CD11b for macrophages, EpCAM for tumor cells), annexin (ANX, apoptosis) and SYTOX Blue™ (SYT, necrosis) as indicated in **Table S1** and analysed by flow cytometry (FC). Data from quadrant plots were separately calculated for EpCAM+ and CD11b+ (not shown) cells and expressed as % ANX+ (early apoptotic) or SYT+ (necrotic) single positive or ANX+/SYT+ double positive (late apoptotic) cells ± S.E. compared to unstained samples (n.s., 2way-ANOVA with Bonferroni post-test, n = 1 patient P22, n=3 replicates per macrophage type, n=1 healthy donor per replicate). **A**, Y-axis: “Dead” cells comprise the arithmetic Σ sum of the percentages (%) from early, late and necrotic EpCAM+ cells displayed in **B**.

**S8 Supportive data on flow cytometry in HT29 cocultures**

PVR/CD47 blockage in combination with REVERBα antagonist enhances tumor cell death. Ligand-treated macrophage monolayers from **S7** were then overlaid with HT29 cells at an effector:target ratio of ∼10:1 and subsequent co-culture for 3 days in presence of PVR/CD47 Abs (as pos. control) or IgG1 isotype control (2 µg/ml each). After Accutase™ dissociation, single cells were stained and analysed by FC as in **S7** (*p<0.05 *vs.* ITC, 2way-ANOVA with Bonferroni post-test, subgroup analysis with t-test, n=3 replicates per macrophage type, n=1 healthy donor per replicate). **A**, Y-axis: “Dead” cells comprise the arithmetic Σ sum of the percentages (%) from early, late and necrotic EpCAM+ cells displayed in **B**.

**S9 Supportive data on flow cytometry in PDO cocultures with PVR Ab**

PVR blockage in combination with REVERBα antagonist enhances PDO death in a patient-dependent manner. Cocultures from **Fig.5D-E** were dissociated with Accutase™, and single cells were stained and analysed by flow FC as in **S7** (*p<0.05 *vs.* ITC, 2way-ANOVA with Bonferroni post-test, n=2 patients P22/P7, n=3 replicates per patient and macrophage type, n=1 healthy donor per replicate). **A**, Y-axis: “Dead” cells comprise the arithmetic Σ sum of the percentages (%) from early, late and necrotic EpCAM+ cells displayed in **B**.

**S10 Gating strategy for flow cytometry of PDIOs**

Single cells from dissociated co-cultures were first analysed (**A**) for size (FSC-A) and granularity (SSC-A), and (**B**) cell aggregates / duplets were excluded by gating for FSC-W *vs*. FSC-A. Then, (**C**) a quadrant plot for 2-color stainings of EpCAM+ tumor cells *vs.* CD11b+ macrophages was generated. Therein, gates were set for (**D**) CD11b+ macrophages and (**E**) EpCAM+ tumor cells. Finally, in these gated populations the intensity and percentages of single positive SYT+ (necrotic), ANX+ (early apoptotic) and double positive ANX+/SYT+ (late apoptotic) cells were quantified, together with the number of total viable cells.

### Abbreviations

7AAD: 7-aminoactinomycin D
Ab: antibody
ADCC/ADCP: antibody-dependent cellular cytotoxicity/phagocytosis
APC: antigen-presenting cells
CC: co-culture
CD: cluster of differentiation
CPA/I: activatory/inhibitory (immune) checkpoint
CRC: colorectal cancer
CRISPR: Clustered Regularly Interspaced Short Palindromic Repeats
DBD: DNA-binding domain
Fc: fragment crystallizable (Ab isotype)
EpCAM: epithelial cell adhesion molecule
FC: flow cytometry
FL: full-length
FFPE: formalin-fixed and paraffin-embedded
H&E: hematoxylin and eosin
LI: interleukin
IF: immunofluorescence
IFN: interferon
IgG: immunoglobulin G
IHC: immunohistochemistry
ITC: isotype control
LAK: lymphokine-activated killer
LBD: ligand-binding domain
LPS: lipopolysaccharide
LUC: luciferase (reporter)
MSI/MSS: microsatellite instable/stable
Mo/s: monocyte (suspension)
M0: macrophage (adherent)
M1(IFNγ/LPS): “inflammatory” macrophages
M2(IL4/IL13): “regulatory” macrophages
NK: natural killer
DO: optical density
PBMCs: peripheral blood mononuclear cells
PD1: programmed cell death 1 (*PDCD1*)
PD-L1: programmed cell death 1 ligand 1 (*PDCD1LG*)
DO: patient-derived (tumor) organoid
PDIO: patient-derived immune (tumor) organoid
PVR: (CD155) *Poliovirus* receptor
R/NR: responder/non-responder
RAS: Rous sarcoma oncogene
REVERBα: (*NR1D1*) Nuclear Receptor Subfamily 1 Group D Member 1
RORE: retinoic acid receptor-related orphan receptor-responsive (DNA) element
SC: single culture
TIGIT: T-cell immunoreceptor with Ig and ITIM domains
TNF: tumor necrosis factor
WB: Western Blot
WT: wildtype.

