## Supplement Tables for "Macrophages target PVR/CD155 on colorectal cancer cells via REVERBα"

### Supplementary Tables

| Table S1 Antibodies (Anti-Human) | | | |
|---|---|---|---|
| **Application** | **Clone** | **Cat. No.** | **Company** |
| **Functional Grade (FC, mouse IgG1)** | | | |
| PVR (CD155) | D171 | MA5-13493 | Thermofisher |
| TIGIT | MBSA43 | 16-9500-82 | eBioscience |
| DNAM1 (CD226) | 11A8 | 338302 | BioLegend |
| NKG2D | 1D11 | 320802 | BioLegend |
| CD47 | B6H12 | 16-0479-85 | Thermofisher |
| Isotype control | P3.6.2.8.1 | 16-4714-82 | eBioscience |
| **Functional Grade (FC, human IgG1)** | | | |
| EGFR (cetuximab) | 17D25-MM | hegfr-mab1 | InvivoGen |
| CD20 (rituximab) | 16C23-MM | hcd20-mab1 | InvivoGen |
| Isotype control | QA16A12 | 403502 | BioLegend |
| **Surface Detection (FC, mouse IgG1)** | | | |
| PVR (CD155) | SKII.4 | 337618 | BioLegend |
| PVRL2 (CD112) | TX31 | 337410 | BioLegend |
| TIGIT | MBSA43 | 16-9500-82 | eBioscience |
| DNAM1 (CD226) | 10E5 | 338314 | BioLegend |
| CD107 (LAMP1) | H4A3 | 328606 | BioLegend |
| CD16 (Fc$\gamma$RIIIa) | 3G8 | 302006 | BioLegend |
| CD45 | HI30 | 304029 | BioLegend |
| CD56 (NCAM1) | HCD56 | 318310 | BioLegend |
| EpCAM | 9C4 | 324206 | BioLegend |
| CD11b | ICRF44 | 982604 | BioLegend |
| Isotype control | MOPC-21 | 400102 | BioLegend |
| **Western blot (WB)** | | | |
| REVERBA (NR1D1) | Rabbit | 14506-1-AP | Proteintech |
| REVERBA (NR1D1) | Rabbit | 13418 | Cell Signaling |
| PVR (N) (CD155) | Rabbit | NBP1-88131 | Novus |
| PVR (C) (CD155) | Rabbit | ab229553 | Abcam |
| DNAM1 (CD226) | Rabbit | A304-362A | Thermofisher |
| TIGIT | Rabbit | 99567 | Cell Signaling |
| STAT1 | Rabbit | 9172 | Cell Signaling |
| P-STAT1 | Rabbit | 7649 | Cell Signaling |
| ERK1/2 | Rabbit | 4370 | Cell Signaling |
| P-ERK1/2 | Rabbit | 4695 | Cell Signaling |
| SRC | Mouse | 2102 | Cell Signaling |
| P-SRC | Rabbit | 2105 | Cell Signaling |
| AKT | Rabbit | 4685 | Cell Signaling |
| P-AKT | Rabbit | 9271 | Cell Signaling |
| BMAL1 | Rabbit | 14020 | Cell Signaling |
| CLOCK | Rabbit | 5157 | Cell Signaling |
| HSP90 | Mouse | sc-13119 | Santa Cruz Bio. |
| Beta-Actin | Mouse | A1978 | Sigma / Merck |
| **Microscopy (IHC)** | | | |

| | | | |
|---|---|---|---|
| PVRL2 (CD112) | Rabbit | NBP1-91211 | Novus |
| PVR (CD155) | Rabbit | NBP1-88131 | Novus |
| **Microscopy (IF)** | | | |
| REVERBA (NR1D1) | Mouse | PA5-29865 | Thermofisher |
| CD68 (LAMP4) | Mouse | AM33123SU-L | Acris |
| E-Cadherin | Goat | AF648 | R&D Systems |

| **Table S2 Oligonucleotides**<br><br>Legend: CT = C-terminus with Stop codon (bold); N = N-terminal; C = C-terminal region of cDNA. | | | |
|---|---|---|---|
| **Type** | **Sequence (5' > 3')** | **Amplicon (bp)** | **Gene** |
| **sgRNA** | | | |
| *5-sgRNA-REVERBA(C)* | **CACCG** GGTCATGCTGAGAAAGGTCA | 12_2234 | *NR1D1* |
| *3-sgRNA-REVERBA(C)* | **AAAC** TGACCTTTCTCAGCATGACC **C** | | |
| *5-sgRNA-REVERBA(N)* | **CACCG** GGAGTGCCATACCTTCTCCC | 0_1117 | *NR1D1* |
| *3-sgRNA-REVERBA(N)* | **AAAC** GGGAGAAGGTATGGCACTCC **C** | | |
| *5-sgRNA-PVR(C)* | **CACCG** GTTGGCAGACTAGAGTACAG | 38_18186 | *PVR* |
| *3-sgRNA-PVR(C)* | **AAAC** CTGTACTCTAGTCTGCCAAC **C** | | |
| *5-sgRNA-PVR(N)* | **CACCG** GCTCTCGTGCTCCACCTTGC | 55_5196 | *PVR* |
| *3-sgRNA-PVR(N)* | **AAAC** GCAAGGTGGAGCACGAGAGC **C** | | |
| **ChIP** | | | |
| 5-4556RORE-*PVRp* | ACTCCTGGACCTCTGGTGATCC | 180 | *NR1D1* |
| 3-4556RORE-*PVRp* | GTTAGAGGCTTCAGTGAGCTAC | | |
| **Cloning** | | | |
| 5-KpnI-*1kb-PVRp* | ATGGTACCGTGATTTTCCTGTCTTAGCCTC | 981 | *PVR* |
| 3-HindIII-*1kb-PVRp* | ATAAGCTTCATGCCAGTTGCTCCGAGCAGC | | |
| **RT-qPCR** | | | |
| *5-REVERBA(C)* | CTGGGAGGATTTCTCCATGA | 168 | *NR1D1* |
| *3-REVERBA(C)* | TTCACGTTGAACAACGAAGC | | |
| *5-REVERBA(CT)* | GGTGCCATGGGCATGGGAGAC | 320 | *NR1D1* |
| *3-REVERBA(CT)* | TCCGGGTGGACGCCCAG**TGA** | | |
| *5-PVR(N)* | AACATGGAGGTGACGCATGT | 218 | *PVR* |
| *3-PVR(N)* | GTGACGAACAGGCAGGTGTA | | |
| *5-PVR(C)* | AGTGAGCACTCAGGCATGTC | 143 | *PVR* |
| *3-PVR(C)* | GGACACAGATGACAGTGCCA | | |
| *5-IL1B* | GGACAAGCTGAGGAAGATGC | 120 | *IL1B* |
| *3-IL1B* | TCGTTATCCCATGTGTCGAA | | |
| *5-IL10* | TTACCTGGAGGAGGTGATGC | 148 | *IL10* |
| *3-IL10* | GGCCTTGCTCTTGTTTTCAC | | |
| *5-STAT1* | TTCAGGAAGACCCAATCCAG | 112 | *STAT1* |
| *3-STAT1* | TGAATATTCCCCGACTGAGC | | |
| *5-STAT6* | CAGGGGTTATGTCCCAGCTA | 132 | *STAT6* |
| *3-STAT6* | ATGCTCATGGAGGAATCAGG | | |
| *5-ACTB* | GTCTTCCCCTCCATCGTG | 113 | *ACTB* |
| *3-ACTB* | AGGGTGAGGATGCCTCTCTT | | |
| *5-GAPDH* | GGAGCGAGATCCCTCCAAAAT | 197 | *GAPDH* |
| *3-GAPDH* | GCGTGTTGTCATACTTCTCATGG | | |
| *5-B2M* | TGCTGTCTCCATGTTTGATGTATCT | 86 | *B2M* |
| *3-B2M* | TCTCTGCTCCCCACCTCTAAGT | | |

**Table S3 Patients Characteristics**

* Detailed genetic & clinical information (MSS+ CMS2 CRC) is presented by Betge J. et al. 2022.

§ Immunohistochemistry (IHC) on FFPE sections using Abs listed in (**Table S1**).

Legend: wt = wild type; MSS = microsatellite stable; CMS = consensus molecular subtype;
FAP = familial adenomatous polyposis; pTU = primary tumor tissue; AC = adenocarcinoma;
UICC = tumor staging.

| **Patient** | **PDO** | | | | **pTU** | | |
|---|---|---|---|---|---|---|---|
| **ID** | ***APC*** | ***KRAS*** | **§ PVR+** | **PVRL2+** | **H&E** | **AC** | **UICC** |
| **P18** | **wt** | **wt** | **+** | **-/+** | - - - | Sigma | II |
| **P13** | **mut** | **wt** | **-** | **-** | Inflammation | Rectum | I |
| **P30** | ***SMAD4*** | ***A146T*** | **+** | **+** | Inflammation | Ascendens | II |
| **P19** | **mut** | ***A146T*** | **+** | **+** | Inflammation | Sigma | I |
| **P7** | **mut** | ***G12D*** | **+** | **+** | Necrosis (FAP) | Rectum | III |
| **P21** | **mut** | ***G12D*** | **+++** | **-/+** | - - - | Rectum | I |
| **P22** | **mut** | ***G12C*** | **+** | **+++** | n.a. | Ascendens | IV |

**Table S4 Nucleotide sequence alignment.**

NCBI BlastN (top) of Alibaba version 2.1 (bottom) prediction of potential REVERBα binding elements (RORE) in the upstream regulatory DNA sequence of the human *PVR/CD155* gene (Genbank® ID: NG_008781.2; 5.30 kb region from 1 to 5302 bp). A target site for REVERBα binding (4556-4561) was localized in the -1 kb proximal promotor. Color legend: **yellow** = other transcription factor sites (e.g. AHR); **pink** = RORE; **green** = CDS (5300 > 5302); **blue** = mRNA (5113 > 5302); *underlined* = PCR amplicon, **bold** = ChIP amplicon.

4321 *gtgattttcc tgtcttagcc tc*cccagtag ctgggattac aggcatgtgc caccatgccg
4381 gctaattttt gtatttttag taaagatggg tttttgccat gttggccagg ctggtcttga
4441 **actcctggac ctctggtgat cc**tcctgcct cggcctccca aagtgctggg attacaggcg
4501 tcagccactg cgcttgctcc cccagccccc aaactttttt tttttttttt gagac**aggtc**
4561 tggctctgcc acccaagctg tagtgtagtg gcgcaatg**gt agctcactga agcctctaac**
4621 tcctgggttc acacagtcct cctgcctcag cctcccaaaa tgctggaatt caggcctgag
4681 ccaccgcgcc ctactatgcc catttttcag ataaagaatc tgagggccct cgaggagaaa
4741 ccacttgccc aacatgagcc ctggtcttgt ctgaccccaa agcccatggg aagttta**ggc**
4801 **tgcgtgg**aag gacagcctgg tgggctcagg atctgtccca tcacgagttg gaacctcagc
4861 tctgccactc ggcaaccgcg agccgttagt aagttgctga atctctccgt gtgcctccgt
4921 ttcctcacct ggaatgtggg cgacctcaca caggaagccg cctcttctag tgggcaccga
4981 cggagcgagc gcgcgtctgc agcgggactg agcgccggga gagacctgcg caggcgcagg
5041 cgcgcgggga gggccagcct gggtggccca ccccgcgcct ggcgggactg gccgccaact
5101 cccctccgct cc**agt**cactt gtctggagct tgaagaagtg ggtattcccc ttcccacccc
5161 aggcactgga ggagcggccc cccggggatt ccaggacctg agctccggga gctggactcg
5221 cagcgaccgc ggcagagcga gcgggcgccg ggaagcgagg agacgcccgc gggaggccca
5281 *gctgctcgga gcaactggc* **atg**

```
==================================================================================
seq(  180..  239)     tcagccactgcgcttgctcccccagcccccaaacttttttttttttttttgagacaggtc
Segments:
1.1.1.6    174  183   BP1=
2.3.3.0    174  183   ind=
1.1.1.1    177  186   c-Jun==
2.3.1.0    181  190    ====Sp1===
1.1.3.0    199  208                      =C/EBPalp=
2.3.1.0    200  209                       ====Sp1===
2.3.2.3    200  209                       ====WT1===
1.1.1.5    212  221                                   ====GCN4==
3.3.2.0    213  222                                    ===HNF-3B=
2.3.2.2    214  223                                     =====Hb===
2.3.2.2    221  230                                            ===LyF-1==
2.1.2.3    230  239                                                     =REV-ErbA=
==================================================================================
seq(  240..  299)     tggctctgccacccaagctgtagtgtagtggcgcaatggtagctcactgaagcctctaac
Segments:
2.3.1.0    240  249   ====Sp1===
3.5.1.2    243  252      ===Adf-1==
9.9.539    247  256          ====NF-1==
1.1.3.0    267  277                              =C/EBPalpha
==================================================================================
```

**Table S5 Nucleotide sequence alignment.**

NCBI BlastN of sgRNA nucleotide sequences in the human *NR1D1* mRNA Genbank® ID: NM_021724.5. The target site for the C-terminal sgRNA (1878-1897 bp) was localized in the C-terminal exon E6 (1744-1929 bp), the N-terminal sgRNA in the 5-UTR upstream of the ATG start codon (not shown). Color legend: **blue** = E6; **pink** = sgRNA targeting site; **green** = start codon CDS; **yellow** = stop codon CDS.

```
        1 agagtgaaat attactgccg agggaacgta gcagggcaca cgtctcgcct ctttgcgact
       61 cggtgccccg tttctcccca tcacctactt acttcctggt tgcaacctct cttcctctgg
      121 gacttttgca ccgggagctc cagattcgcc accccgcagc gctgcggagc cggcaggcag
      181 aggcaccccg tacactgcag agacccgacc ctccttgcta ccttctagcc agaactactg
      241 caggctgatt cccctacac actctctctg ctcttcccat gcaaagcaga actccgttgc
      301 ctcaacgtcc aacccttctg cagggctgca gtccggccac cccaagacct tgctgcaggg
      361 tgcttcggat cctgatcgtg agtcgcgggg tccactcccc gcccttagcc agtgcccagg
      421 gggcaacagc ggcgatcgca acctctagtt tgagtcaagg tccagtttga atgaccgctc
      481 tcagctggtg aagacatgac gaccctggac tccaacaaca acacaggtgg cgtcatcacc
      541 tacattggct ccagtggctc ctccccaagc cgcaccagcc ctgaatccct ctatagtgac
      601 aactccaatg gcagcttcca gtccctgacc caaggctgtc ccacctactt cccaccatcc
      661 cccactggct ccctcaccca agacccggct cgctcctttg ggagcattcc acccagcctg
      721 agtgatgacg gctccccttc ttcctcatct tcctcgtcgt catcctcctc ctccttctat
      781 aatgggagcc cccctgggag tctacaagtg gccatggagg acagcagccg agtgtccccc
      841 agcaagagca ccagcaacat caccaagctg aatggcatgg tgttactgtg taaagtgtgt
      901 ggggacgttg cctcgggctt ccactacggt gtgcacgcct gcgagggctg caagggcttt
      961 ttccgtcgga gcatccagca gaacatccag tacaaaaggt gtctgaagaa tgagaattgc
     1021 tccatcgtcc gcatcaatcg caaccgctgc cagcaatgtc gcttcaagaa gtgtctctct
     1081 gtgggcatgt ctcgagacgc tgtgcgtttt gggcgcatcc ccaaacgaga gaagcagcgg
     1141 atgcttgctg agatgcagag tgccatgaac ctggccaaca accagttgag cagccagtgc
     1201 ccgctggaga cttcacccac ccagcacccc accccaggcc ccatgggccc ctcgccaccc
     1261 cctgctccgg tcccctcacc cctggtgggc ttctcccagt ttccacaaca gctgacgcct
     1321 cccagatccc caagccctga gcccacagtg gaggatgtga tatcccaggt ggcccgggcc
     1381 catcgagaga tcttcaccta cgcccatgac aagctgggca gctcacctgg caacttcaat
     1441 gccaaccatg catcaggtag ccctccagcc accaccccac atcgctggga aaatcagggc
     1501 tgcccacctg cccccaatga caacaacacc ttggctgccc agcgtcataa cgaggcccta
     1561 aatggtctgc gccaggctcc ctcctcctac cctcccacct ggcctcctgg ccctgcacac
     1621 cacagctgcc accagtccaa cagcaacggg caccgtctat gccccaccca cgtgtatgca
     1681 gccccagaag gcaaggcacc tgccaacagt ccccggcagg gcaactcaaa gaatgttctg
     1741 ctggcatgtc ctatgaacat gtacccgcat ggacgcagtg ggcgaacggt gcaggagatc
     1801 tgggaggatt tctccatgag cttcacgccc gctgtgcggg aggtggtaga gtttgccaaa
     1861 cacatcccgg gcttccgtga cctttctcag catgaccaag tcaccctgct taaggctggc
     1921 acctttgagg tgctgatggt gcgctttgct tcgttgttca acgtgaagga ccagacagtg
     1981 atgttcctaa gccgcaccac ctacagcctg caggagcttg gtgccatggg catgggagac
     2041 ctgctcagtg ccatgttcga cttcagcgag aagctcaact ccctggcgct taccgaggag
     2101 gagctgggcc tcttcaccgc ggtggtgctt gtctctgcag accgctcggg catggagaat
     2161 tccgcttcgg tggagcagct ccaggagacg ctgctgcggg ctcttcgggc tctggtgctg
     2221 aagaaccggc ccttggagac ttcccgcttc accaagctgc tgctcaagct gccggacctg
     2281 cggaccctga acaacatgca ttccgagaag ctgctgtcct tccgggtgga cgcccagtga
     2341 cccgcccggc cggccttctg ccgctgcccc cttgtacaga atcgaactct gcacttctct
     2401 ctcctttacg agacgaaaag gaaaagcaaa ccagaatctt atttatattg ttataaaata
     2461 ttccaagatg agcctctggc cccctgagcc ttcttgtaaa tacctgcctc cctcccccat
     2521 caccgaactt cccctcctcc cctatttaaa ccactctgtc tcccccacaa ccctcccctg
     2581 gccctctgat ttgttctgtt cctgtctcaa atccaatagt tcacagctga

Gene   1-2630
Exon1  1-526
CDS    496-2340
Exon2  527-865
Exon3  866-954
Exon4  955-1099
Exon5  1100-1743
Exon6  1744-1929
Exon7  1930-2140
Exon8  2141-2630
polyA_site 2630
```

**Table S6 Amino acid sequence alignment.**

NCBI BlastP of the human wild-type (WT) full-length (FL, aa 1-614) and the C-terminally truncated ΔCT (aa 1-468) proteins.
UniProt® ID: Query = REVERBA, WT/FL (NP_068370.1; P20393_HUMAN); Subject = REVERBA, ΔCT.

```
FL  1    MTTLDSNNNTGGVITYIGSSGSSPSRTSPESLYSDNSNGSFQSLTQGCPTYFPPSPTGSL  60
ΔCT 1    MTTLDSNNNTGGVITYIGSSGSSPSRTSPESLYSDNSNGSFQSLTQGCPTYFPPSPTGSL  60

FL  61   TQDPARSFGSIPPSLSDDGSPSSSSSSSSSSSSFYNGSPPGSLQVAMEDSSRVSPSKSTS  120
ΔCT 61   TQDPARSFGSIPPSLSDDGSPSSSSSSSSSSSSFYNGSPPGSLQVAMEDSSRVSPSKSTS  120

FL  121  NITKLNGMVLLCKVCGDVASGFHYGVHACEGCKGFFRRSIQQNIQYKRCLKNENCSIVRI  180
ΔCT 121  NITKLNGMVLLCKVCGDVASGFHYGVHACEGCKGFFRRSIQQNIQYKRCLKNENCSIVRI  180

FL  181  NRNRCQQCRFKKCLSVGMSRDAVRFGRIPKREKQRMLAEMQSAMNLANNQLSSQCPLETS  240
ΔCT 181  NRNRCQQCRFKKCLSVGMSRDAVRFGRIPKREKQRMLAEMQSAMNLANNQLSSQCPLETS  240

FL  241  PTQHPTPGPMGPSPPPAPVPSPLVGFSQFPQQLTPPRSPSPEPTVEDVISQVARAHREIF  300
ΔCT 241  PTQHPTPGPMGPSPPPAPVPSPLVGFSQFPQQLTPPRSPSPEPTVEDVISQVARAHREIF  300

FL  301  TYAHDKLGSSPGNFNANHASGSPPATTPHRWENQGCPPAPNDNNTLAAQRHNEALNGLRQ  360
ΔCT 301  TYAHDKLGSSPGNFNANHASGSPPATTPHRWENQGCPPAPNDNNTLAAQRHNEALNGLRQ  360

FL  361  APSSYPPTWPPGPAHHSCHQSNSNGHRLCPTHVYAAPEGKAPANSPRQGNSKNVLLACPM  420
ΔCT 361  APSSYPPTWPPGPAHHSCHQSNSNGHRLCPTHVYAAPEGKAPANSPRQGNSKNVLLACPM  420

FL  421  NMYPHGRSGRTVQEIWEDFSMSFTPAVREVVEFAKHIPGFRDLSQHDQVTLLKAGTFEVL  480
ΔCT 421  NMYPHGRSGRTVQEIWEDFSMSFTPAVREVVEFAKHIPGFRDLSQHDQ

FL  481  MVRFASLFNVKDQTVMFLSRTTYSLQELGAMGMGDLLSAMFDFSEKLNSLALTEEELGLF  540

FL  541  TAVVLVSADRSGMENSASVEQLQETLLRALRALVLKNRPLETSRFTKLLLKLPDLRTLNN  600

FL  601  MHSEKLLSFRVDAQ  614

Region 127-215 "NR_DBD_REV_ERB" DNA-binding domain with C4-type zinc fingers
Region 418-611 "NR_LBD" Ligand-binding domain
Site   448-473 "Coregulator recognition site"
```

**Table S7 Nucleotide sequence alignment.**

NCBI BlastN of sgRNA nucleotide sequences in the human *PVR* mRNA Genbank® ID: NG_008781.2. The target site for the C-terminal sgRNA (23496-23515 bp) was localized in exon E8, the N-terminal sgRNA (11246-11265 bp) in exon E3. Color legend: **blue** = Exon; **pink** = N-terminal sgRNA targeting site; **red** = C-terminal sgRNA targeting site; **green** = start codon CDS; **yellow** = stop codon CDS.

```
   1 aataaataac tttttagacc atatatttct tttttttttt tttttttttt tgagacagag
  61 tctcactctg ttgcccagac tggaatgcag tggcatgacg ttgactcact gtaacctcca
 121 cctcaagtga ttcagcctcc caagtagctg ggattacagg tgtgcacgac taccacccag
 181 ctaatttttg tgtttttagt acagatgggg tttcaccatg ttggccaggc tggtcttgaa
 241 ctcctgacct caaaagtgtg ggggaaagaa agagagatca gactgttact gtgtctatgt
 301 agaaaaagga agacataaga aactccattt tgatctgtac taagaaaaat tcttctgcct
 361 tgagatgctg ttaatctgta accctagccc caaccctgtg ctcgcagaaa cgtgtgctgt
 421 attgactcaa ggtttaatgg atttagggct gtgcaggatg tgctttggta aaaatgtgtt
 481 tgcaggcagt atgcttggta aaagtcattg ccattctcca gtctcaagta accagggaca
 541 ccatacactg cggaaggcag cagggacctc tgcccaagaa aacctgggta ttgtccaaga
 601 tttcccccac tgagacagcc tgagagatgg cctcgtggga agggaaagac ctgaccgtcc
 661 cccagcccga cacccataaa aggtctgtgc tgaggaggat tagtgaaaga ggaaggcctc
 721 tctgcagttg agataagagg agggcatctg tctgctgctc gtccctggga atggaatatc
 781 tcggtgtaaa acccgatcgt acattctatt tactgagata ggagaaaact gccttatggc
 841 tggaggtgag acacactggc agcaatactg ctctttactg cactgagata tttgtgtaaa
 901 gtcaaacata aatctggcct acgtgcacat ccaggcacag ctcctttcct taaatttatc
 961 tatgacaccg agtcctttgc tcacgttttc ctgctgaccc tctccccacc attaccctat
1021 agtcctgcca catccccctc accgagatag tagggatagt gatcaataaa tactgaggga
1081 actcagagac cagtgccggt gcaggtcctc acttgctgag caccggtccc ctgggcccac
1141 ttttcttcct ctatactttg tctctgtgtc ttatttcttt tctcagtctc tcgtctccac
1201 cttgtgagaa atactcagag gtgtggaggg gcaggccccc ttcaaaatga tccacccacc
1261 tcagcctccc aaagtgctgg gattacaggc atgagccacc atgcccagcc acatataact
1321 tatttctaaa taacttattt aactaacacc acatgatttt atgacccccct attttatgga
1381 tgagaaaaaa gaggcaagag aagcgatacc agttgcttag gattcccagc taccaattag
1441 tggaagcaga atttgaacca agggagtctc tggagccctc gttcccaatc tccttttctt
1501 cctctccatg tctttctttc tccccctgct cccaggcctc ctacatcaca cccttctgga
1561 caaagcttca catctgacat tttctttttt ggggtggaca gagtcttact ctgtcaccga
1621 ggctggaggg ccgtggcgca atcatggctc atggcagcct caaactcctg ggcttgaggg
1681 atcctccagc ctcagcctcc caagtagctg ggactatggg catgtgccac cacacctggc
1741 taattttttta atttttgta gaaatacagg gggtgggggc atatctgaaa attttaaacc
1801 ttccctgact gccgtgagca tttctgggta gaagccactc cacgacctta ctttgctggc
1861 atgagtgctg gacagcaatt tggcaacatc ctctagacca caaatgtgca tggcctttgg
1921 cccagcaaat cagcttctgg gaataaatat gacagatact gttacataca tgggaaatgc
1981 catggacatg atgacccagg gaagcacagt ttgcaagtac aaaagggctg gaaacaccca
2041 aaaagcccat cccctgaaga cagtttcaaa aatgacggtg cattcatgta atggaatagt
2101 acacatttgt gaatgaatga atgaggtagc tcaatatatc tgcctatggg aaaatcacca
2161 agatatgttg ctaagtgaaa aacagaaaga tacagaataa tgctaccatt ttattttta
2221 atgggagagg gaagaataag gaaatggatt catatttgct taattacaca agcttcctga
2281 tgatcctttt ggatccagaa gtttgaaaac cacagcatca agaggatgcc cattgggcat
2341 ggtggctcac gcctgtaatc ccagcacttt gggaggctga ggagggtgga tcgcctaagt
2401 tcaggagttc gagaccagcc tggccaacat ggtgaaaccc catctctact aaaaacacaa
2461 aaaaattagc tgggtatggt ggtgcaggcc tgtaattcca gctacttggg aggctgaggc
2521 gcaagaatca cttaagcctg ggaggcggag gctgccgtga gccgagatca cgccattgca
2581 ctccagcttg ggagacagag caagactgtc tcaaaacaaa caaacacaca caaaaaataa
2641 gatgcccatt ggagatctcc attcactatt ttatcagatg atatttcagc acttgagtga
2701 agaaaatgtc tacaactggt ccaaacactc aaggaattta gactcttgac agagaaacac
2761 acagaagcaa gcagatatgc cccgaaccaa acttgctttt tcttctaagc ctgcttcttc
2821 ggagcagcct ctgaccctcc agggccctga gaggagtccc tggagccaca ggagcttggg
2881 agcccagaga agtgggggaa tcccagatgg tgacagcatc atccagagga ggaacataac
2941 gatggccaca gattcccaat ttggaaataa ctcagcaaaa caacactgac agaaaattca
3001 gagatttcca agagactagc ctccctgttt ccatttagaa attcacctac atacatccag
3061 ggtgagagac cccaagcatc agatgtgtca tacacaagca tcagatccaa gttagagagg
3121 gcacccgggg acagaaacct ccaagccagc caaaagccaa aggaaggatt ttacagaaag
3181 atacaggaat gtggggccag gcacagtggc tcatgcctgc aatcctagca ctttgggagg
3241 ccgaggtggg aggatcgctt gcggccagga attcaaggcc agcctgagca acatagcgag
3301 atcctatctc tacaacaaat ttaaaaacta gctgggtgtg atcacatatg cctgtggtcc
3361 caactaccca ggaggctgag gcaggagttg agcccaggag gtggaggctg caatgagcaa
3421 tgatggcgcc acttcactcc agcctcagtg aaatagtgag accctgtcta aaaataataa
3481 taaaaataaa aaacatataa agatgtcaga gcacagggac cagagctgtt agtcggcctc
3541 atggagaccc tggccaggac agccacctct ggggaccagg aagtctctca gtttctgtct
3601 ctgagggttt ggctctctga gtctctctgt cacctctcat ttctctctca tctccctctg
3661 tctatgctgt gtctctatct ctttcaatct ctctgtctct gtctcccagc ccttatctct
3721 gagtgtgtct gtttctctct catttctctc tctctctctc tctctttctc tctctctctt
3781 tctctctctc tctttctctc tccccctcc ctcctcccct ctccctactg caggctggtc
3841 cataaccctt atcttcccca gacctgatgg agttctggta ttaatctgtt tatggtctgt
3901 ttccctccac tagaacacgg tcttcattaa aggcagagat ttttgtctgt cttattttct
3961 gctgtatacc cagtgcctgg gatactggct ggatcaataa atatatgctg aatgaagaat
4021 gaatgaatga ccattctcat gaacattatt atttgatgcc aactgtaaag caacaataat
4081 aataattgtc actgccactg ctgctattgt tggtatatca gctttttatt tagtacaaac
4141 tatatccccg aaattgagct aggcagtttc ttgtttaagt tttacagacc acgctgtgag
4201 gtaagtactg ttagtatgcc ctttttttctt tctttttgag acagagtctc actctgtcgc
```

4261 ccaggctgga gtgcagaggc ccgatcttgg ctcactgcaa cctctgcctc ccgggtacaa
4321 gtgattttcc tgtcttagcc tccccagtag ctgggattac aggcatgtgc caccatgccg
4381 gctaattttt gtatttttag taaagatggg tttttgccat gttggccagg ctggtcttga
4441 actcctggac ctctggtgat cctcctgcct cggcctccca aagtgctggg attacaggcg
4501 tcagccactg cgcttgctcc cccagccccc aaactttttt tttttttttt gagacaggtc
4561 tggctctgcc acccaagctg tagtgtagtg gcgcaatggt agctcactga agcctctaac
4621 tcctgggttc acacagtcct cctgcctcag cctcccaaaa tgctggaatt caggcctgag
4681 ccaccgcgcc ctactatgcc catttttcag ataaagaatc tgagggccct cgaggagaaa
4741 ccacttgccc aacatgagcc ctggtcttgt ctgaccccaa agcccatggg aagtttaggc
4801 tgcgtggaag gacagcctgg tgggctcagg atctgtccca tcacgagttg gaacctcagc
4861 tctgccactc ggcaaccgcg agccgttagt aagttgctga atctctccgt gtgcctccgt
4921 ttcctcacct ggaatgtggg cgacctcaca caggaagccg cctcttctag tgggcaccga
4981 cggagcgagc gcgcgtctgc agcgggactg agcgccggga gagacctgcg caggcgcagg
5041 cgcgcgggga gggccagcct gggtggccca ccccgcgcct ggcgggactg gccgccaact
5101 cccctccgct ccagtcactt gtctggagct tgaagaagtg ggtattcccc ttcccacccc
5161 aggcactgga ggagcggccc cccggggatt ccaggacctg agctccggga gctggactcg
5221 cagcgaccgc ggcagagcga gcgggcgccg ggaagcgagg agacgcccgc gggaggccca
**5281** gctgctcgga gcaactggc**a** **tg**gcccgagc catggccgcc gcgtggccgc tgctgctggt
5341 ggcgctactg gtgctgtcct ggccaccccc aggaaccggt gagtgacccc cgcgcagtcc
5401 ggtggcccct gtctggctcc cagctccgca tccaggccgg gtactcgccc ccttgggttc
5461 ccgaagatga cggccccgcc cggcgccacc caccccccat ctctgccccg gggttggaag
5521 gaacacggtg cgagggtgcg agggcaggac cctcctcttg gccacccgga gagcggtcgg
5581 agttgggtac agaaattcct cgaaattggc agcaacacgg gtcgggcaca agaccctcct
5641 ctgggcccct cggaaagggg atggagttgg gtgaggaaat ttcccaaaaa ggcaagtgaa
5701 tcccggaaaa tcagcagatg gtgacgcatg ccggattacc ttacccaggt tcctgctcag
5761 catacccagg ccataaccca gcagcgtgct ggcaggctgg gtctggacgg aggggatggt
5821 tgatgttggg gaggaggaca gtgggtgaca cttgacacca ttacccctct gacccccaat
5881 gtctgactcc aggggcctgg ccattcattc attctttcat tctttttttt ttttttttga
5941 gacagtctca ctccatcacc caggctggag tgcagtgtcc tgctctcggc tcactgcagc
6001 ctcctcctcc cgagttcaag cgattctcct gcttcagcct cccgagtagc tcggattaca
6061 gacatgtgcc actatgcctg gctaatattt ttgtattttt agtagataca ggatttcacc
6121 atgttggcca ggctggtctc aaactcctga taaatatata tattatatat tattattata
6181 tattacatat tatagtaata tataataaaa atatataata tatattatag taatatataa
6241 tatataataa aatatataat atatattata gtaatatata atatataata aaatatataa
6301 tatatattat agtaatatat aatatattat aatatattat atatattatt ataatatata
6361 atatgtatat tatatattat atatattatt ataatatata atatgtatat tatatattat
6421 atatattatt ataatatata atatgtatta tgtattatat aattgataat atataataat
6481 atatttctta ataaatatat attaaatata ttatatattt ataacatata ttatttatat
6541 aataaataat atatatttta ttatacattt aaagagtaaa tagataacat attttattat
6601 ttatttatat gttttgtgta tatatatata tatatatata tatatatagt gcgtgtgtgc
6661 gtgtatgtgt gtgtgtgtat gagagagaca aggtcttgct ttgttgccca ggctggagtg
6721 cagtggtaca atcccggctc attgcaaact cgacttccca ggctcaagcg atcctctcac
6781 tgtggcttct ggagtagctg ggattatagg tgctcacctg gctaattttt tttgtatttt
6841 tggtagagac agggtttcgc tatgctgccc aggctggttg cagccaagca cagtggctca
6901 tgcctgtaat cccaacactt tgggaggctg tggtgggagg atcacttgag cttaggaatt
6961 caaaaccagc ctgggcaata tggcaagacc ccatctctac aaaaataatt aactaaataa
7021 aaatttaaaa ataaaaaaga atgcatagga ccatgtttat tagtgtcaat aagcgctgtc
7081 acattttgag gacctactat gtgctagggc ccttagcaga aagattttta tataggacct
7141 gggggattag aggtcagtct ctgaccgccc aagttcaaat cccacctctg ccactaaacc
7201 tctgtatgac ttggaggaca ctatactctg tgtcagtttc cctgtctgta aaatgatgtt
7261 aacagtagtg cttgcctcag gggggctatt tgaggattaa atgccaagat atatgtgaag
7321 cacccagaat agcccctgat ctgtgtcagc accctgtgag tgtccactgt tctaagtctc
7381 attttacaaa ctaggaaaaa gaatcagaga ggttaagtaa ccggcctgaa gctccacagc
7441 agaaaactga aggacccaa aactcctggg ctataattgc cttcctctcg tgcctcaatc
7501 ccccacattc ttccaaccag gacccaccca gagtaataat gcagtgtttc atggatttga
7561 agacccaaga tttctccccg ttttctacat tggggttatt ctaaccccta atggcaagtt
7621 gtgctataaa tgacaacagt catggtatgt gaaacctacg acactggtgc gatgaaaaag
7681 ttctgggcca ggtgcggtgg ctcatacctg taatcccagc actttgggag gccgtagtgg
7741 gaggatcact tgaggtcagg agttcaagac cagcctggcc aacatggtga aaccccatct
7801 ctgctaaaaa caaaaataca aaaaattagc caggtgtggt ggtgggagcc tgtcatccca
7861 gctacgcgga aagctgaggc aggagaaccg cttgaacccc agaggcagag cttgcagtga
7921 gcagagatcg cgccactgca ctccccctgg gcgacagaga gagactccgt ctcaaaaaaa
7981 aaagagaaga aaagaaaaag ttctggcgct atatagtagt aacggttgca caatatcgtg
8041 aaatgtacat agtgccactg aactgtatgc tttaaatgac ttaaatggta aatttagttt
8101 tttaaccaca atatttacca caattttta aatcagtgat gttgagattc agggaaatac
8161 ggtaataata gaagaaatac ggtaataaat agccaaagtg aaggagcacg gccaagttgc
8221 agaccatccc agacgccctt accctcctcg cctgccatgg ccccaactag tgccccagtt
8281 ccccctcccc ccacacccca cggtccaccg gggatccagg gccccagcgt gcaagtgcac
8341 gcccaggtgc ccgggctcct ggagcccctc cctatctagt ccaagaacgc cccgggtctg
8401 acaccttctc ttcggttctc cgcaggggac gtcgtcgtgc aggcgcccac ccaggtgccc
8461 ggcttcttgg gcgactccgt gacgctgccc tgctacctac aggtgcccaa catggaggtg
8521 acgcatgtgt cacagctgac ttgggcgcgg catggtgaat ctggcagcat ggccgtcttc
8581 caccaaacgc agggccccag ctattcggag tccaaacggc tggaattcgt ggcagccaga
8641 ctgggcgcgg agctgcggaa tgcctcgctg aggatgttcg ggttgcgcgt agaggatgaa
8701 ggcaactaca cctgcctgtt cgtcacgttc ccgcagggca gcaggagcgt ggatatctgg
8761 ctccgagtgc ttggtgagca gggggttttg gggaggctga atgaaaggca gagacttggt
8821 gggaggatca gggaagttgg caaagagcgg ggaggcctgg gagggaggga gattccctcc
8881 acagcagatc ccctggggac aaaaggaggg ggcagcgcaa tgatgtgggg tggggtgggg
8941 ggaggtgagt gggggagggg gctgcagggc aggaagaaag cagagatctc agggagtaag
9001 gacccccaag cctgggatca gaaagccctg agggaggagg aggggtatat tggaaaggga

9061 gctgggaggg gactgggaag gaagggagga ggcctgggtg gcctcttctg ggtccctcct
9121 ttcctcatct gtaaaagggt tggtgacaac agcatcaccc ttggaaagct gtcctgagtg
9181 tcatcactgc ctaggaaaga gtctgtctgc cccttggac tgtgtaccca tgagggcaga
9241 accggggcta ttttgatcac tgctgggccc cagcactttc caccaaacaa tatttattga
9301 gcacttatta tgtgccattc taagagcttt acctgcatat tctaattgcc tgtgattcca
9361 tcattatcac ctgttgacag acaaggaaac tgaggcacag acttgcccaa gaccaaagct
9421 gcagccttag ggcagcggcg cttcaccgct ggcccatgtg agccccttag agctgtcggt
9481 tactgttact ttatctggtg aagaaggaga aagttgaaac agacagacag gaaagcaagg
9541 atcctgatac ccaaaagagg aggggatcat gctttcattt tcatgagctc catgaagcct
9601 tcctcttgga acttgagcaa gaagtctctg tgcctcagtt tcctcatctg tgggataggg
9661 atcaccatga ggattcaaaa aaactgaccc tttttttttt tttgagatag tttagttctg
9721 tcgcccaggc tggagtgcag tggtgtgatc atagtaggtc ctcactgcag cctggaactc
9781 ctgggctcaa gggatgttcc taccctccct gtagctggaa ttacaggaat gtgccaccat
9841 gcccaacttg ttttctgttt tgttttgttt tgttttgttt tcttttcttt ttaagcttat
9901 gtgccaagaa ctattccaag cctgtgcatg agaaactcat gaaatgtcaa gggcttggaa
9961 ctgtgcctag cacacaacac ccaaccaaag gcagctgttg tcactccagc aagcctggga
10021 ttctgggtat catggtttag gatgagggta aaacagaaac agtgagtcaa gttaaggaga
10081 aatcttcggt aaataatata ggtaatgatg gggaaggaag ttccaggttt gggaatcacc
10141 aggatgatga ttaagagatg gtcaaggtgg ttttggagtc agacagactt tatgtgaaag
10201 cccagctctg ctgcttacag ctgtgtgacc atgggcaagt cacttccctt ctcaacttca
10261 gggctcaggc ttcctctcct ccactgagcc ccagaccagg ataacccct ttgggtttcc
10321 cctttccagc cctgccctct ccgagtcatc actgcctagg aaagagtccg tctgcccccт
10381 tggactgtgt acccatgagg gcagaaccag ggctattttg gtcactgctg ggctccagca
10441 ctttccacca aacaatattt attgagcgct tattatatac cattctaaga gctttacctg
10501 catattctca ttgcctgtga tcccatcact atcccctgtc aacagacaag gaaactgagg
10561 cacagacttg cccaagacca cacagctgct cagggacaca gctgggattt aaaccatgcc
10621 attctgtatc tttttttttt tttttttttt gagacagaat ctcactctgt cgcccaggct
10681 ggagagcagt gacacaatct cagctcactg caacctccgc ctcccgagtt caagcaattc
10741 tctggtctca gcctcccaag taactgggat tactggcgtg tgccaccata ccctggctaa
10801 tttttaatag agacagtttc tccatgttgg ccaggctggt ctcgaactcc tgaccccaag
10861 tgatccaccc acctcagccc cccaaagttc tgggattata ggcgtgagcc accccgccca
10921 gcaaccatgc catcctgtac ccttaatgaa tgccccсttc tgccacggag gggttcattg
**10981** aatgacttgt tgcttttgtt cctcttccca g**c**caagcccc agaacacagc tgaggttcag
11041 aaggtccagc tcactggaga gccagtgccc atggcccgct gcgtctccac agggggtcgc
11101 ccgccagccc aaatcacctg gcactcagac ctgggcggga tgcccaatac gagccaggtg
11161 ccagggttcc tgtctggcac agtcactgtc accagcctct ggatattggt gccctcaagc
11221 caggtggacg gcaagaatgt gacct**gcaag gtggagcacg agagc**tttga gaagcctcag
**11281** ctgctgactg tgaacctcac cgtgtac**t**gt gagtgtgccc aagtcagcga tggcaagaac
11341 ccctgccggg ctgcccccac cactgtctac actgactccc caaggcactg taggcattgc
11401 ttccatcatc tgcaccggct cccctgactc ctgtctacac gacccacccc cattgtctgc
11461 accagctccc ctgggccgta aacaacactg tccctattgt ctacaccagt ttacttgggc
11521 catagtcaat gctggtaccc agactgtctg ccctggctcc tctagcggct gtctgcacgg
11581 atacctgaca ctctgttgac tcccttgagc acagtctaca ctgcccctaa actgcctaca
11641 cccatttccc caggcaagta tccacaatga ccctccccac attgctacac tggttcttct
11701 ggccacgatc aacactgcct cccttattgt ctacacccac tcccttgtcc ttttctttct
11761 ttcctttttt tttttttttt tttgaaatag agtctcactc tgttacccag gctgcagtac
11821 agtggcatgt tctcggctca ctgtaacctc tgcctcccaa attcaggtgg ttctcctacc
11881 tcagcctccc aaatagctgg gattaaaggt gcacaccacc acacctggct aatttttgta
11941 tttttagtag agacagggtt tcatcatgtt ggccaggctg gtttcgaact cctgacctca
12001 ggtgatcggc ctgcctcagc ctcccaaagt atgctggaat tataggcatg agccactgtg
12061 cccggctttt tgttttgttt tgtttttcct ttggacaggg tctcgctctg tcacccaggc
12121 tggagtgtag gggcatgatc atggttccta cagcctcacc cttccaggct caagcaatcc
12181 ttctccctca gcctcccaag taactgggac tacaggtgca gccaccatgc ccggctaatt
12241 ttacagtttt tttctagaga gagttttgct gtgttgccca ggctggtctc aaactcctag
12301 gcttaagtga tcctcccgcc tcagtctccc gaagtgctga gattacaggc gtgagccact
12361 gcacctggcc ccttgtccat tttctacact gccttaccat tgtctacact ggctctcctg
12421 ggccacagac agcactaacc ccattgtcta caccacctat tttggccacc agctgtgggg
12481 tgttttcact gtctcccata gtgtctacac tggctcctat gaccactgtc tatgtaaaca
12541 ccccactttt tctacactgg ttctcctagg cactctctaa agtgcccccт gacactgtct
12601 acactgactc cctggagaac tgtctgcatt cacctcccac attgtatcga ccagttcccc
12661 tgagccacga tcaactgtct ccgccgggtg ctagttgcat tgatcttctc acattgtttg
12721 tgctggctcc cctaagcagt gtctgcagag tcccccagtg tagtctccat gttcctctgc
12781 ccaatattgt ctacactggc tcatctacag gctatccagt ctcaccccac atcaccacac
12841 tggcctcctt gtgttctgtc tgcacagccc cctgacagca tctgtcctga ttctcctgca
12901 cagcgttacc tccaccagct ccctcatggc atcatcgaca ttgacaccca cacatttatc
12961 tccactgaca gccccaggac tctctaacca gtgctgtcta acagaaatag catgcaagct
13021 gcatatagga ttttaaattc tccagcggcc accccattaa aaaattaaaa gaaagaggcc
13081 aggcacggtg gctcacgcct gtaatcccag cactttggga ggccgaggca ggtggatcac
13141 ctgaggtcag gcattcaaga ccagcctgat caatgtggtg aaactctgtc ttcactaaaa
13201 aatacaaaaa ttagccaggc acggtggcgg gtgcctgtaa tcccagctac tcgggagact
13261 gaggcaggag aatcacttga acctgggagg cggagattgc aatgagcgag attacgccat
13321 tgcactccag cctgggcagc agagcaaaac tccatctcaa aaacaagaaa agaaagagat
13381 gaaatttatt tgtttattta tttatctatt tatttatttg agacggggtc ttactctgtc
13441 acccagactg gagtgcagtg gcacagctcc ctgcagcctc aacctcctgg gcttaagtga
13501 tcctcccacc tcagcctcag tctcctgagt agctgggact acaggcgtgt gacaccatgc
13561 ctagctaatt tttgtttttg tttttgtttt tgtttttttg ttttgtgatg aagtcttgct
13621 ctgtcaccca ggctagagtg cagtggcacg atctcaactc actgcagcct ccgcctctcg
13681 ggttcaagtg attctcctgc ctcagcctcc cgagtagctg ggattacaag ccctggccac
13741 agtgcccagc taatttttgt atttgtggta gagacgggat ttcaccatgt tggccaggct
13801 ggtctcaaac tcctgacctc gtgatccacc caccttggcc tcccaaattg ctgggattac

```
13861 agacgtgagc caccgcaccc ggcctatttt tattttttct ggagacagga tctccctatg
13921 ttgcccaggc tggtctcaaa ctcctgggct caagtgatcc tcctgcctca acctcccaag
13981 gtactgggat tacaggcatg agccactgtg cctggccaaa attaattttt aataattgcc
14041 tttacttaac ccaatatacc taaaatatta tttcaacttg taattgtttt tttaaaatta
14101 ttcttgagat ctttcccatt tattttgtcc tagtaaagcc tttgaaatct ggtgtgtatt
14161 tcggcctggc tatatttcga gtgctcagta gtcacatgtg gccagtaggt gccatattgg
14221 acagcacaga tctagactaa gctcccacat catctacact ggctcatcca aatactgtct
14281 acacacatac acagagcctg tcttaatttc gtcagctccc ggggacacac tgtatatgtg
14341 ggggtgatct cacccccacc accacatttt caacactggc ccgacaatat agcatctgct
14401 gtaatattcc aacaccctgt ctacactgaa cccatcagta atatccacac aagctcttct
14461 agacatcatt catactgttc ctcccaccat tgtctatact gcttactccc cgccaggcaa
14521 ttggtacaca gattccctca ggtactgtct atactggagt ccctcaacac tgtcactggg
14581 tctgagcgag gaggagatct acactgactc ctacccctcg cccccgaata aaaccctgtc
14641 ccattggcct gcccccaagg acattgaaag gtcttcgctg tcaatcacgt atgggatggc
14701 tgagcagcag cctagagcag ggttatctgg gactctttcc tgcaggactt gtcagaacct
14761 tgaatacact gctgtgtgtt gtgagtggtc attgcctatg agtgagaaaa ggaggaggca
14821 ttttcctgca ggacacccaa gacatgcctg gtccaggcta cccaggaaag ctagtggaga
14881 aaagcagctc ctcagaagcc ttctccatgt ccccactgac caaggactcc taggcttctg
14941 ctctctgtgg caccccatct acctagactc ctcctaggcc tctccagctc ccgctgtttt
15001 ccaaatatcc ccgggccttt tcctgcagtg tcgtgaatcc cgcgtagccc caggcccccc
15061 aaagcctccc agtctctgaa cctctgtatc catttcctgc agaccccccca gaggtatcca
15121 tctctggcta tgataacaac tggtaccttg gccagaatga ggccaccctg acctgcgatg
15181 ctcgcagcaa cccagagccc acaggctata attggagcac gtgagtcctg ggtctcaggg
15241 aggaggggct gggggtctgg atttctagca ctgagggagg aggggctggg ggcctggact
15301 cctgggactg agggagaagg ggctggggc ccggacccct gggtctgagg gacgagaggc
15361 tgggagcccg gactcctggg tctgagggaa gaggggctgg gggtctggac ccctgggact
15421 gagagaggag gggctgggcc tggacccctg ggtctgaggg aggaggggct ggggggtctg
15481 gacccctggg tctgagggag gagaagctgg gagcctggac ccctgggtct gaggcaggac
15541 agactggggc ctggactcct ggtctaagag aggagggaga ttcagaaaag agaaaaagag
15601 gaggttgact tcgggccccc agtgggggc atctctgttt tgacatctct gttttgcggt
15661 tgaggcgttt agcaggtggt tgaatcctgg ctggtggatg cacctgcttt ttggggtgtg
15721 cctgctctgt gcccggtact tcccccgaat cccatgtgac cccatgctgt gcacagtcag
15781 cagctgttgc cctgcccagg ttaaagacca gccacgagag ggcagaagcg gcccttgaat
15841 cagcgctcgg ctccaaagct ggtcctccgc ctcttggcca agcttgtcta acctgcaggc
15901 cacatcattt gtaaactttc ttaaaacatt atgagacttt ttttttgcga attttttag
15961 ctcattagct attgttagtg ttagtgtatt ttatgtatgg cccaggacag ttcttcttct
16021 tccaatgtgg cccggggaag ccaaaagatt ggacaccccc actctagact gtaaccacct
16081 caccgtcacc ttgtgtaacc tacttggtta tctctggatg gagcaagtac tatatatccc
16141 cattgtctag atgagaaagt tgaggctcac agcaataaga tttccccaga ttcacactga
16201 ggctgagatc aagagcaagg taagaaaaag gcaaagagat aatacataat tttataatac
16261 ataattttac ataaattaca taattttact gaaccgccct tttttttttaa agaaggggct
16321 ctgttgccga ggctggagtg cagtggcgtg atcatagctc actgcagcct ggacctccct
16381 gggctcaggt gatcctccca cctcagcctc ctgagtagct gggactacag gcacacgcca
16441 ccacactcag ctaattttgt tgtatttttt ttgcagagac agggttttgc cacattgccc
16501 aggctggtct gcaactcctg ggctcaagca atcaacccac ctcagcctcc taaagtgctg
16561 ggattacagg cgtgagccac ctcgcccagc cttaggtctt tttccttatt acaaaaataa
16621 cttgtgtcca ctgtagaaat atcaaacaat ttagttatga gcaaaaagaa gaaagtagtg
16681 cctatagttc aagtgtaccc agccacctcc agaaataccc agtgacccct tattatacat
16741 cctgtctttt ttctctctgt ttctaaacaa agactggaat tatgctgtac atataatttt
16801 gaacctttc gtctgtcttg tgaataccttt tccatgtcaa agcacataga tccacaactc
16861 cattttcccc cacgttgtat taggaagact gtcaaacata cagaaaaatc ataggaattt
16921 tccagtgaac acttaagatt ccaccacgtg gattcccctg ctaacatttt actgtgcttg
16981 ctttatcacg tttctgtgat tccttcagtc cctctattca tccatcaatc atattatgca
17041 tttcagagca cccccagctc tcctgctttt tttttttttt tttttttttt gaggcagggc
17101 ctcactctgt tgctcaggct gtagtgccgt ggcacaatca cagctcactg cagccttgaa
17161 ttcctgggct caagcgatcc tccagcttca gcctcccaga tagctgggac tacaggcaca
17221 caccaccacg cctggctaac ctggcccctt ttttagtgtt gctgggattc caccacttta
17281 aaaaaaaaat tcaaagaata gagacaggat ctcactatgt tgcccaggct ggtcttgaac
17341 tcctgtactc aagcagtcat ctcacctggg cctcccaaag ggcagggaag ggattccaac
17401 atatgtggat gtcatcagtt atttaaccag tccctagtag aggacacaga tgtcattttc
17461 cattttcaac tctgcaaatg ccatggtggt gtccatcccc actggtcact tgtcagcagt
17521 tgtccgtgga aggaattgcc ggtcaaaggt tttcaactgg tgtgatggtg cctttactgt
17581 cctgcctcaa agcaccgcaa ttcaggctct tgagccagat agctgcgttt gcatcccagc
17641 tctgcggagc acccaaagcc tgatgcttcc atctcctcca cctctgttca ttaaaccagt
17701 gttccttgac tacttactgt gtgccgggtg caattaacca acatacgttg gtcgataaaa
17761 cagaccaaat ttcctcccca tatagagctt ccattctagt ggtgaaagaa acaagaagca
17821 agataaagtg tgtagtgtgt gtgcaatggt gataaatacc atggagaaga caaagcagag
17881 aagggaatgg gactgggcgc catgactcat acctgtaatc ccagcacttt gggaggcaga
17941 gatgggagga tcacttgagg ctaggagttc aagaccagcc tggccaacat ggcgaaaccc
18001 catctctacc aaaaaagata caaaaattgg ccgggtgtgg tggcatgcac ctgtagtccc
18061 agctactcag gaggctgagg tgggaagatt ccatgcaccc aggaggtgga ggttgcagtg
18121 agtcaagatg gtgccactgc actccagcct gggtgacaga gtgagacgct gtctcaaaaa
18181 aataataatt aaataaatta aagagaaggg gatgggaagg gtgatcatca cagttgaaat
18241 agggtggcca agtgaggtgt tactaaggtg acatcccagc agaggcctga aggagatgag
18301 ggtgagctct gcgggtgcct agggggaaag cattccagca gagggaacag catgggcaaa
18361 cccatttcta tccatcagtc cctgcataag caaacaccct ggacacctaa ctggagagtt
18421 tgaggaccaa caaagaggcc actgtggatg cagaggcggg agcaaagggg agagtaatgg
18481 aaaacgtggg cagaggtagc aggggccaga tcacgtgggg cctgtgacac cagaagactt
18541 cggtttaact ctgagccagg ctaggccccc agagggtact gagctgaggt atgggaactg
18601 acttagggtt cacaggcgac ctctggtggc catgtgggga acagacgagg gggcgagggc
```

```
18661 agaaacggag cccagcgtgg aggcggctgc tgggggccgg gatgcacagc gccagagcag
18721 ggaggtaccg tggaagccct gaggagaggt gggattctgg atggatctta aggcggggcc
18781 gacatgattg gctaatgggt tgggtgtggg catgagagga agggaagagt tggggggcct
18841 ccagagttgc tctccgctta tgctatagag ggcgtgaggg tttctgggca cgtagtgcgt
18901 gctcaatcac tgtcactccg gacctgcagc agaggccacc tcctcacctt tctgtctctc
18961 ccaggaccat gggtcccctg ccaccctttg ctgtggccca gggcgcccag ctcctgatcc
19021 gtcctgtgga caaaccaatc aacacaactt taatctgcaa cgtcaccaat gccctaggag
19081 ctcgccaggc agaactgacc gtccaggtca aaggtgagga actccctggg tgggaagaac
19141 agggaccaaa ggaagaggtc gaggcagggt tccctgtcta atgctctctg gctcccatcc
19201 ttctcctaag ccttctcaca aaaggaccag tggggccagc tttactcccc tgttaactcc
19261 atacctatat cactggtacc gttgttcagc tgcctggata cactccctcc acacacacag
19321 gttgggatag atgtggaatc cctctctctc agggactcag gcctttccaa atatattagg
19381 cagacctcta tcaattgcaa ttgacagaaa cacaaagtag tctaagcaaa aaaaaccaaa
19441 caaaacagga ctttattgaa aattcacctt caggtatggt tagatccagg ggctccaacc
19501 ttgtaaccca ggaactttta ttttatacat ctcttgtctc tgcttttcgt tatactggct
19561 ttattctcag gcaggctctg aagcaagatg gccctcggca gggacaggct cacattctac
19621 cagaatggca acagcaggag aaaaaacata ccactttcag taactgtagc catagtccca
19681 gggctggctc tcattggccc agcttaggcc acagcccacc tatgaaccaa tcatagcagc
19741 cagggagaat ggaatattct gattggccag gcctgtgtca aagtcccatt ctgggcacca
19801 gagggtggga tccattccac ctgaattaaa tagactgagg cagtggagtg gtggctccct
19861 aaggaaattc aagatgccat ttcaagaatg agggataaac cccaagagtt tccttaccta
19921 aatacctgtt tccttctctt tcagagggac ctcccagtga gcactcaggc atgtcccgta
19981 acgccatcat cttcctggtt ctgggaatcc tggtttttct gatcctgctg gggatcggga
20041 tttatttcta ttggtccaaa tgttcccgtg aggtcctttg gcactgtcat ctgtgtccct
20101 cgagtgagca tcaccagagc tgccgtaatt gagcacctac tacgggctct gtgctgagtc
20161 cttccagtgt gcctctcact gaatcctcac cccactgcca tgaggtttcc cccatttgac
20221 tgatgagggt gcagagccag ggagccttgt tcactggttc attgattaca tttacaaata
20281 ttatttacag agtgggagaa gagcgtatag ggtcttaatg ccatggtagg gactttggaa
20341 tttaattcag gtgatgtggg agtccctgcc agatggatgg agggtgaaat aatgcttaag
20401 ccctcagcct tactccatct gccttgttca ggagtgactc ctaggaggag gtggcaggaa
20461 ctggcacctc tgatcctcag ctctaaaagg gtctgatgct ttcacacctc aagagatttg
20521 cacaggccgt tcccctgcct ggagcaccct cccatggctc cttctggaga agtgtgccag
20581 ctcattctgt aatcagagga acgatgcttt gctgagcact gtctacatgc cactttgtat
20641 gtataacccc ctcgatttta atttttaatt tttttatta tttatgtatt tatttattta
20701 tttttgagac aggctctcac tgtcgcccag gctggagtgc agtggtgcga tctcggctca
20761 ctgtagcctt cacctcccag gttcaagcga ttctcatgcc tcagcctccc aaggagcggg
20821 gattacaggc acgcactact atgcccagca atttttgtat ttttaggaga gacagcgttt
20881 cgccatgttg gccaggctgg tctcaaaccc ctggtctcct gtgatccacc cgcctcagcc
20941 tcccaaagtg ctgggattac aggcatgagc caccacacct ggcctgtaca atctccttga
21001 atgcccacaa ccacccagca aagtggttct cccgtgagcc ccatgttaca gaggaggaag
21061 tatttgcttt gagagatgaa gttacctgcc aatggtcaaa cagctggaca tggttgttgg
21121 ggagaagaag ccagatcttt gactaccaag cttttccttc aaccactaaa caaagctgct
21181 cccttacaac ccaggcggat actgcctcgt ccaggaagcc ctccctgacc cccaagcatg
21241 agccaagcat ccactaggct ctcccactcc agccttgccc attctgggtc atcaccgtgg
21301 gggacttgtt tgtcatcccc agtggactgt gagccccaag atagcagacc tgaggctgtc
21361 actgtcatga ctgtgtcccc aggaccgcct accactgggc cgggcataaa agaggcactc
21421 gaggaagtgt ttgctgagtg attcaatgtg tcagctccag agccagactg cctgtatctg
21481 ggccctaaag ctactgcttg attagtgatg tcaccatagg caaatagctt aagctgtatg
21541 actcagtttc ctcatctgta aaatggggtg atggtaccta ataatacagt agagcactta
21601 acacagtagg tggcacatgg taagccacat actgagtatg tatttgctat gattgttatt
21661 gctattatta ctattattgt tattattaac ataaactgca caagagcagg ctttttgttt
21721 gtttgtttgt gttttgtttt gagacaagat cttgctctgt tgcccagact ggagtagtgc
21781 agtgatgtga tcttggctca ctgcagcctc aacctcccag gctcaggtga ttctcccacc
21841 tcagcctctc gagtagctgg gattataggc gtgcactccc acgcctggct aatttttat
21901 atttttagta gagatggagt ttcgttgtgt tgcccaagct ggtctcaaac tcctagactc
21961 cagcaatccg cccacctcag cctcccagag tgctggaatt ataggtgtga gccaccacgc
22021 ctggccaaga gcaggcattt ttatctgttt tgttcacagc tgtgtcccca gcacctagaa
22081 tgtacctggc acaaagtaga cattcaccat ttgctgaata atactactat ttgctaaata
22141 aacccctgtt gaatagtcta ggaaactgaa cacagagagg ttaagtcact tgtccaagtt
22201 cacacagctc ctaaaggaca gagatggaat ttgaacccat ccatctggtt ccagagccag
22261 gctcgatatt tatggatatt tcataaccaa gtgaataact ggatggctaa atccttagaa
22321 agcacaccct ccatccctgt ctgacacacg tgtataccat taagggaaat ggcaccccca
22381 tgtaatagca gaggcagaac ttgacatttt cagggccctg acacccaggc atgtcttctt
22441 cccatgcttc ccagcattat tccttcctga cgacttcccc tcctatttcc ccaggtacag
22501 agcatgccag cgcctcagct aatggggtaa gtgcagtgtt ggagctaagg agggtgtggg
22561 gtctgcacga actacagacc agacgtcttc agatgcccac ggcccggcct caggagcctg
22621 ggtcttttct cctggtgcct cgagcagcat ttcccaaccc ttcctgctgg gcggaatccc
22681 ctgtagagtt tgctaagacg cagccccaga cagtctggtc tgtgatgggg ccagggaatc
22741 tgcattttaa tatccgctct ctcccccctg ccccacgccc cttcaattcc acccatggcc
22801 acaccttgag aagcactgcc ctggcttcca gggagagagg aggggacggg cctgcagttc
22861 tttgcacatg gggcttcagt gggaagatgt ttgattttgt tcaaaagagc cccaaggcta
22921 aaatttgaaa accctcttct agcatgtctc ctattcagct gtgagcagag agaacagctc
22981 ttcccaggat ccacagacag agggcacaag gtgacagcgt cgggactgag aggggagaga
23041 gactggagct ggcaaggacg tgggcctcca gagttggacc cgaccccaat ggatgaagac
23101 cccctccaaa gagaccagcc tccctccctg tgccagacct caaaacgacg gggcaggtg
23161 caagttcata ggtctccaag accaccctcc tttcatttgc tagaaggact cactagactc
23221 aggaaagctg ttaggctcac agttacagtt tattacagta aaaggacaga gattaagatc
23281 agcaaaggga ggaggtgcac agcacacgtt ccacgacaga tgaggcgacg gcttccatct
23341 gccctctccc agtggagcca tataggcagc acctgattct cacagcaaca tgtgacaaca
23401 tgcaagaagt actgccaata ctgccaacca gagcagctca ctcgagatct ttgtgtccag
```

23461 agttttttgt ttgtcttgag acagggtctg gctct**gttgg** **cagactagag** **tacag**tggtg
23521 agatcacagt tcattgcagc cttgacttct caacgccaag tcatcctccc acctcagcct
23581 cctgagtagc tatgactaca ggtatgtgcc accacgtctg gctaatcttt ttattatttg
23641 taaagtcgag gtttccctgt gttgcccagg ctggtcttga actcttggct ccaagtgata
23701 cttctgcctt ggcctcccaa agtgctgaat taagcagctc accatccaca cggctgacct
23761 catacatcaa gccaataccg tgtggcccaa gacccccacc ataaatcaca tcattagcat
23821 gaaccaccca gagtggccca agactccaag atcagctacc aggcaggata ttccaagggc
23881 ttagagatga atgcccagga gctgaggata aagggcccga tctttctttg ggcaaggtta
23941 agcctttact gcatagcaga ccacacagaa gggtgtgggc caccagagaa ttttggtaaa
24001 aatttggcct ctggccttga gcttctaaat ctctgtatcc gtcagatctc tgtggttaca
24061 agaaacagcc actgaccctg gtcaccagag gctgcaattc aggccgcaag cagctgcctg
24121 gggggtgtcc aaggagcaga gaaaactact agatgtgaac ttgaagaagg ttgtcagctg
24181 cagccacttt ctgccagcat ctgcagccac tttctgccag catctgcagc cagcaagctg
24241 ggactggcag gaaataaccc acaaaagaag caaatgcaat ttccaacaca agggggaagg
24301 gatgcagggg gaggcagcgc tgcagttgct caggacacgc tcctatagga ccaagatgga
24361 tgcgacccaa gacccaggag gcccagctgc tcagtgcaac tgacaagtta aaaaggtcta
24421 tgatcttgag ggcagacagc agaattcctc ttataaagaa aactgtttgg gaaaatacgt
24481 tgagggagag aagaccttgg gccaagatgc taaatgggaa tgcaaagctt gagctgctct
24541 gcaagagaaa ataagcagga cagaggattt gctctggaca gagatggaag agccgggaac
24601 agagaagtgt ggggaagaga taggaaccag caggatggca ggggcaaagg gctcaagggt
24661 gaggaggcca gtgggacccc acagagttgg ggagataaag gaacattggt tgctttggtg
24721 gcacgtaagc tccttgtctg tctccagcac ccagaatctc attaaagctt atttattgta
24781 cctccagcgg ctgtgtgcaa tggggtcttt tgtggaaatc aaggagcaga caggtttcat
24841 gtgtactgtc accacgtggg atggaaccag aggcatggaa gcaagacgct aaatgaagag
24901 ggccataagg gctgggattc ccaggcacct taggaacagc ttgtcttttt tttttcctc
24961 tccaaaaaaa atgtttaagg gacggtgtct cctgtcaccc aggctggagt gcaatggcac
25021 gatcatagct cattgcagcc tctaactccg ggctcaagc aatcctccca cctcagccta
25081 ccaagtagct gtgaccacag ctgcccctca ccatgctaag ctaatttttt taattagata
25141 gtacataaac gtcccaaaat tagaagataa aaagacatga gggatccatt ctaatttgtg
25201 tttggagtgt aatggtccag ctccattctt ctgcacatgg atatccagtt ttacacaaca
25261 ctgtgaatgt aatgaatgcc actgaatcat acactcaaaa atagctaaaa tggcaaattg
25321 tctgttatct ctttttaacc accatttttg aaaattaatt ataccaaaaa accattgaat
25381 agtgcacttt atttatttat ttatttgttt atttatttat ttattttaga aataagagtc
25441 tcactttgtt gcccaggctg gagtgcagtg gcgtgatcat ggctcattgc agcctcgacc
25501 tgctgggctc gggctatcct tccatctcag cctcccgagt agctgggact ataggtgggc
25561 gccaccccac ctggctaaat ctctttttaa cttttgtaga gataggcatc tcgctatgtt
25621 gcctaggctg ggctggaact cctgggctca agtgctcctc ctgccttggc ctcccaaagc
25681 gctaggatta cagatgtgag ccaccgcgcc caccctgaac cttacttttt ttgctcagtt
25741 tctggtaatt cagagaatgc ctcctgagtt gttctacacc cacctcatat tccatgggag
25801 ggctgtacag ggctttttta acgaggcctc taaggacagg catttgtatc ctttccagcc
25861 tttcactatt acaatgttgt agtgaataac tttacacact gtcatttatt ttactttttt
25921 tttttttat tttagagaaa ggaatcttgc catcttgccc aggctggtct caaattcctg
25981 ggcccaaaca atcctcccgc cttggcctcc taaagtactg ggatttatag gcataagcca
26041 ccgtgcctgg ccaatgcaca ctgtcattta gctcatgtta acacctgagt gtaggacaca
26101 ctcctggagg tggaattgct gggccaaaga gtatgtttct tgtcattgtg atagatattg
26161 acaaatgaac cctcacagaa gttgtgctga gttctgttcc caccagcgac gtaggcgatg
26221 acctttttct ggagggaggg ggcatccttg gagtccacag agccaggaat ggagagtggg
26281 cccagaattt tggtataggt gttgtataaa cttatagtaa ggttaagaaa accgcaacta
26341 tccttatcag agacttggcg gggggcaggg tatgatggag atcataagga ggctaaaaca
26401 ctccacaccc tccctctgca ttgctcctgc acgggagtcg ggaatctttt caggttgata
26461 cgatctcacc ttgaggagct gtgaggtccc agaagcctct gggttgcaga ttgcttgggg
26521 tgaaaatgtc tgtgctactg aaatctaact ttttacaaaa aattacgggc tgggcgcagt
26581 ggctcacgcc tgtaatccca gcactttggg aggctgcagc gggtggatca cttgaggtaa
26641 ggagttcaag accagaccat agtgaaaccg tgtctctaca aaaaaaatta gccaggtgtg
26701 gtggtgcatg cttgtaatcc cagctactca gaaggctgag gtgggagaat cccttgaacc
26761 cgggaagtgg aggctggagt aaaccatgat cgagttactg cactccagcc tgggtgacaa
26821 gagtgagact ctgtctccaa aaaaaaaaaa aaaaaaaaaa aaactggatt gcctggctct
26881 actccgggca cagcatgcag gcccagttct gctgctctgc tgtttgttct gctttcctcc
26941 acatattggc atcaccctct ggtgccaaga tggctgctgc attccaggca tcacatccag
27001 actcagaccc agagaagctg cccatcccta cctgggtgag cctttgtagg aacgagaaac
27061 cgcatccagc agcagaaacc tcacccagca gcgtcttttc cggtctcatt caccagcgcc
27121 gcccaccgct caaccaatcc ctggccaaaa gaatgggacc gcctggaagg ctggaccaaa
27181 caggacctgc cctctggggc tggggagagg cccagatgaa ggctgcagga caggatggac
27241 tcctagacct ctgttaccag cagtgactac ctctgtctgg gtggttggaa catgtttgaa
27301 ttttattcta agtactgtct acaagttctg caataaacct tgactcttct tttaataatg
**27361** caaa**a**ggaat cgaagtgatt gtttgaaagg gagaggaaga aagagagagg gagggaggga
27421 agaatggagg gaggcaggga aggagacaga gagagtagaa tccagccacc ggaaaatcca
27481 gaatagctgg ctttgcttaa tccatgcctg gaaataactg ctgggtttgc aacaacttct
27541 ctcccggaga cagaccaagg aaactacaaa actgcaggaa ggattgaagg gccgggcaca
27601 gtggctcacg cctgtaatcc caacactttg ggaggccgag gtggattgga tcacctgaga
27661 tcaggagttc gagaccagcc tggccaacat ggtgaaacct cgtctctgct agaaatacaa
27721 aaataagccg ggcgtggtgg cgtgcgccta tagtcccagc tactcgggag gctgaggcaa
27781 gagaatcgct tgaacctggg aggtggaggt tgcagtgagc tgagatcgtg ccactgcact
27841 ccagcttggg cgacagagtg agaatctgtc aggtggattc tgagggtact ctgaccccac
27901 gaacccagca tgaaatcttc caggcattaa ggaacattga ggatgtggag gattggaaat
27961 ccctccaggg aaaatacact tctcttgagc agggtcacga ggggtggggg caatgtggga
28021 ggtggggcaa atccaaagac aaggtcaaca gcaataataa taataacaat gacaggcatt
28081 tgttgcttgt tggctgggcc tcttgcatac agttctccca acatccctgt gaggtggtga
28141 tgcttatccc cattctccag gggaggaggc tgaggcccag agaggtgaag tcgtgtgcca
28201 caaaataccc cagccatatg gtggtagaac caagttttca gccagactca ccagctcaca

```
    28261 gcagagctgc cctggactct acagccattc agtgtatagc tgggccactc catcattcaa
    28321 ggcagttcaa cgtcaagatt cctgatttgc ctcaacctgg ggcctctaga agataccatc
    28381 acccccaaca agctgaagac gtgtgtccgg aacactcaga cacacacaaa gtgctatgga
    28441 tacaccacac cccgacctag ctgtctcttc ctgatggacc tcactgggta gaaatgcctg
    28501 gcaccgcctg cctcagatcc ctaggtcagc tctcactccc caatagaagt ttctgagatg
    28561 atgatgaaaa cgttctccat ctgtgctgtc caatggagta gccactgatc acttgaggct
    28621 agtgagcact tgaaggtggc tagtttgagg aactgcattt tttaatgcat ttaattttag
    28681 ttactttgaa tatatataca catatataca tttatatata catattcata cacatacatt
    28741 ttacatatat tttatatgta tatgtaatat atatataaat tatatatgta tatattttta
    28801 tatataaata tatattatat atttataaat atataaatat ataatatata ttatatagat
    28861 atataaatat aaatatatat tatatatata aatatataat atatattata taaatatata
    28921 taaattatat atttatatat ataatatata aattatataa attatatata tttatatata
    28981 atatataaat atatataaat tatataattt atatattata tataaattat atattttata
    29041 tataatatat acaatatata taaattatat ataatatata aatatatata aattatatat
    29101 aatatataaa atatatataa attatatata atatataaaa tatatataaa ttatataaat
    29161 ctatataaat atatactata ttataaatat agtatatatt tatatagatt tatattattt
    29221 atatagtata tattatatat ttatatgttt atgtatatat aaatatataa tatataaata
    29281 tatttatata ttatatattt atataatatt ttatataata tgttatatat ttttatataa
    29341 tatattatat atatttatat

Gene  1 - 29360
Exon1 5113 - 5378
CDS   5300 - 23014
Exon2 8426 - 8773
Exon3 11012 - 11308
Exon4 15103 - 15220
Exon5 18965 - 19113
Exon6 19945 - 20103
Exon7 22495 - 22526
Exon8 22943 - 27365
polyA_site 27063 - > 28524
```
