## Supplementary figures and images for "Macrophages target PVR/CD155 on colorectal cancer cells via REVERBα"

### Supplement Figure S1

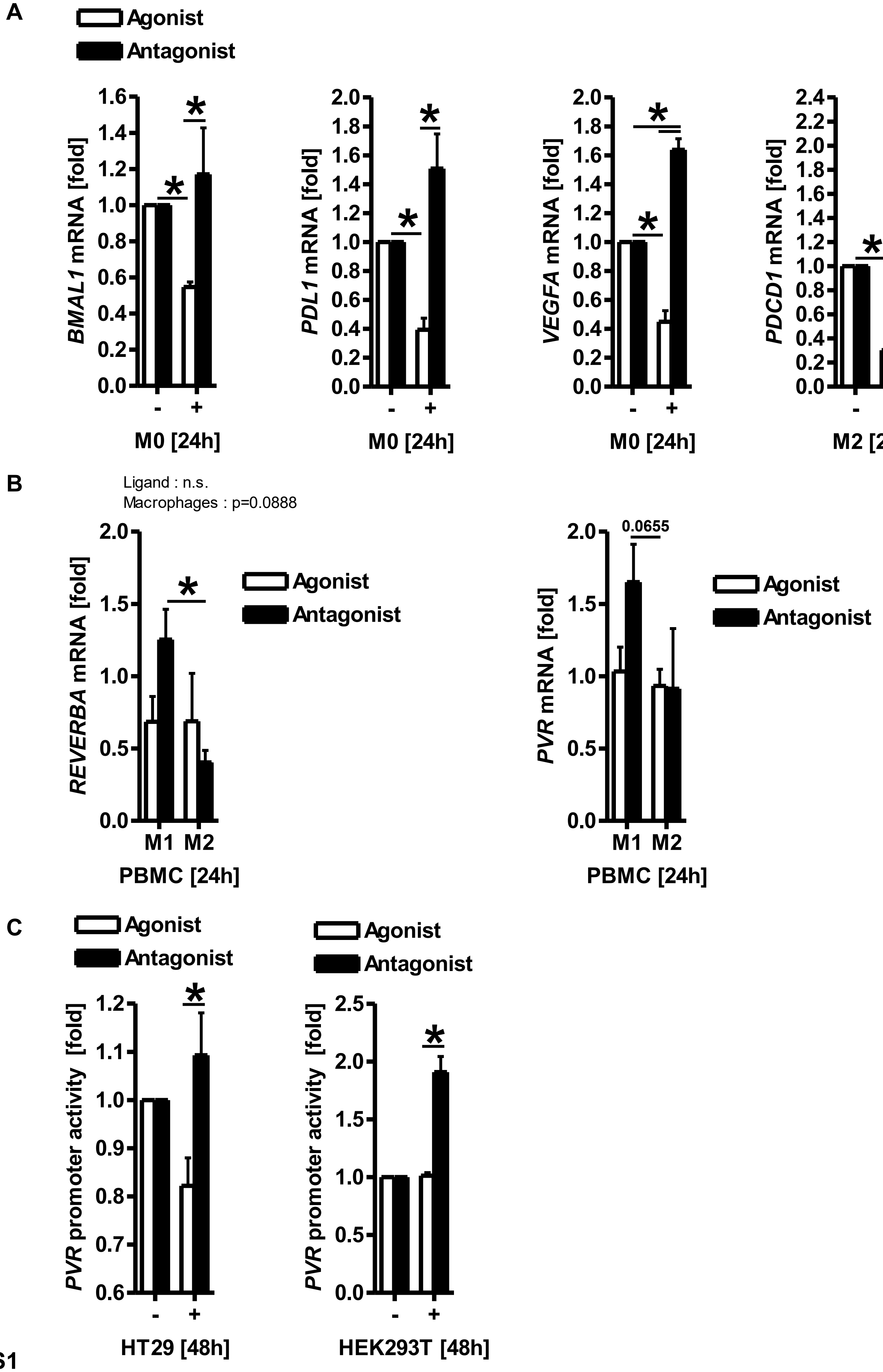

### Supplement Figure S2

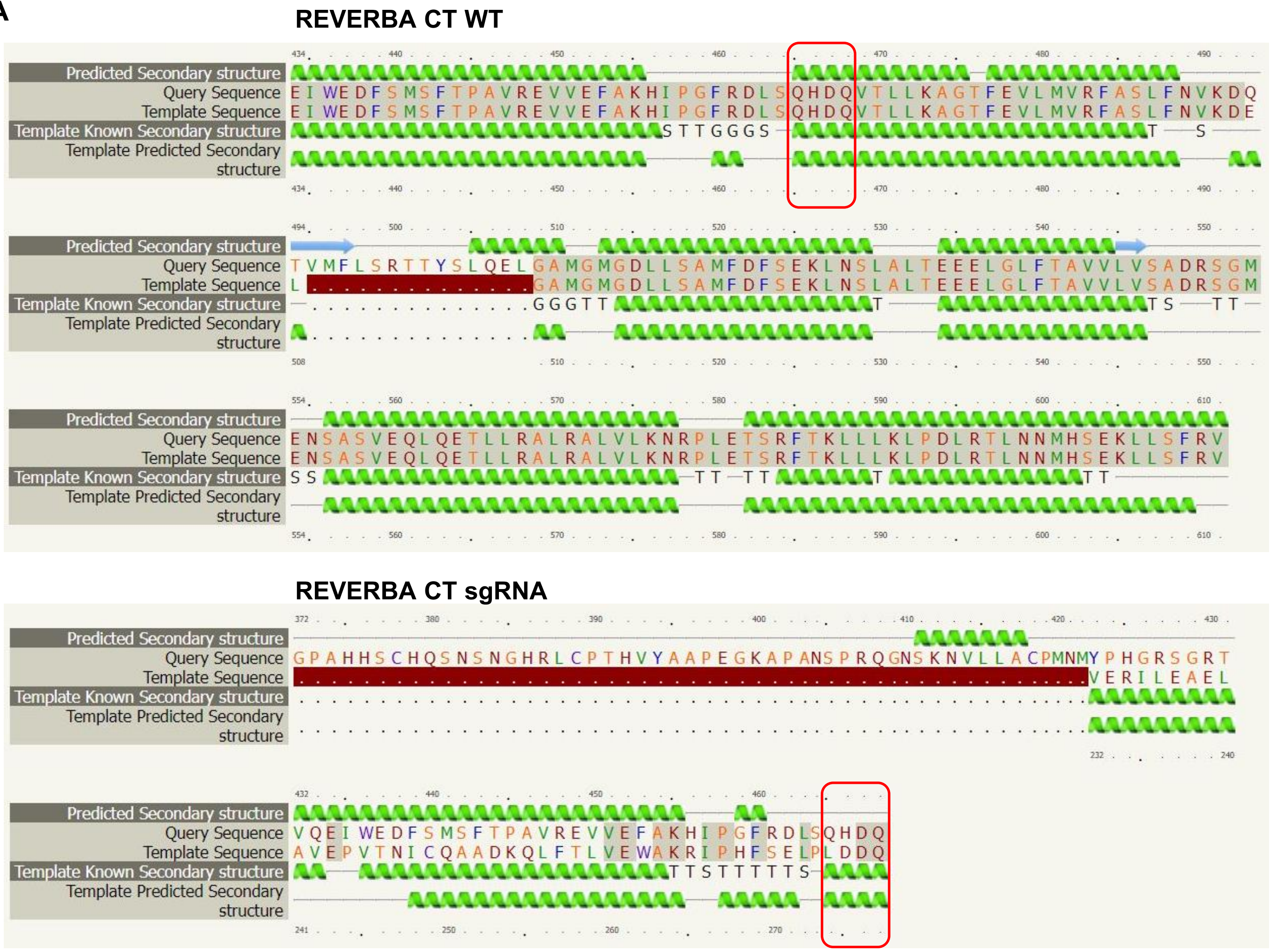

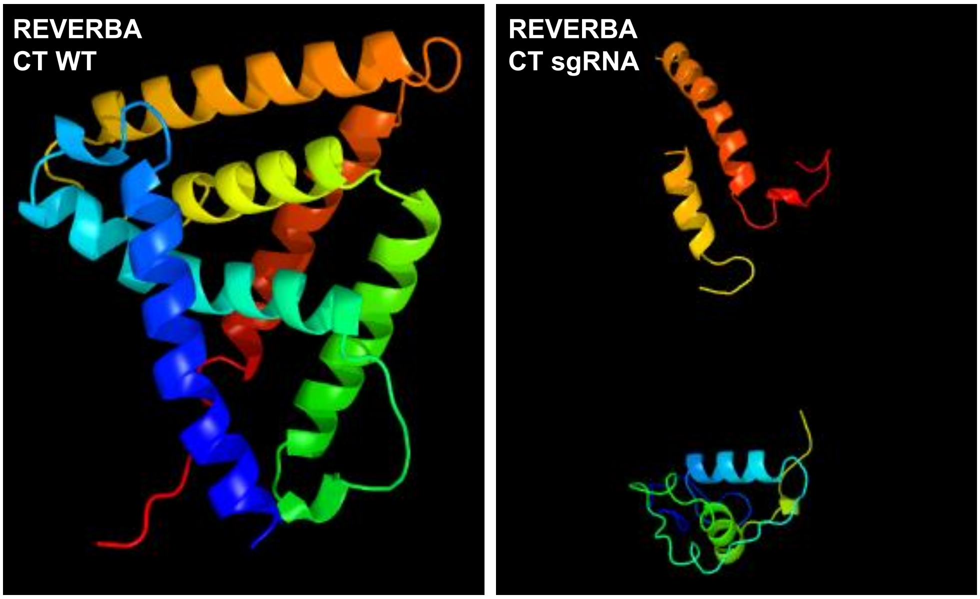

### Supplement Figure S3

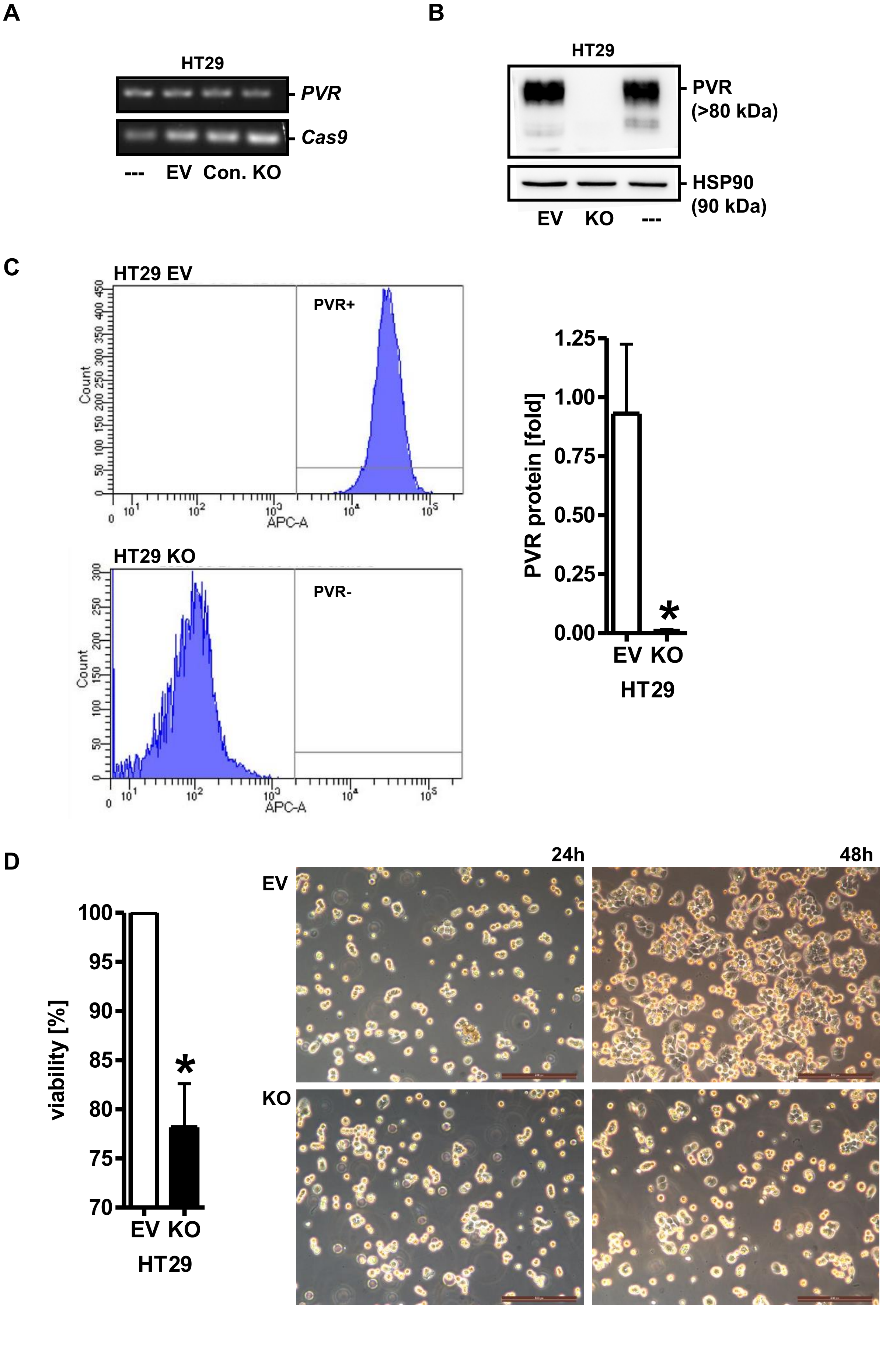

### Supplement Figure S4

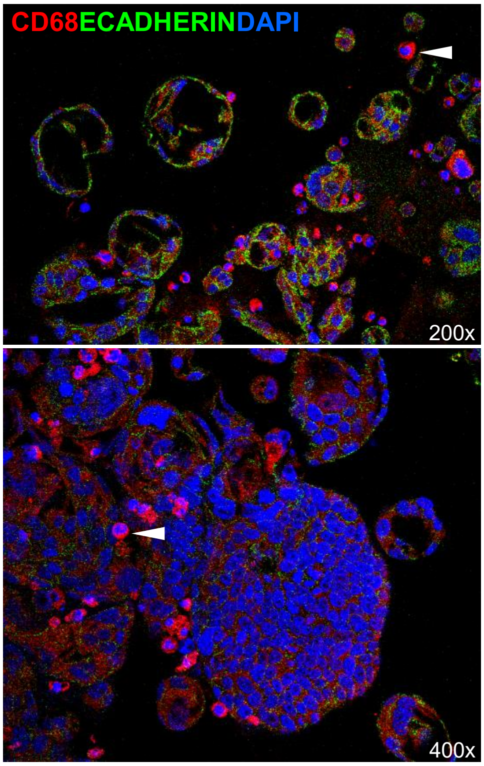

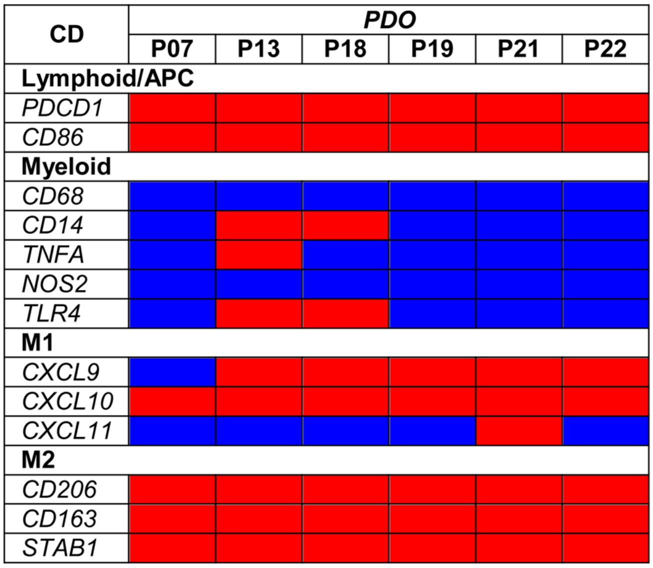

### Supplement Figure S5

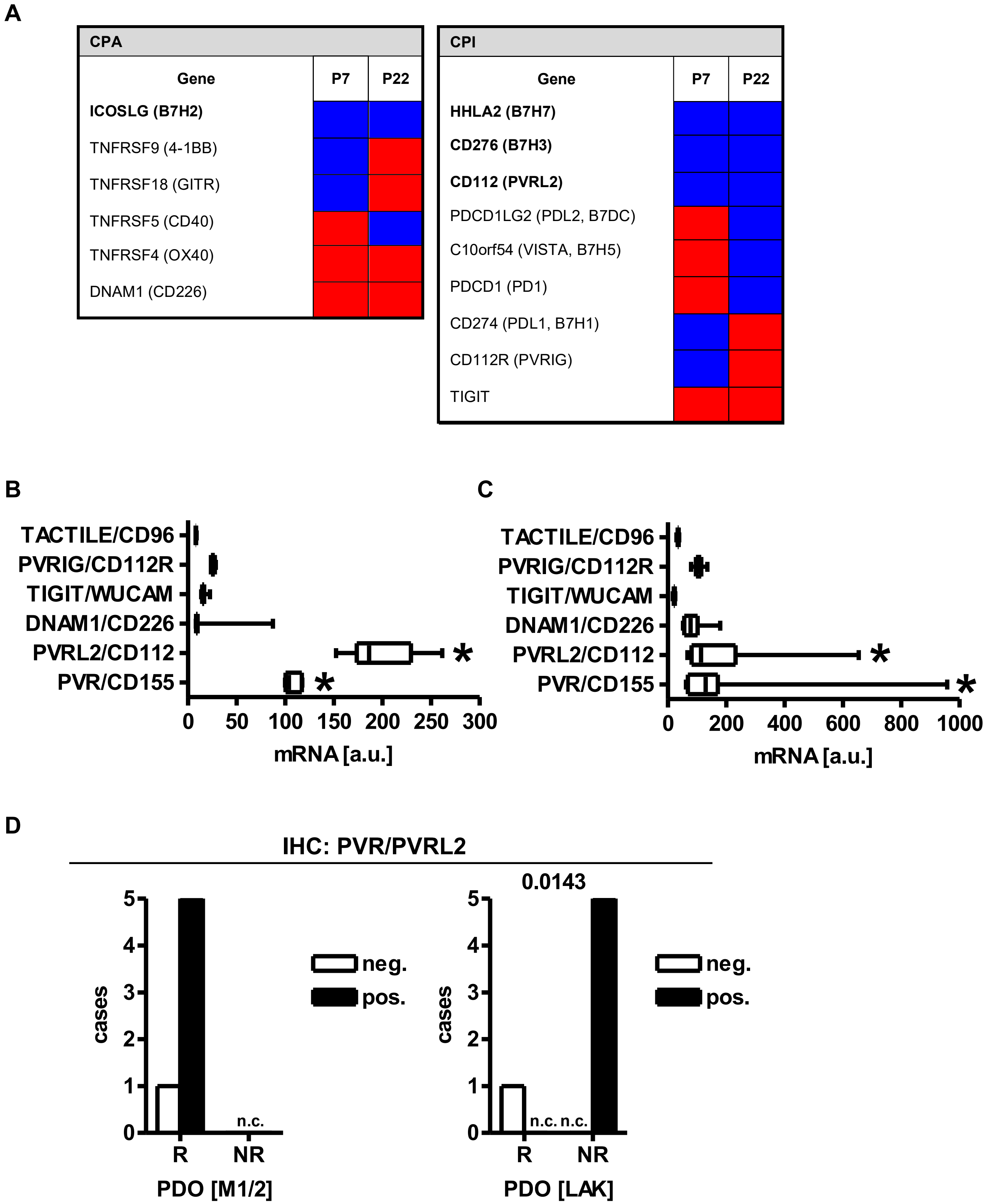

### Supplement Figure S6

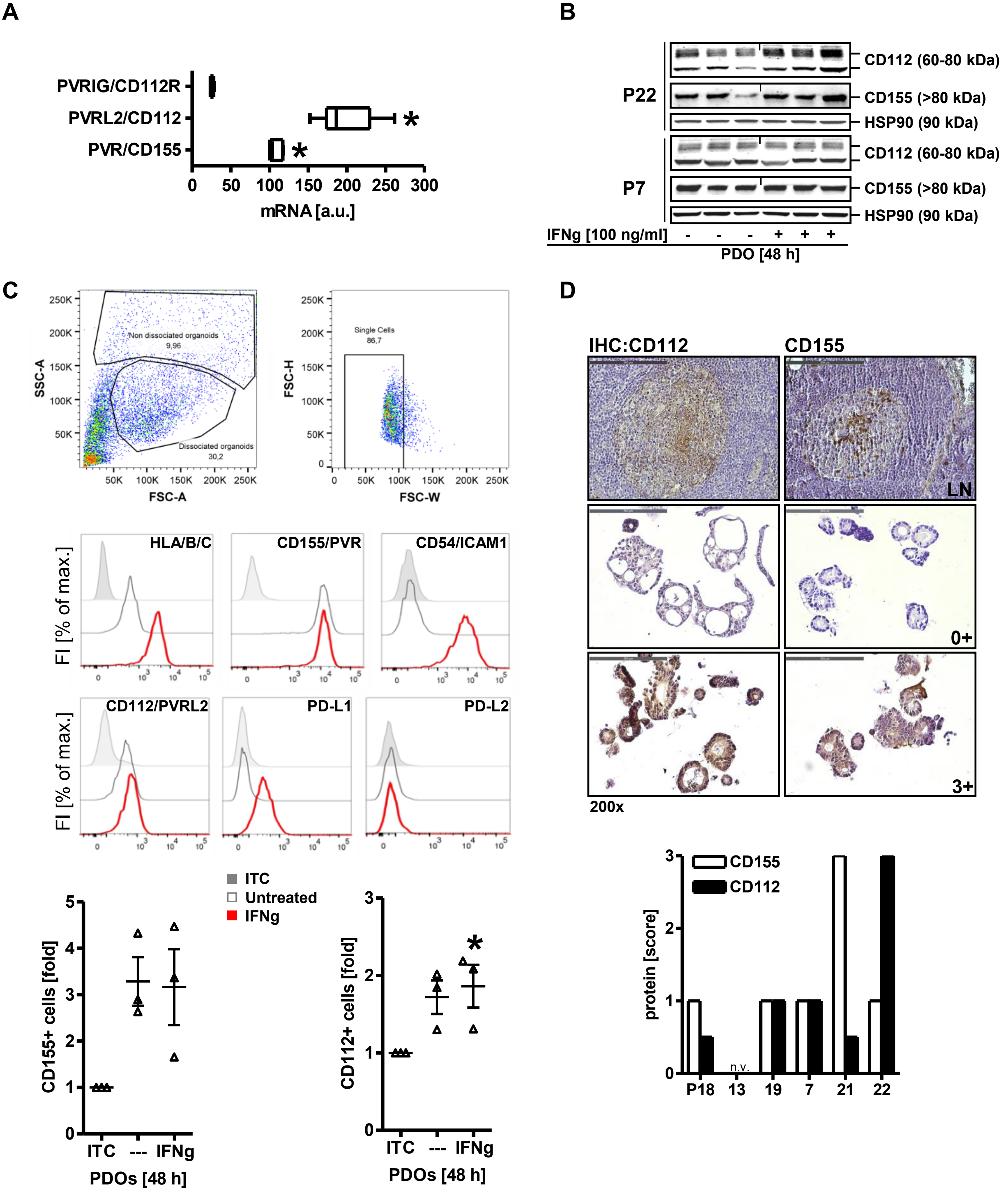

### Supplement Figure S7

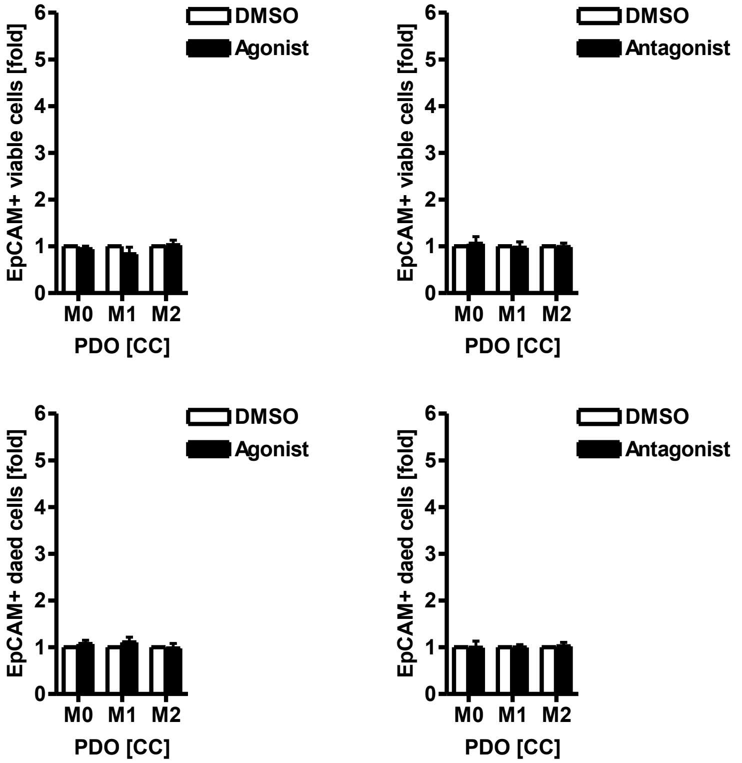

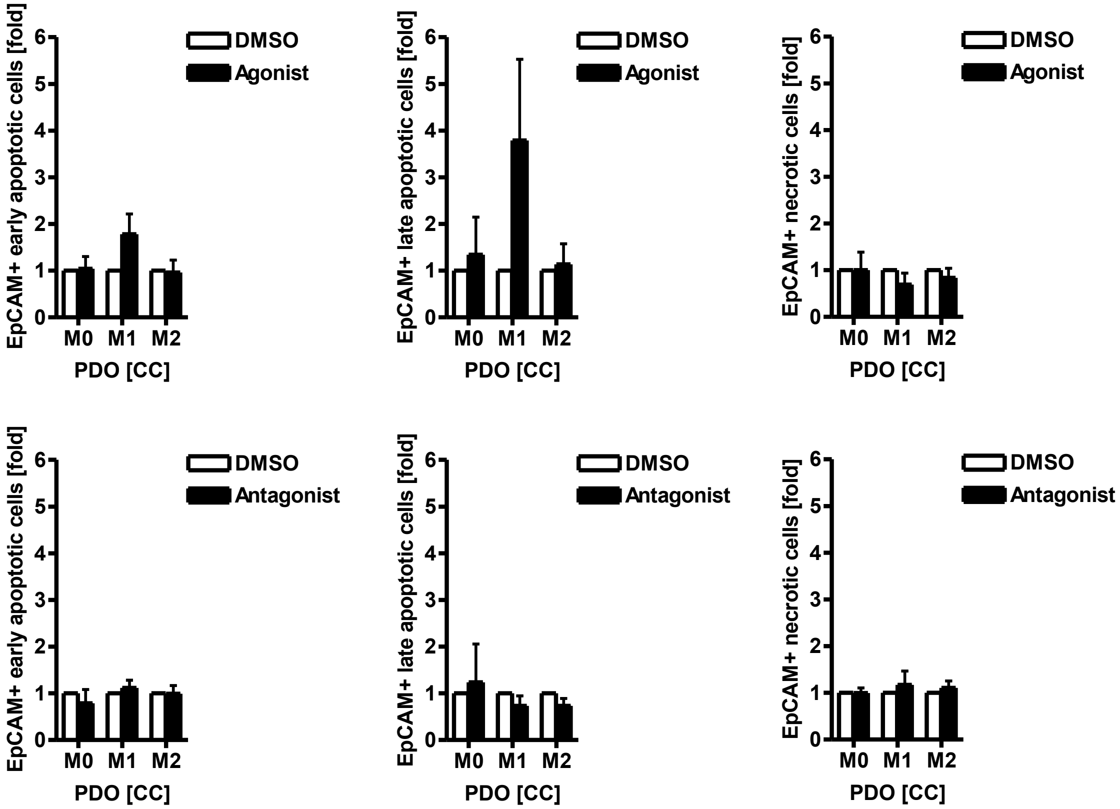

### Supplement Figure S8

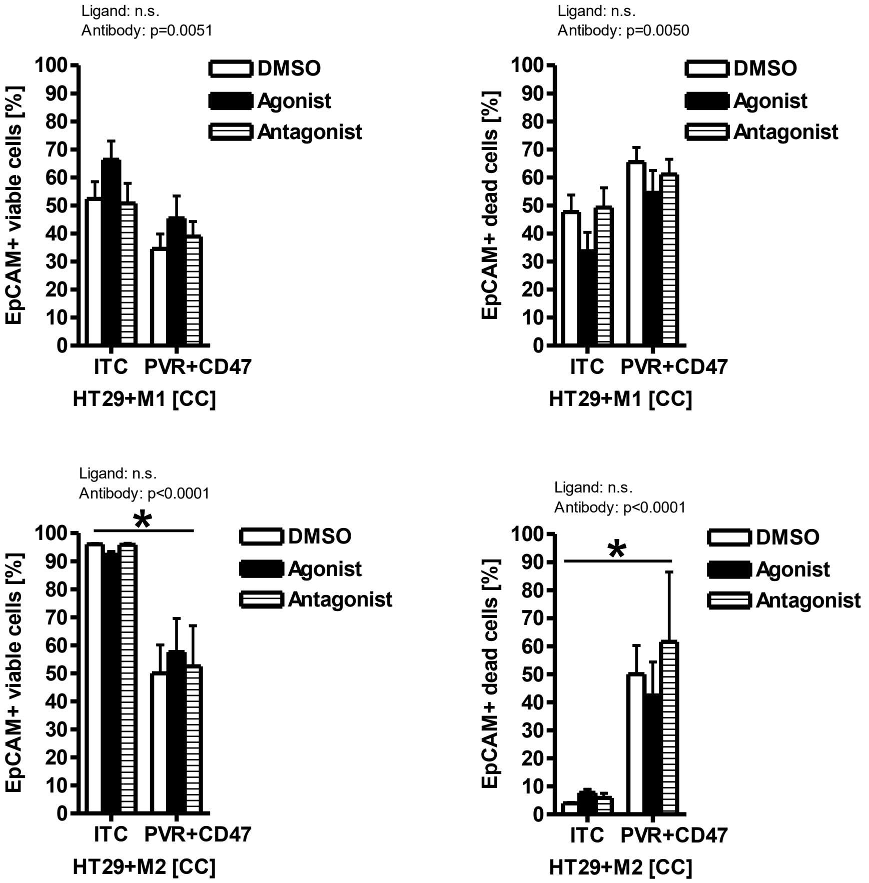

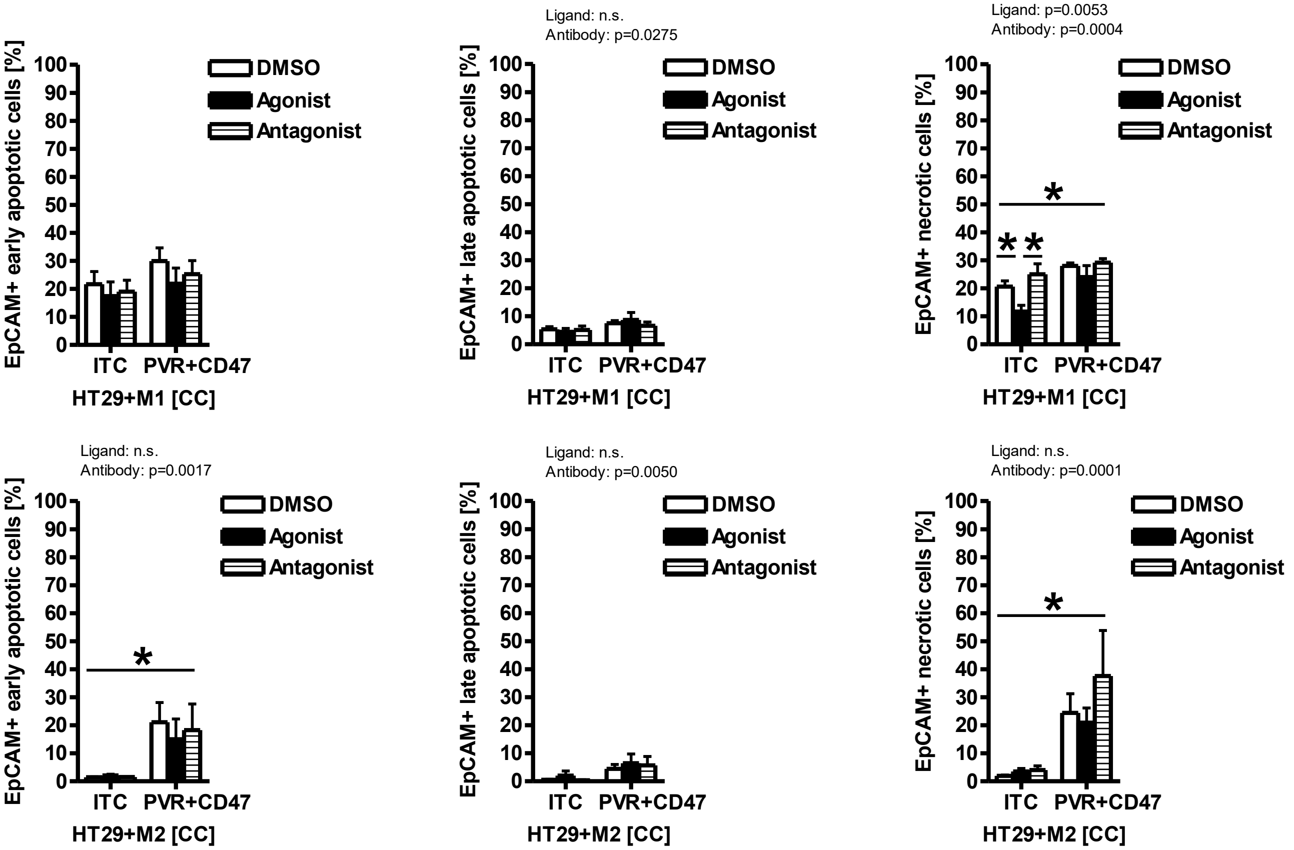

### Supplement Figure S9

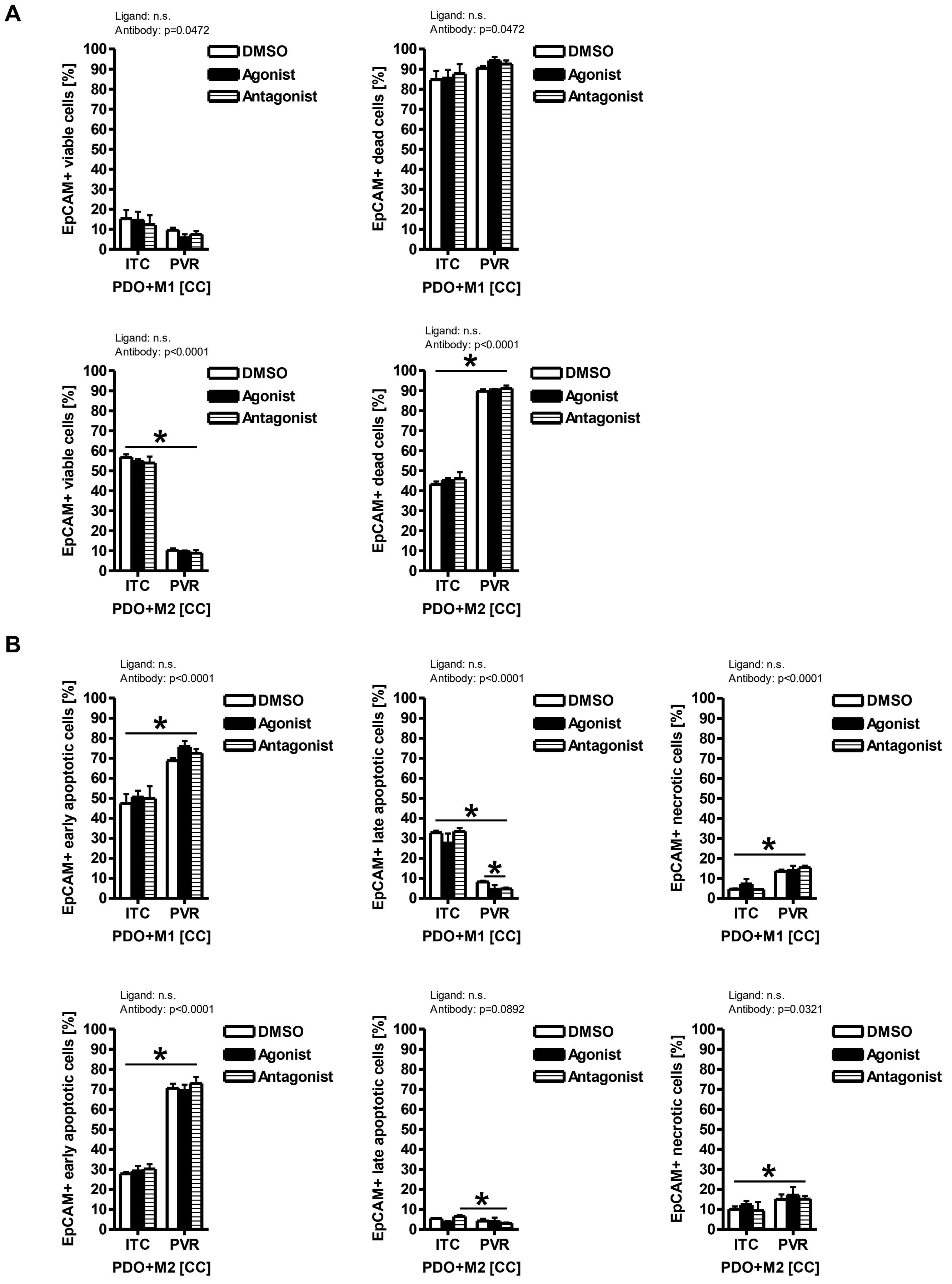

### Supplement Figure S10

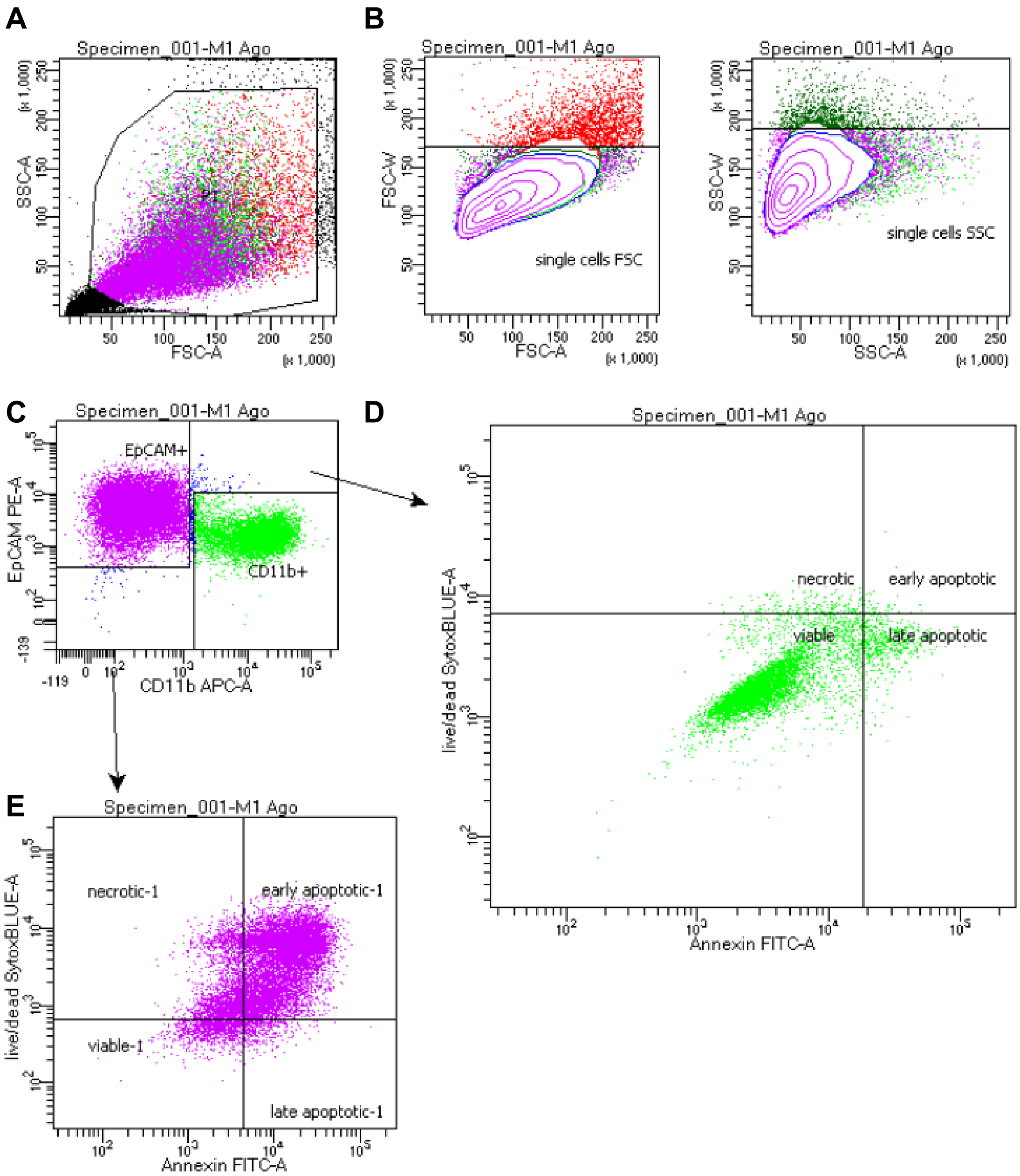
